# Identification of candidate biomarkers and pathways associated with multiple sclerosis using bioinformatics and next generation sequencing data analysis

**DOI:** 10.1101/2023.12.05.570305

**Authors:** Basavaraj Vastrad, Shivaling Pattanashetti, Chanabasayya Vastrad

**Affiliations:** Department of Pharmaceutical Chemistry, K.L.E. College of Pharmacy, Gadag, Karnataka 582101, India; Biostatistics and Bioinformatics, Chanabasava Nilaya, Bharthinagar, Dharwad 580001, Karnataka, India

**Author notes:** Chanabasayya Vastrad, Chanabasava Nilaya, Bharthinagar, Dharwad 580001, Karanataka, India.

**Keywords:** multiple sclerosis, differentially expressed genes, protein-protein interaction network, bioinformatics, hub genes

## Abstract

Multiple sclerosis (MS) is an autoinflammatory disease that might lead to severe disability. The diagnosis of MS is defined due to the urgency for biomarkers with both reliability and efficiency. Demyelination of axons are deeply involved in the pathogenesis of MS. Our study aims to identify the underlying molecular mechanism and screening for related biomarkers and signaling pathways. We obtained next generation sequencing (NGS) dataset (GSE138614) from the GEO database. Differentially expressed genes (DEGs) were screened by the DESeq2 package in R bioconductor with considering specific criteria. Gene Ontology (GO) enrichment analysis, REACTOME pathway enrichment analysis were performed; a protein-protein interaction (PPI) network was constructed; significant modules were analyzed and hub genes were identified by Human Integrated Protein-Protein Interaction rEference (HiPPIE). Subsequently, miRNA-hub gene regulatory network, TF-hub gene regulatory network and drug-hub gene interaction network were built by Cytoscape to predict the underlying microRNAs (miRNAs), transcription factors (TFs) and drugs associated with hub genes. Receiver operating characteristic (ROC) curves analysis was performed to calculate diagnostic value of hub genes. Finally, we performed molecular docking study for prediction of drug molecules against protein targets. A total of 959 DEGs (479 up-regulated and 480 down-regulated genes) were identified in the MS samples and compared with normal control samples. The DEGs were predominantly enriched in an ensemble of genes encoding the immune system process, developmental process, immune system and regulation of cholesterol biosynthesis by SREBP (SREBF). A PPI network was obtained through HiPPIE analysis, and the results were imported into Cytoscape software. The DEGs were sequenced by the Network Analyzer plug-in by various calculation methods, and 10 hub genes (LCK, PYHIN1, SLAMF1, DOK2, TAB2, CFTR, RHOB, LMNA, EGLN3 and ERBB3) were finally selected. Based on the miRNA-hub gene regulatory network and TF-hub gene regulatory network construction, miRNAs including hsa-mir-6794-3p, hsa-mir-3689a-3p, hsa-mir-4651, hsa-mir-548q, BRCA1, HNF4A, TFAP2C and NR2F1 were determined to be potential key biomarkers. Drug-hub gene interaction network constructed from DrugBank, which identified targeted therapeutic drugs (Palivizumab, Cu-Bicyclam, Lumacaftor and Zonisamide) for the hub genes. From molecular docking study we showed good drug - protein bind affinity and amino acid interactions. This study identified novel biomarkers for MS and established a reliable diagnostic model as well as predicted novel drug molecules. The transcriptional changes identified may help to reveal the pathogenesis and molecular mechanisms of MS.

## Introduction

Multiple sclerosis (MS) is the commonest autoimmune inflammatory disorder of CNS white matter [1]. MS is one of leading causes of disability in children’s, young aged adults and middle aged adults in many developed countries [2–4] and it affects 2.3 million people worldwide with an expected increase in the number of cases in future [5]. Main features of MS include selective primary demyelination with partial preservation of axons and reactive astrocytic gliosis [6]. It might affect many other organs and cause other MS associated complications include spasticity [7], paralysis [8], bladder, bowel and sexual dysfunction [9], psychiatric disorders [10], cardiovascular diseases [11], neurodegenerative diseases [12], spinal cord injury [13], obesity [14], diabetes mellitus [15] and hypertension [16]. Autoimmune inflammation is triggered by both genetic susceptibility [17] and environmental factors [18]. However, the etiology and symptoms of MS are complex in clinical practice, making its diagnosis challenging.

Correct early diagnosis assessment of MS is very difficult, even though much disease related genes and cellular pathways related to MS have appeared [19]. The common treatments of MS are interferons, glatiramer acetate, teriflunomide, sphingosine 1-phosphate receptor modulators, fumarates, cladribine and monoclonal antibodies [20], but there are no valid treatment tactics available to treat MS. Therefore, it is vitally important to explore potential diagnostic and prognostic biomarkers, and therapeutic targets of MS.

The fast development of next generation sequencing (NGS) technology and bioinformatics analysis based on high-throughput data provide new tactics to identify differentially expressed genes (DEGs) and discover therapeutic targets for the initiation and evolution of MS [21]. Altered expression of key genes plays an essential role in the initiation and progression of MS, so mastering the alteration in the characteristics of critical genes and signaling pathways promotes to comprehensively understanding of MS progression [22]. Several biomarkers including CLEC16A (C-type lectin domain containing 16A) [23], IL2RA (interleukin 2 receptor subunit alpha), RPL5 (ribosomal protein L5) and CD58 (CD58 molecule) [24], SLC9A9 (solute carrier family 9 member A9) [25], IL22RA2 (interleukin 22 receptor subunit alpha 2) [26] and ANKRD55 (ankyrin repeat domain 55) [27] were identified for the MS. Indeed, some researchers found signaling pathways include mTOR signaling pathway [28], Keap1/Nrf2/ARE signaling pathway [29], NF-κB signaling pathways [30], wnt signaling pathway [31] and Nogo-A/NgR signaling pathway [32] have been shown to be associated with MS. The identification of novel biomarkers and signaling pathway might be helpful to improve the clinical outcome of MS patients.

However, the use of bioinformatics analysis method to find the relevant genes and signaling pathways of MS has not yet been confirmed. In the current study, we downloaded NGS dataset GSE138614 [33] from the NCBI Gene Expression Omnibus (GEO) [https://www.ncbi.nlm.nih.gov/geo/] [34] database, including NGS data from 73 patients with MS brain white matter samples and 25 normal control brain white matter samples. We identified DEGs using the DESeq2 in R bioconductor tool. The GO term and REACTOME pathway enrichment analyses were performed via g:Profiler. Moreover, protein-protein interaction (PPI) network and modules, and the hub genes were identified according to toplogical properties include node degree, betweenness, stress and closeness scores. Subsequently, miRNA-hub gene regulatory network, TF-hub gene regulatory network and drug-hub gene interaction network were constructed and analyzed according to the screened hub genes. Subsequently, ten hub genes were subject to receiver operating characteristic curve (ROC) analysis to determine diagnostic value of hub genes. Finally, molecular docking study was performed for prediction of possible binding affinity of selected ligand molecules. This approach will enhance our understanding of the role of key genes and signaling pathways in MS and help to elucidate the specific molecular mechanisms of damage of white matter in brain. Additionally, the identification of these genes and signaling pathways might provide novel targets for intervention in the treatment of

## Methods and materials

### Next generation sequencing data source

GEO is a public functional genomics data repository of high throughput NGS data (RNA-Sequencing). The GSE138614 [33] NGS dataset generated using the GPL21697 NextSeq 550 (Homo sapiens). The GSE138614 dataset contained 73 MS brain white matter samples and 25 normal brain white matter control samples.

### Identification of DEGs

Determining DEGs between MS and normal control samples was performed using DESeq2 package [35] in R bioconductor tool.The adjusted P-values (adj. P) and Benjamini and Hochberg false discovery rate were applied to strike a balance between finding statistically important genes and limiting false positives [36]. |log FC (fold change)| > 2.2 for up regulated genes, |log FC (fold change)| < −0.866 for down regulated genes and adj. P-value < 0.05 were considered statistically significant. DEGs were visualized by the “volcano” and “heatmap” by using ggplot2 and gplot packages in R software.

### GO and pathway enrichment analyses of DEGs

Online tool g:Profiler (http://biit.cs.ut.ee/gprofiler/) [37] was used to perform the GO functional annotation (http://www.geneontology.org) [38] and REACTOME (https://reactome.org/) [39] pathway enrichment analyses. GO enrichment analysis is a premier bioinformatics analysis for identifying high-quality functional gene annotation based on biological process (BP), cellular component (CC) and molecular function (MF). The REACTOME is a resource of pathway database for the clarification of high-level features and effects of biological systems. Adj. P-value <0.05 was considered as the threshold.

### Construction of the PPI network and module analysis

Information of DEGs’ protein experimental interactions and prediction was obtained by Human Integrated Protein-Protein Interaction rEference (HiPPIE, http://cbdm-01.zdv.uni-mainz.de/~mschaefer/hippie/index.php) [40]. Subsequently, a specific PPI network of DEGs was constructed by cytoscape (version 3.10.1, http://www.cytoscape.org/) [41] based on the interactions retrieved from HiPPIE. To screen the hub genes that might be involved in MS, we applied the Network Analyzer plug-in, using various topological parameters such as node degree (refers to the number of connections (edges) a given protein (node) has within the network) [42], betweenness (measure that identifies the extent to which a node (protein) serves as a bridge or intermediary for communication between other nodes (proteins) in the network) [43], stress (network metric that measures how much a node (protein) is involved in “stress” or bottleneck situations within the network) [44] and closeness (measure that indicates how close a node (protein) is to all other nodes in the network) [45]. In addition, PEWCC analysis [46] in cytoscape was used to identify the significant modules of the PPI network

### Construction of the miRNA-hub gene regulatory network

miRNet database (https://www.mirnet.ca/) [47] is used to explore miRNA- hub gene interactions for the input hub genes from PPI network and assess the effect of the miRNA on the expression of the hub gene. The parameters used were as follows: specify organism: H. sapiens (human); set ID type: Official Gene Symbol; hub gene - miRNA interaction database: TarBase, miRTarBase, miRecords, miRanda (S mansoni only), miR2Disease, HMDD, PhenomiR, SM2miR, PharmacomiR, EpimiR, starBase, TransmiR, ADmiRE and TAM 2. Cytoscape (v. 3.10.1) [41] software was used to visualize the miRNA-hub gene regulatory network.

### Construction of the TF-hub gene regulatory network

NetworkAnalyst (https://www.networkanalyst.ca/) [48] is used to explore TF- hub gene interactions for the input hub genes from PPI network and assess the effect of the TF on the expression of the hub gene. The parameters used were as follows: specify organism: H. sapiens (human); set ID type: Official Gene Symbol; hub gene -TF interaction database: JASPER. Cytoscape (v. 3.10.1) [41] software was used to visualize the TF-hub gene regulatory network.

### Receiver operating characteristic curve (ROC) analysis

ROC curve analysis, which shows indicators of accuracy such as the area under the curve (AUC), provides the fundamental principle and rationale for distinguishing between the specificity and sensitivity of diagnostic performance of hub genes. The diagnostic utility of hub genes in MS was then assessed using a pROC package in R bioconductor tool [49]. Expressions of hub genes within the training and validating datasets were illustrated for assessment. When the area under the curve (AUC) exceeds 0.9, the hub gene will be regarded as good diagnostic value for MS.

### Molecular docking study

#### *Insilico* molecular docking studies

##### Software tools

Swiss-model, RCSBPDB, Prank web, ChemDraw, Avogadro tool, Autodock 1.7.1, and Autodock Vina tools, Biovia Discovery Studio client 2021.

#### Receptor and ligand preparation

The 3D crystal structure of the respective proteins was obtained from the respective genes by using the PDB IDs were downloaded from the RCSB Protein Data Bank (PDB) [50] (Table 1). The missing residues were remodelled by the Swiss-Model web server and downloaded in the pdb format [51]. Then the binding sites were identified by using the server Prank web [52], the pdb format of protein was inserted in the Software Auto Dock Tools 1.7.1, water molecules and hat atoms were removed. The receptor has also been screened for them issuing aminoacid residues, repaired missing residues, Kollman charges, and only Polar hydrogens were added. The grid was fixed using the Autogrid utilityin such a way that to covered the whole receptor, and later saved the grid dimension file [53].The structures of ligands were drawn by using ChemDraw (Fig.1) and copied as SMILES, followed by loading them into Avogadro software for converting them from 2D to 3D structures in pdb format, then pdb format of the ligand was inserted in Auto Dock Tools 1.7.1, then the root of the ligand molecule was detected and chosen. Finally, the ligand molecule was saved in the pdbqt format.

**Fig. 1.**
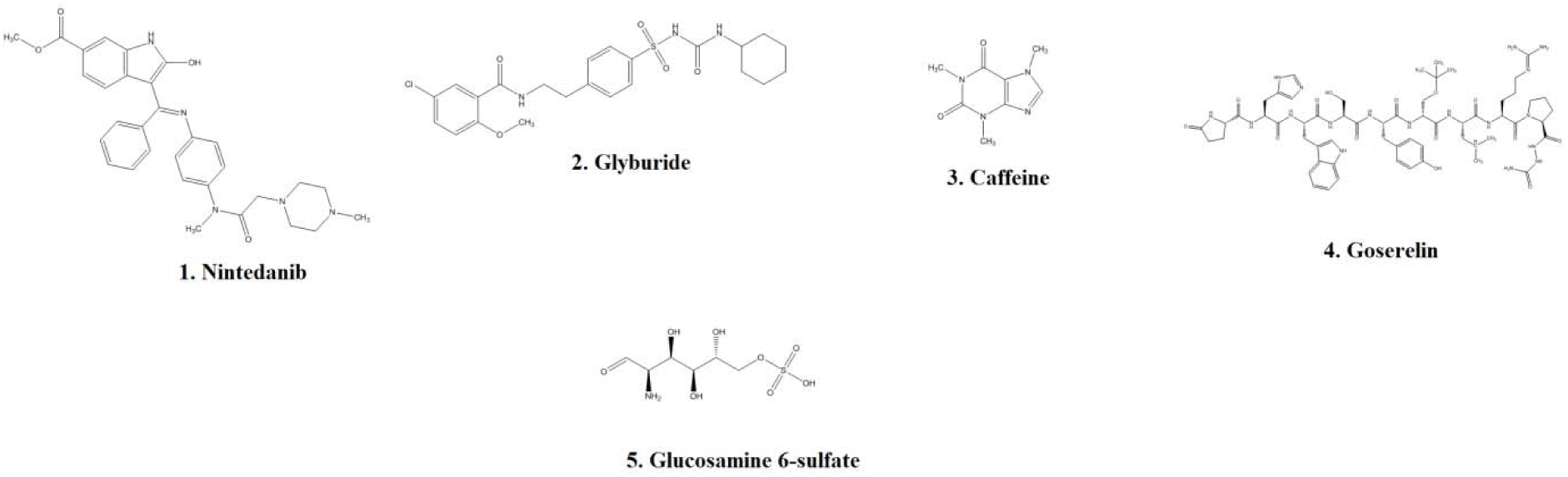
Chemical structures of drug molecules.

**Table 1.** Selected drug targets with their structural information.

| Drug target | PDB Id | Resolution (Å) | Method of collection |
| --- | --- | --- | --- |
| LCK | 2og8 | 2.30 | X-RAY diffraction |
| CFTR | 2pzg | 1.8 | X-RAY diffraction |
| CFTR | 2bbs | 2.05 | X-RAY diffraction |
| CA9 | 5dvx | 1.60 | X-RAY diffraction |
| LHCGR | 1qfw | 3.50 | X-RAY diffraction |
| CA14 | 4lu3 | 2.00 | X-RAY diffraction |
| HGF | 3hms | 1.70 | X-RAY diffraction |
| PRKCQ | 1xjd | 2.00 | X-RAY diffraction |

#### Performing AutodockVina

There are two ways to run Auto Dock Vina: from the Autodock tool or the command line (cmd). For running Autodock Vina, the configuration file was prepared, the grid dimension file saved previously consists of active site, x, y, and z coordinates, and n-pointsof the protein. That data was inserted into the config file; aconfig file was created which included the information of the active site of the protein, also it includes the log file in the form of .txt format and output file in the form of pdbqt. Autodock Vinawas run on the command line. The command “vina.exe -- config config.txt” was given to run Auto Dock Vina once the run was completed, docked coordinates were out in the pdbqt format, the file was used to visualize the receptor-ligand interactions, and the log.txt file was used to find binding affinity [53].

#### Visualization

The docking results have been visualized by using the Biovia Discovery Studio Client 2021software. The output pdbqt file and the receptor pdbqt file format were inserted into the Discovery Studio Client2011software. T he 3D image of the docked ligand in the PNG format, and the 2D image of the docked, which is bound to the different amino acids, are then saved in the form of PNG format [54].

## Results

### Identification of DEGs

Data from an overall 73 MS and 25 normal control samples from NGS datasets (GSE138614) was studied in a retrospective manner in the present research. The DEGs of the metadata were studied via the DESeq2 R bioconductor package. A threshold of adj. P-value <0.05, logFC (fold change) > 2.2 for up regulated genes and logFC (fold change) < −0.866 for down regulated genes were utilized to screen DEGs in the dataset of GSE138614. An overall 959 DEGs were identified: 479 genes were remarkably regulated upward and 480 genes were distinctly regulated downward (Table 2). In the volcano plot, every plot indicated a gene and the green plots were up regulated genes, the red plots were down regulated genes (Fig. 2). Then, heatmap (Fig. 3) identified that the expression difference of DEGs between MS samples and normal control samples was very decided.

**Fig. 2.**
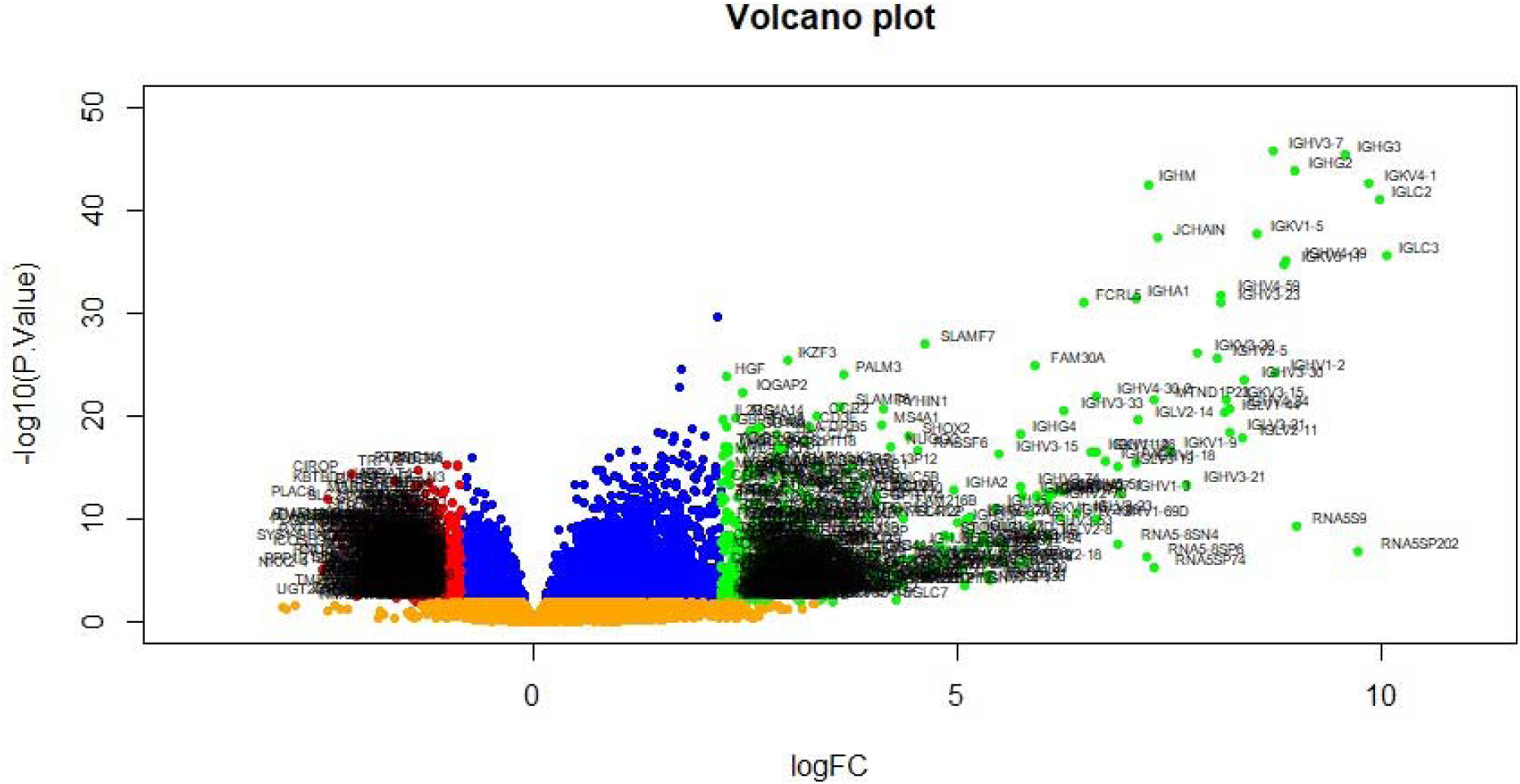
Volcano plot of differentially expressed genes. Genes with a significant change of more than two-fold were selected. Green dot represented up regulated significant genes and red dot represented down regulated significant genes.

**Fig. 3.**
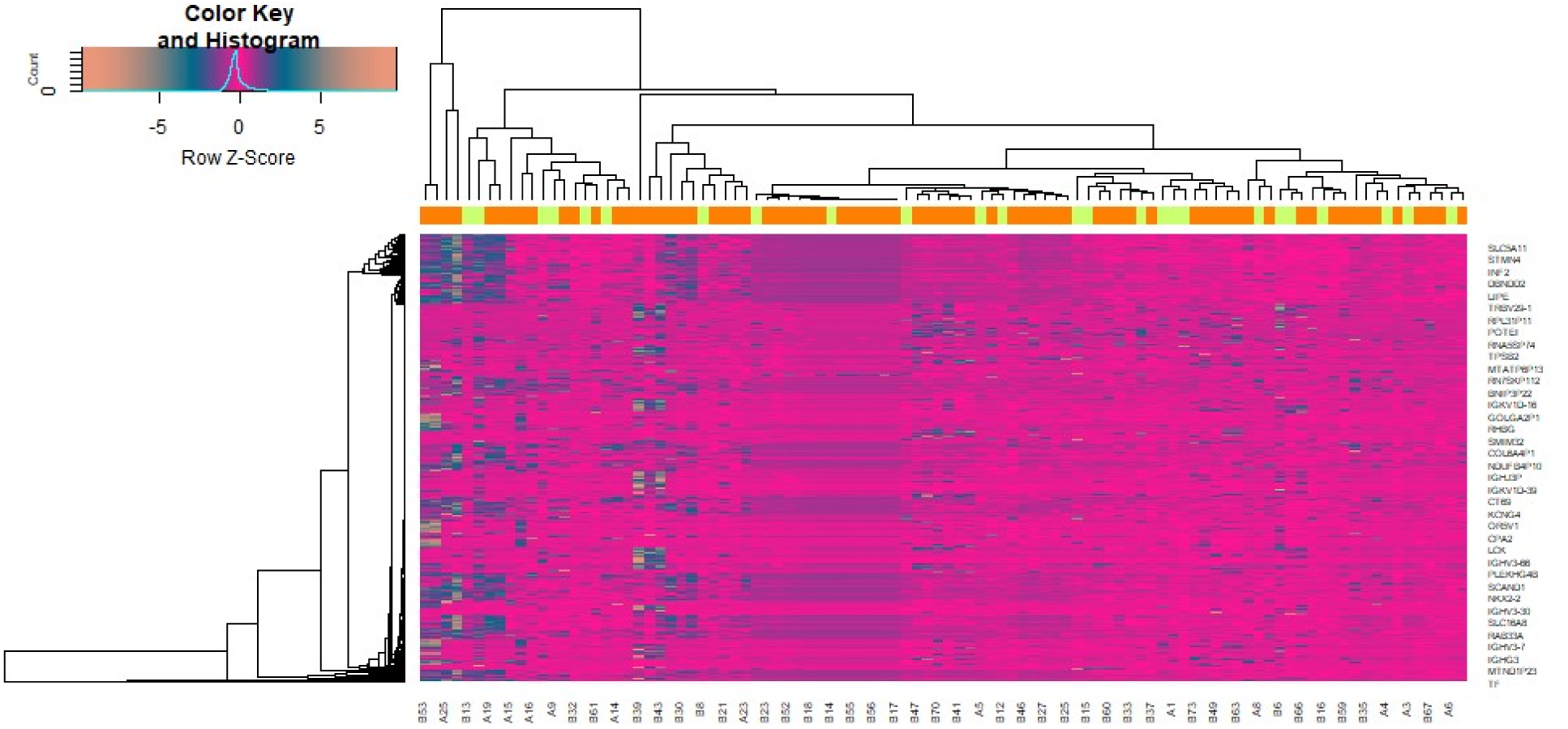
Heat map of differentially expressed genes. Legend on the top left indicate log fold change of genes. (A1 – A25 = Normal control samples; B1 – B 73 = MS samples)

**Table 2.**
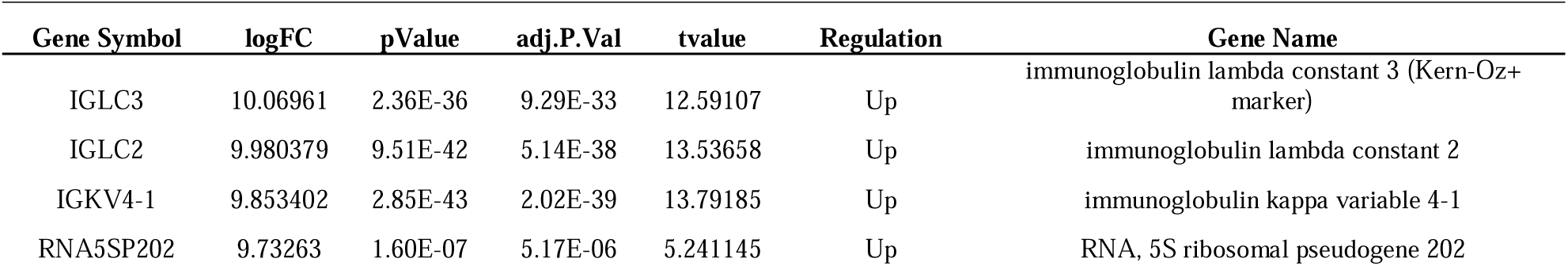
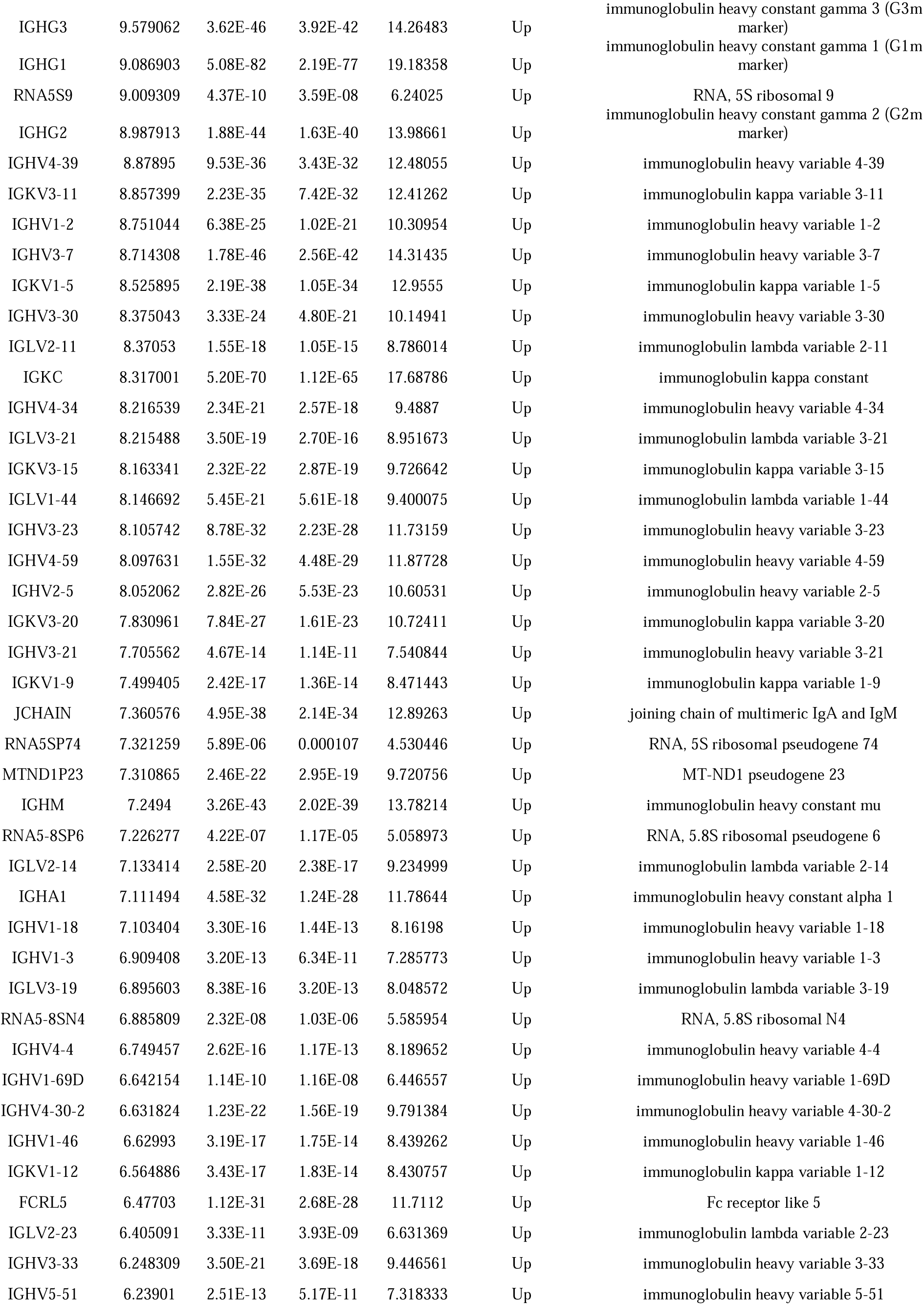
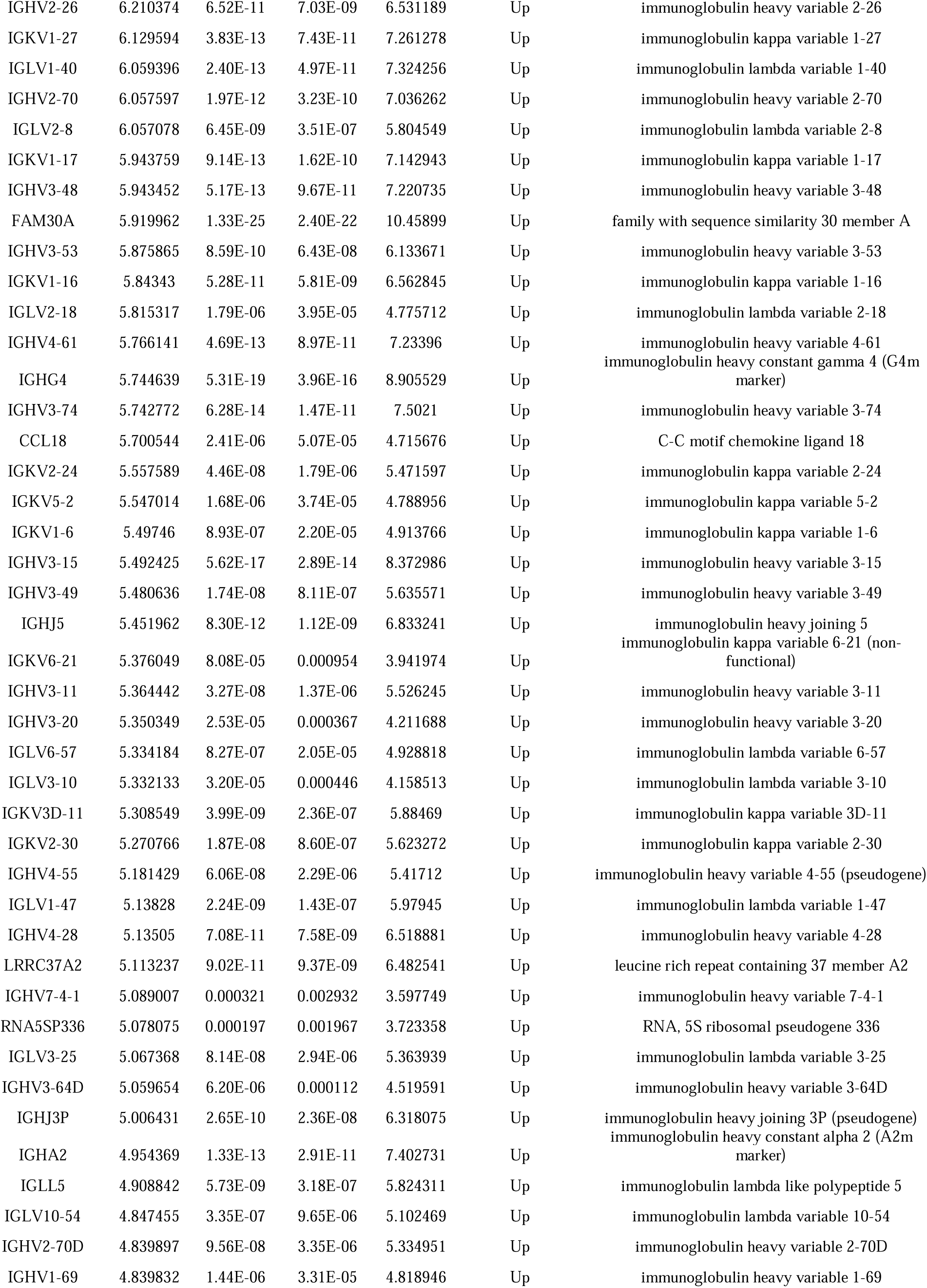
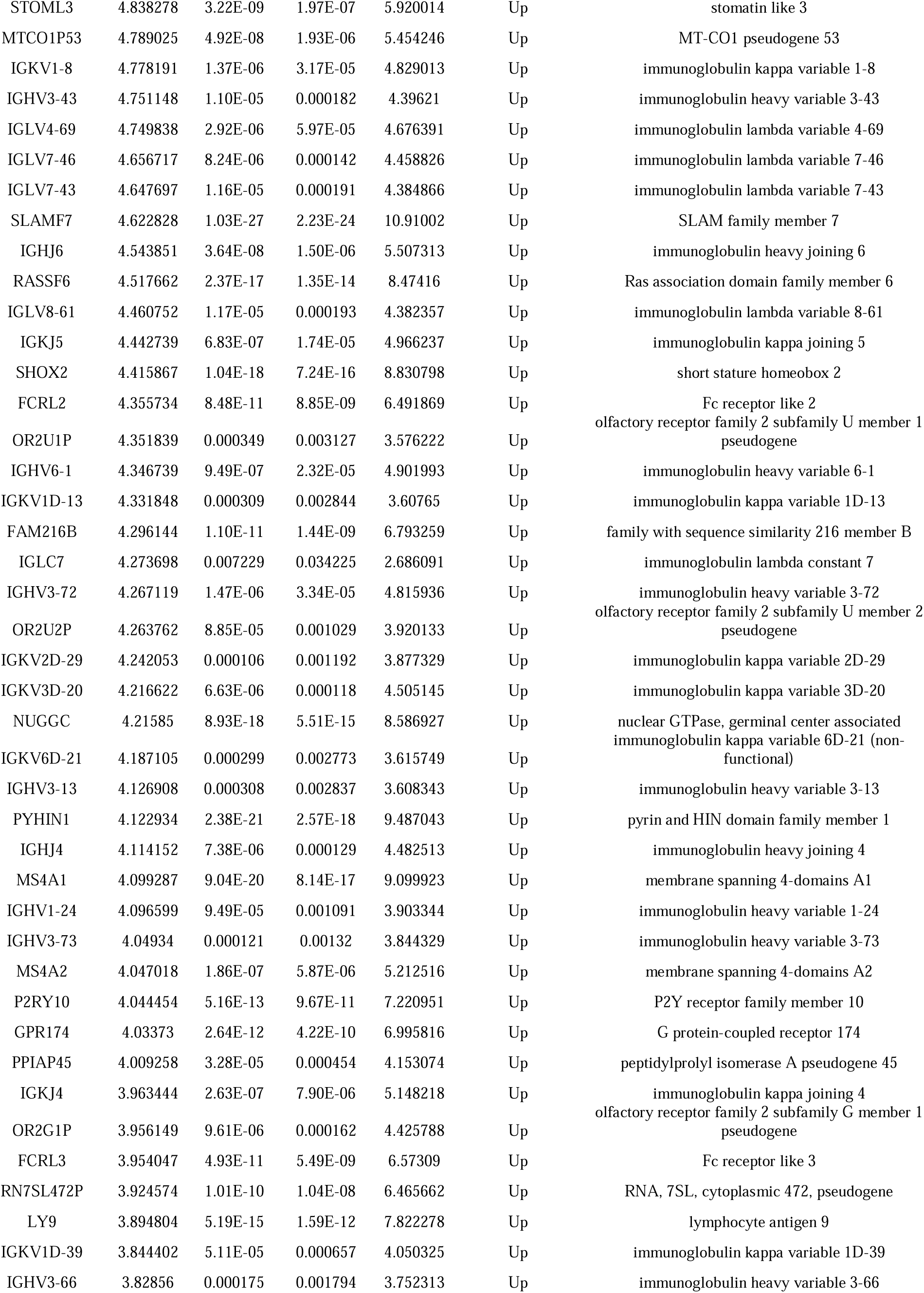
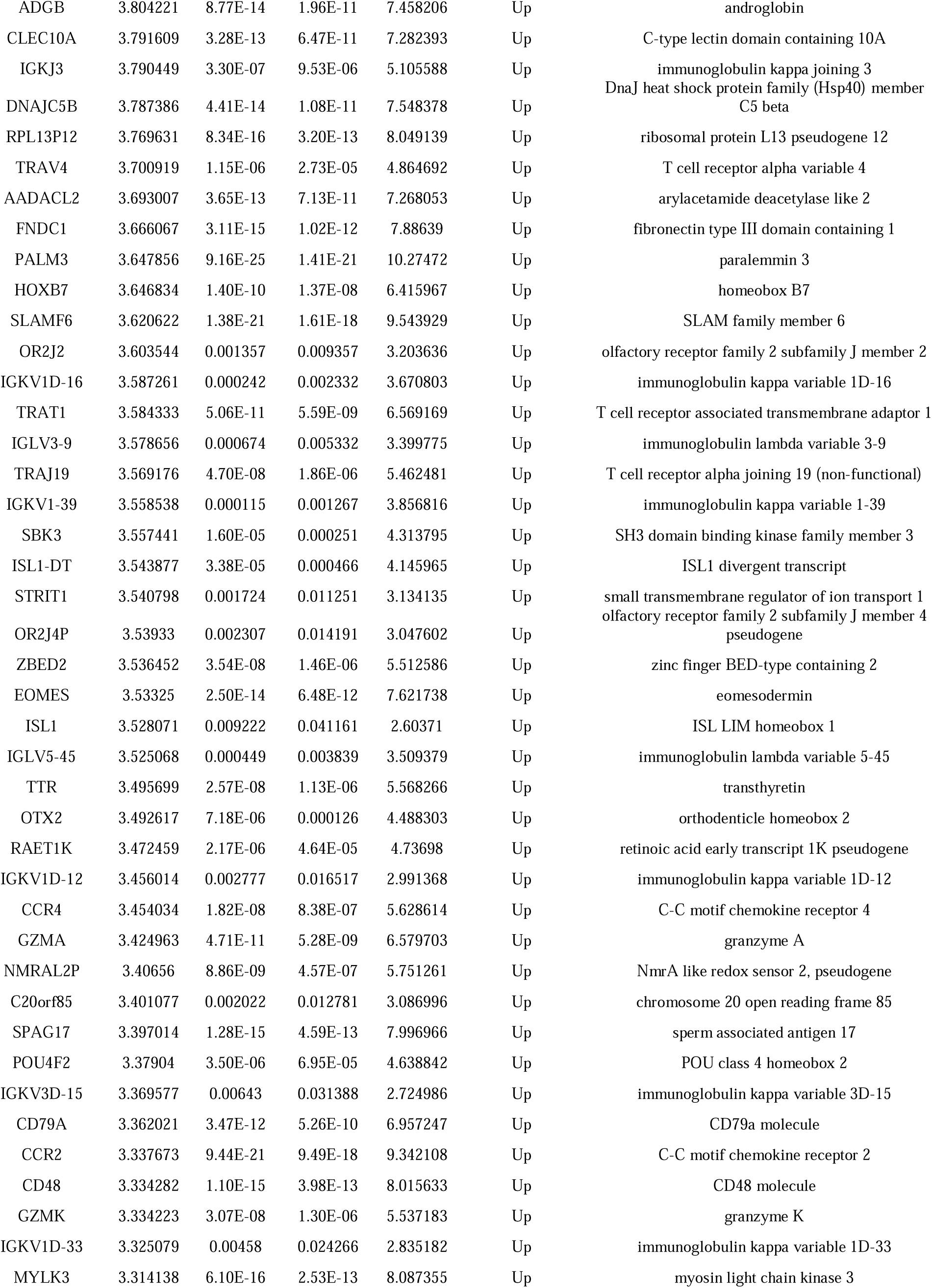
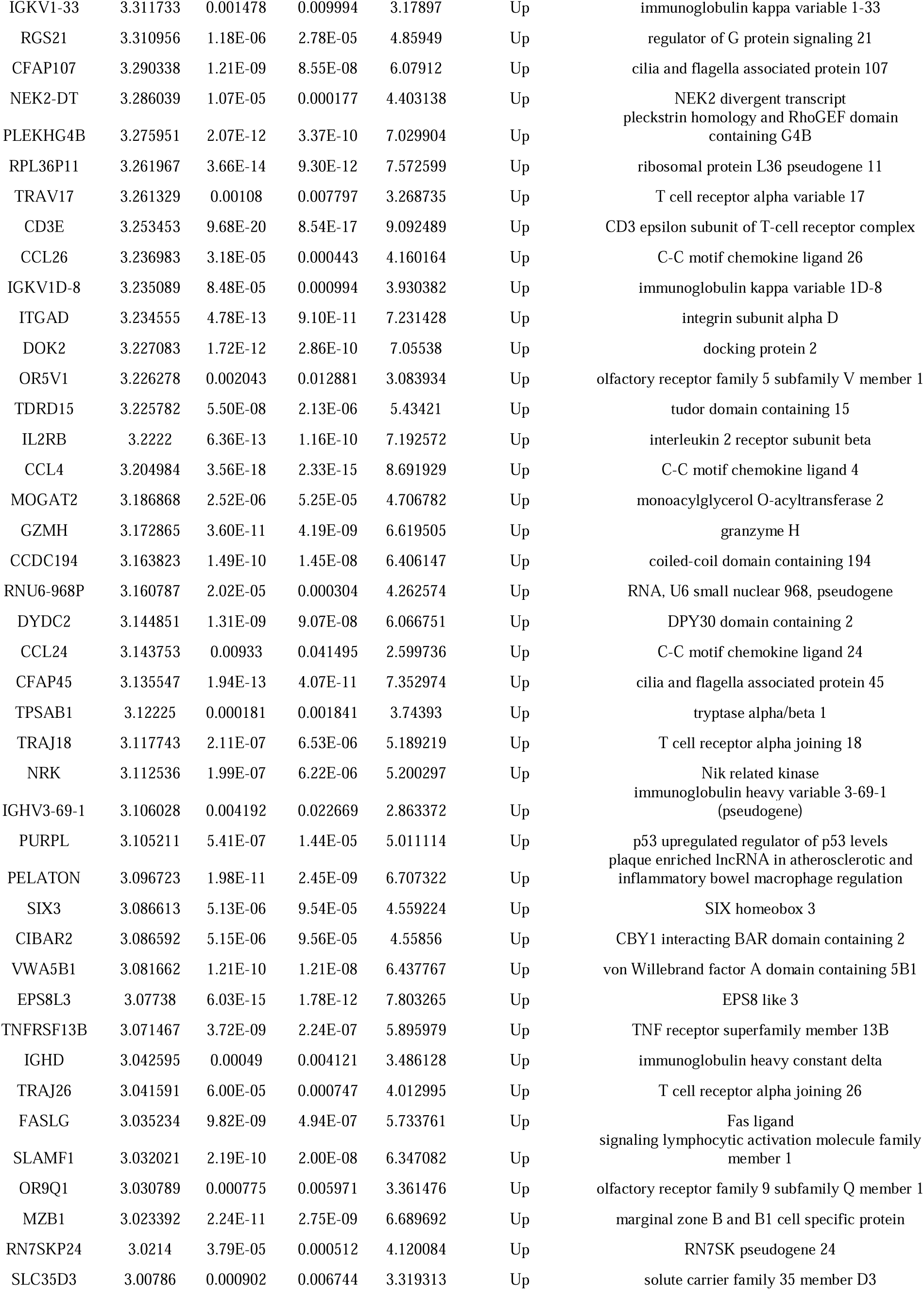
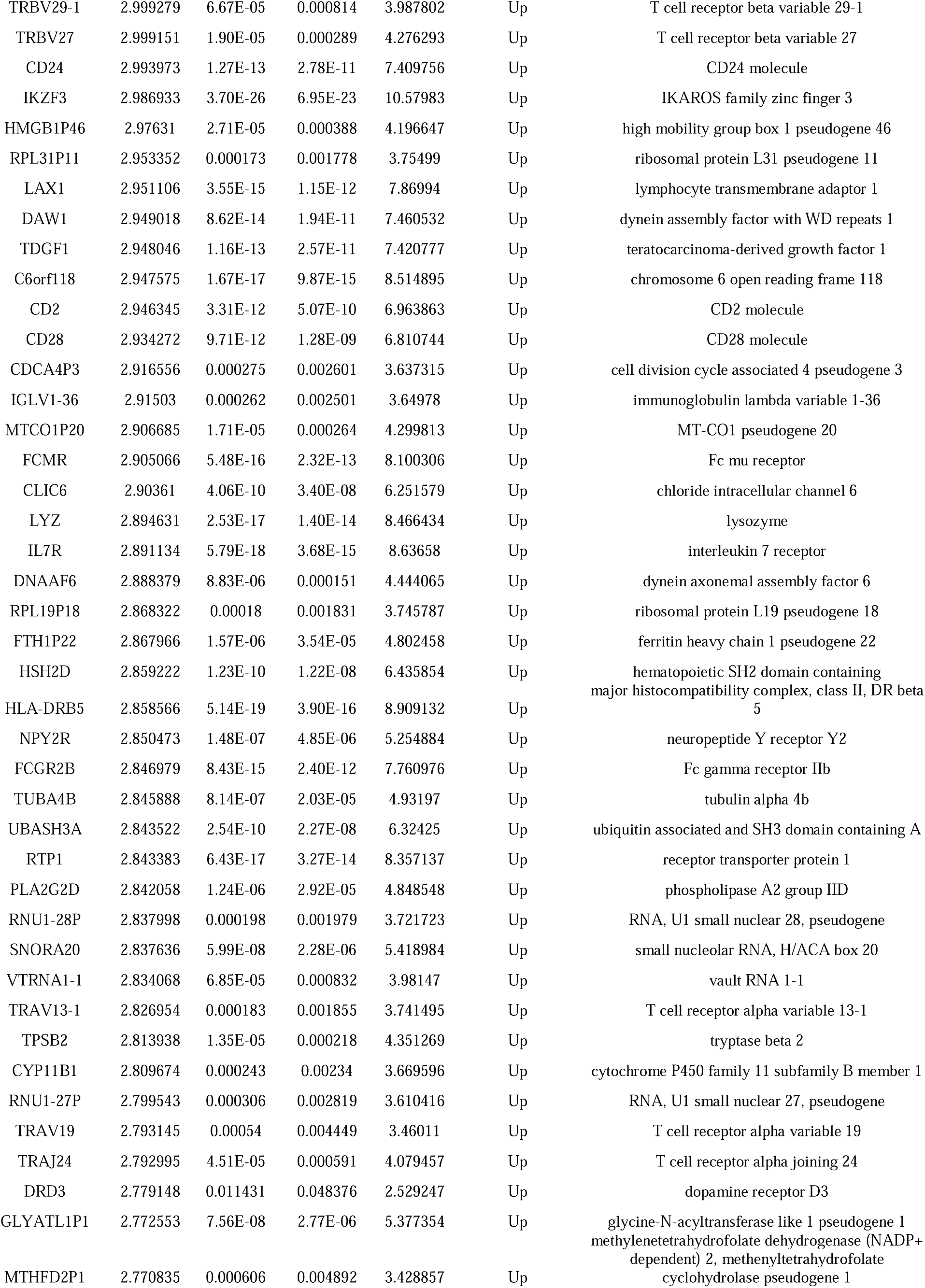
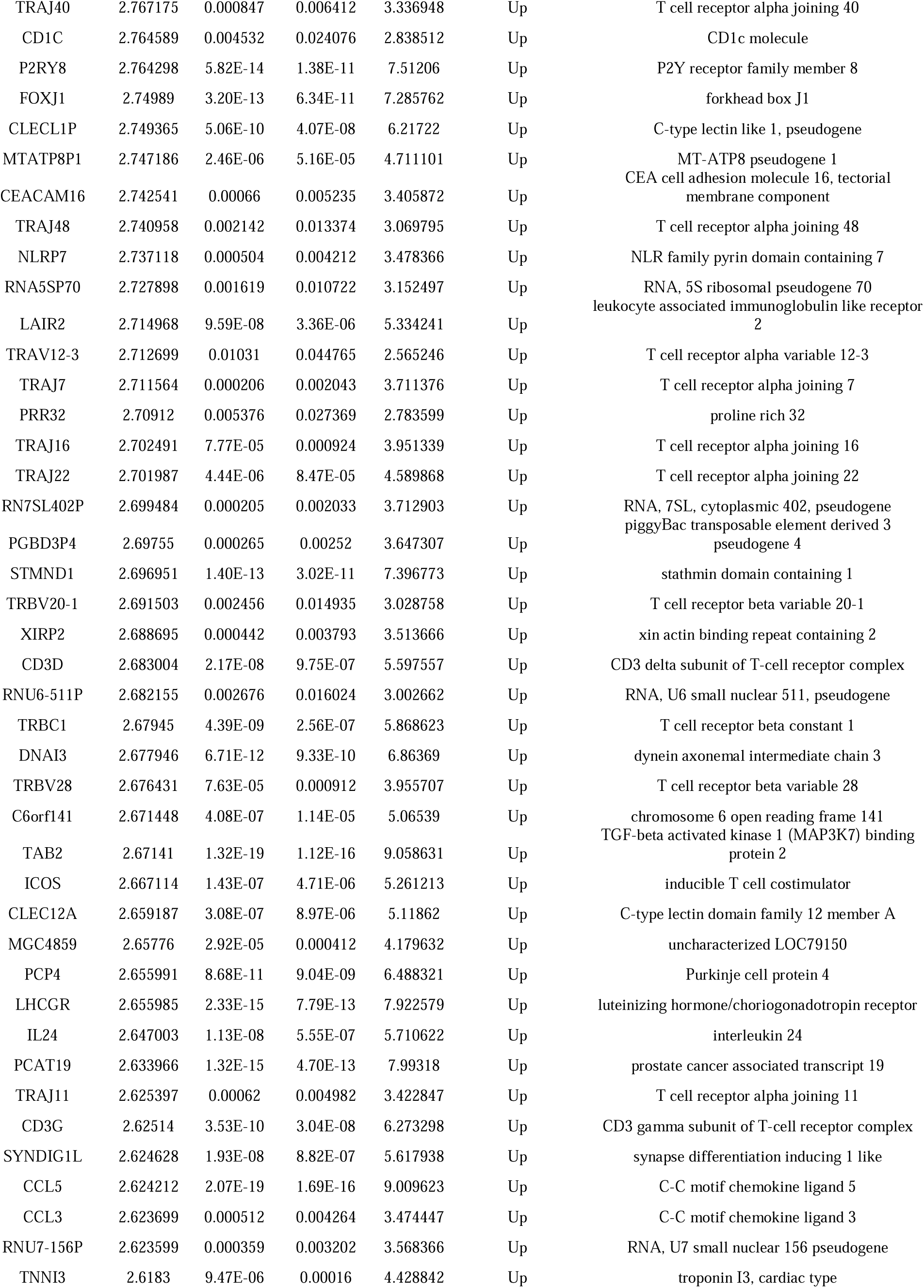
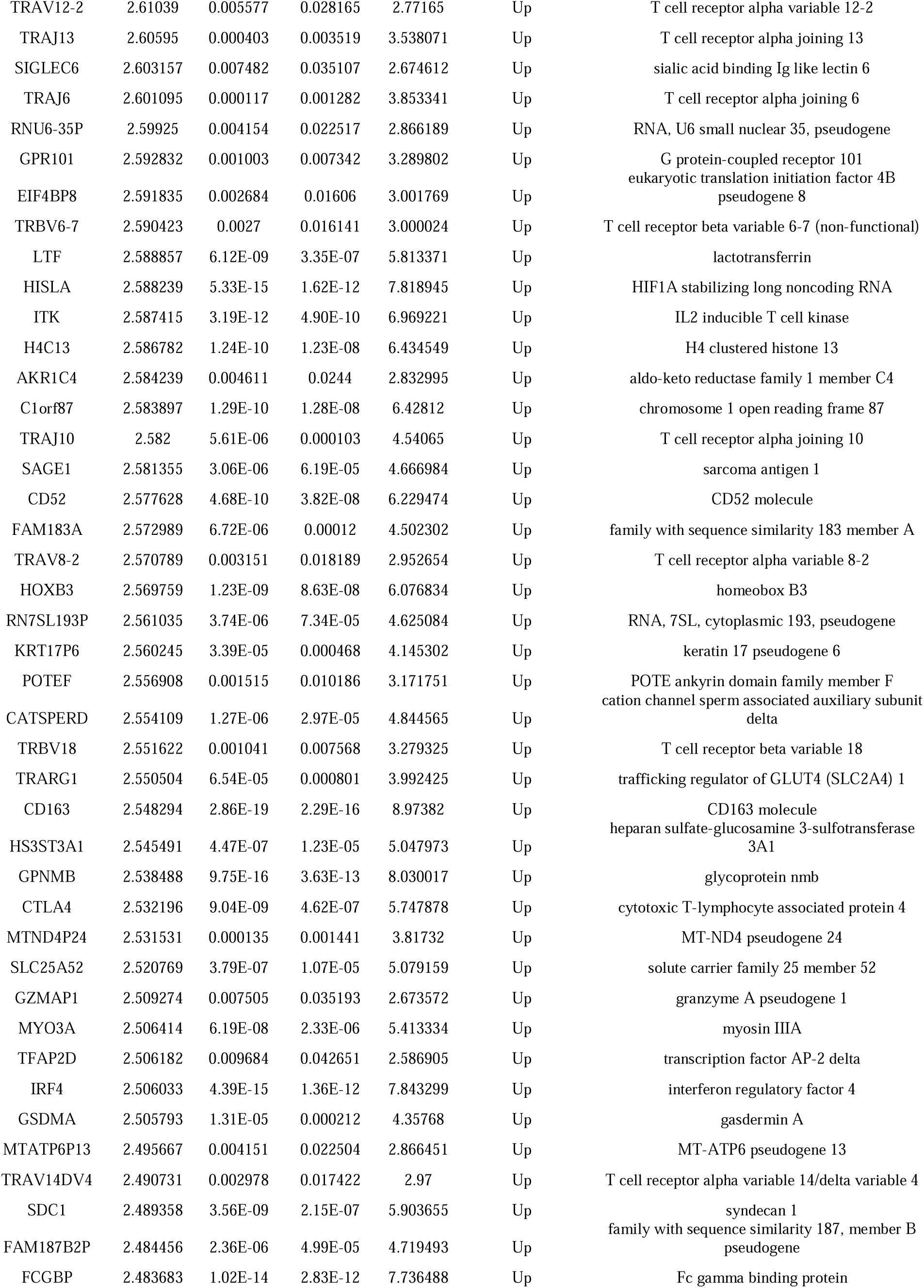
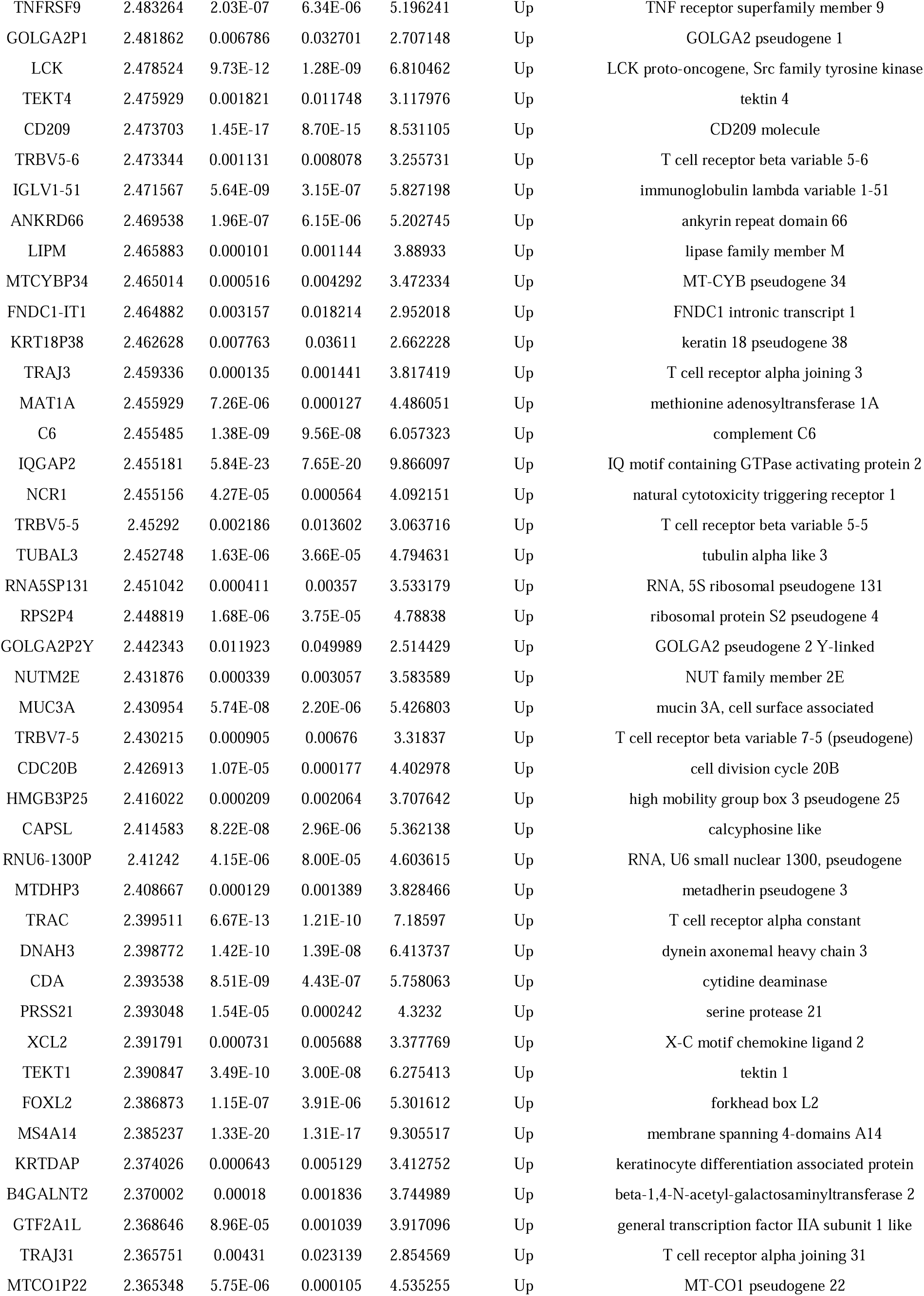
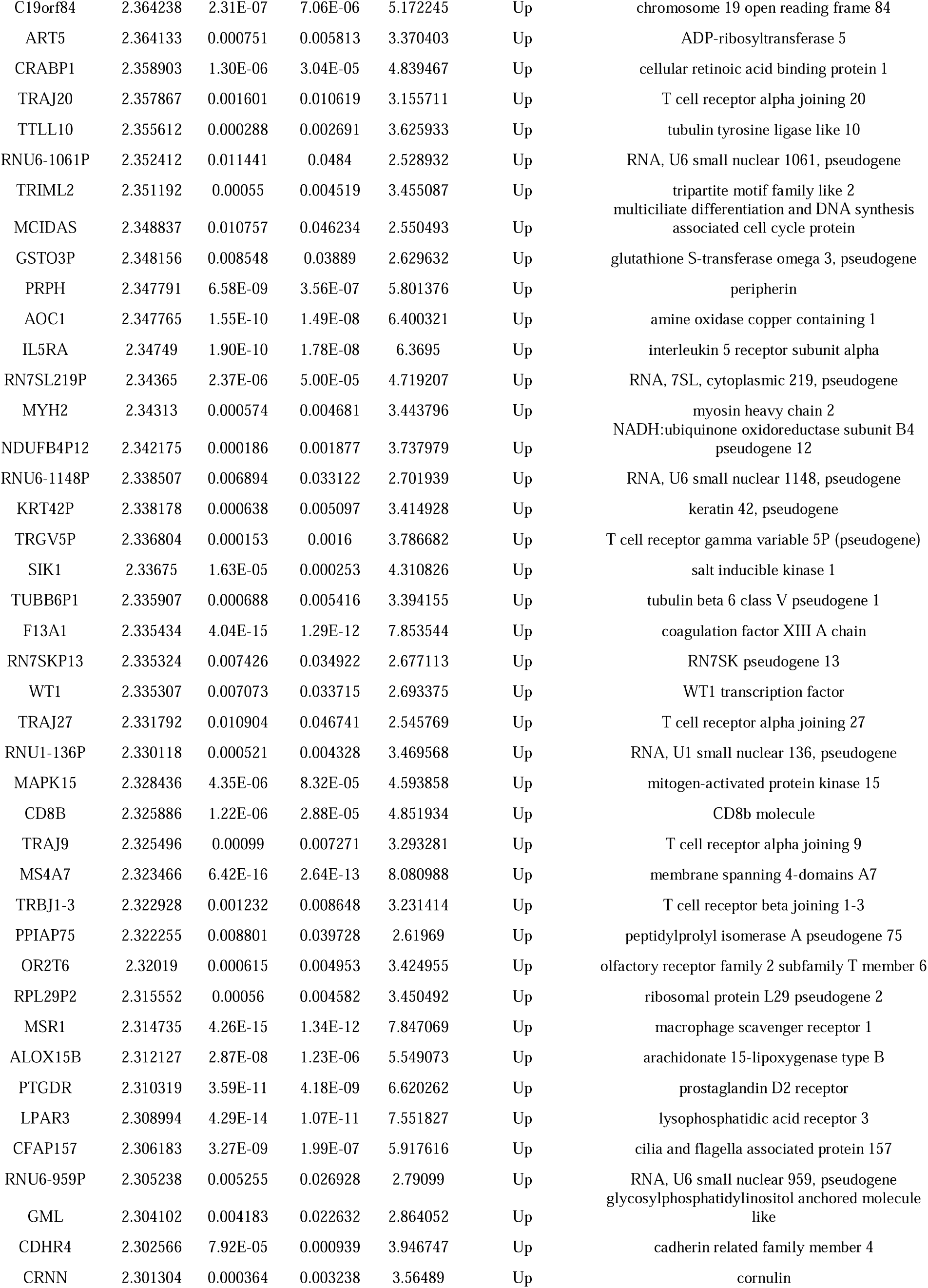
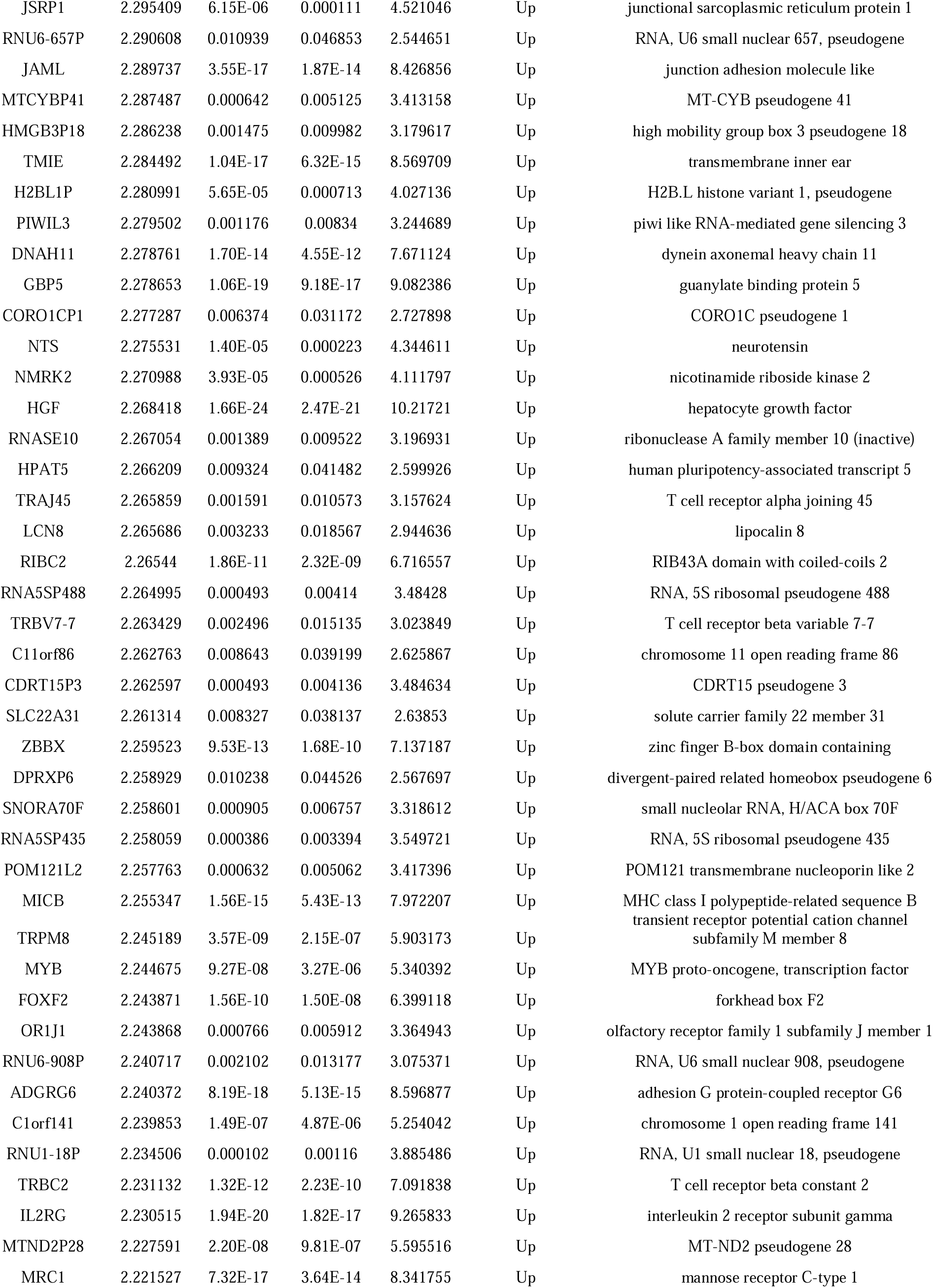
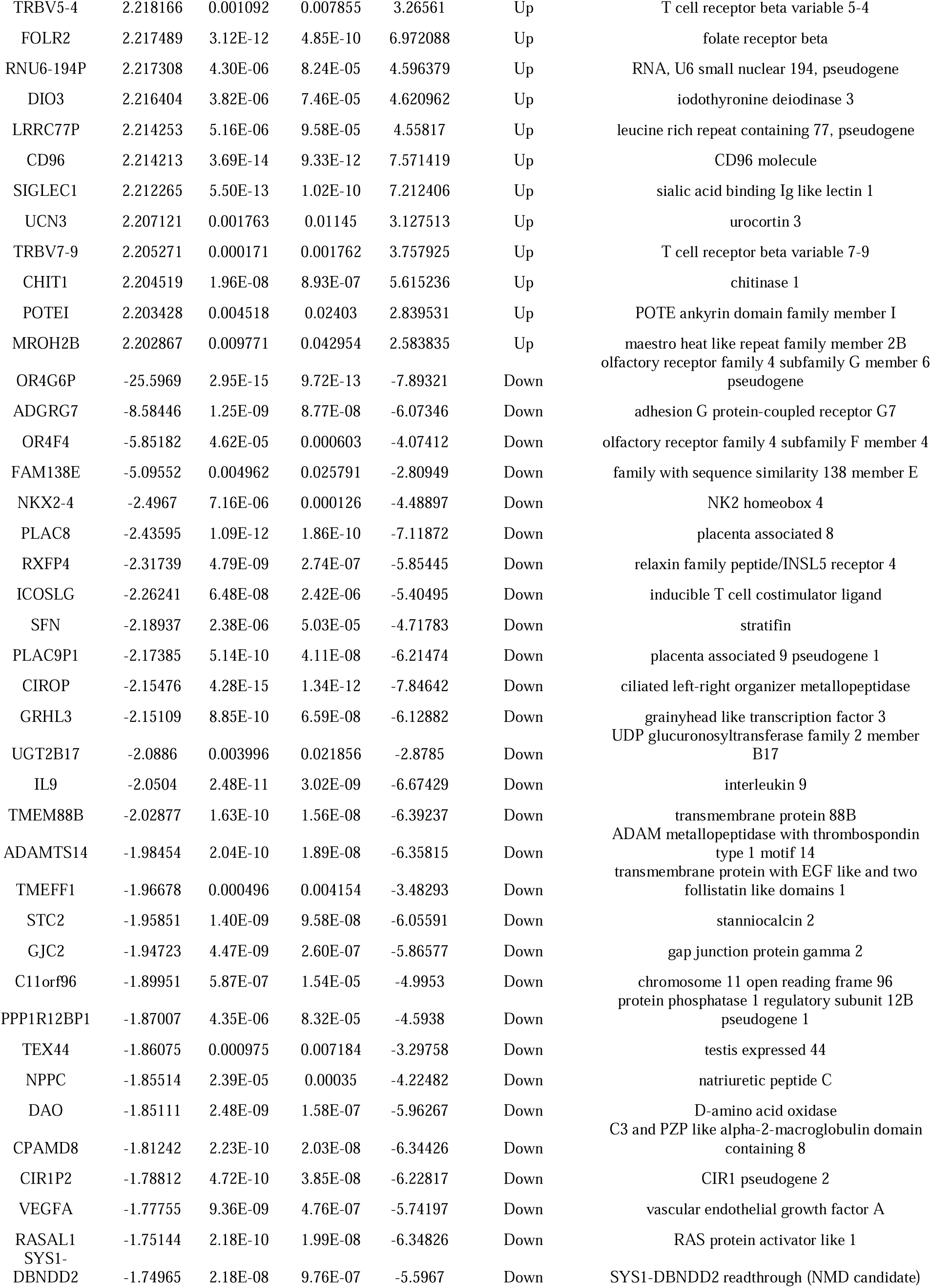
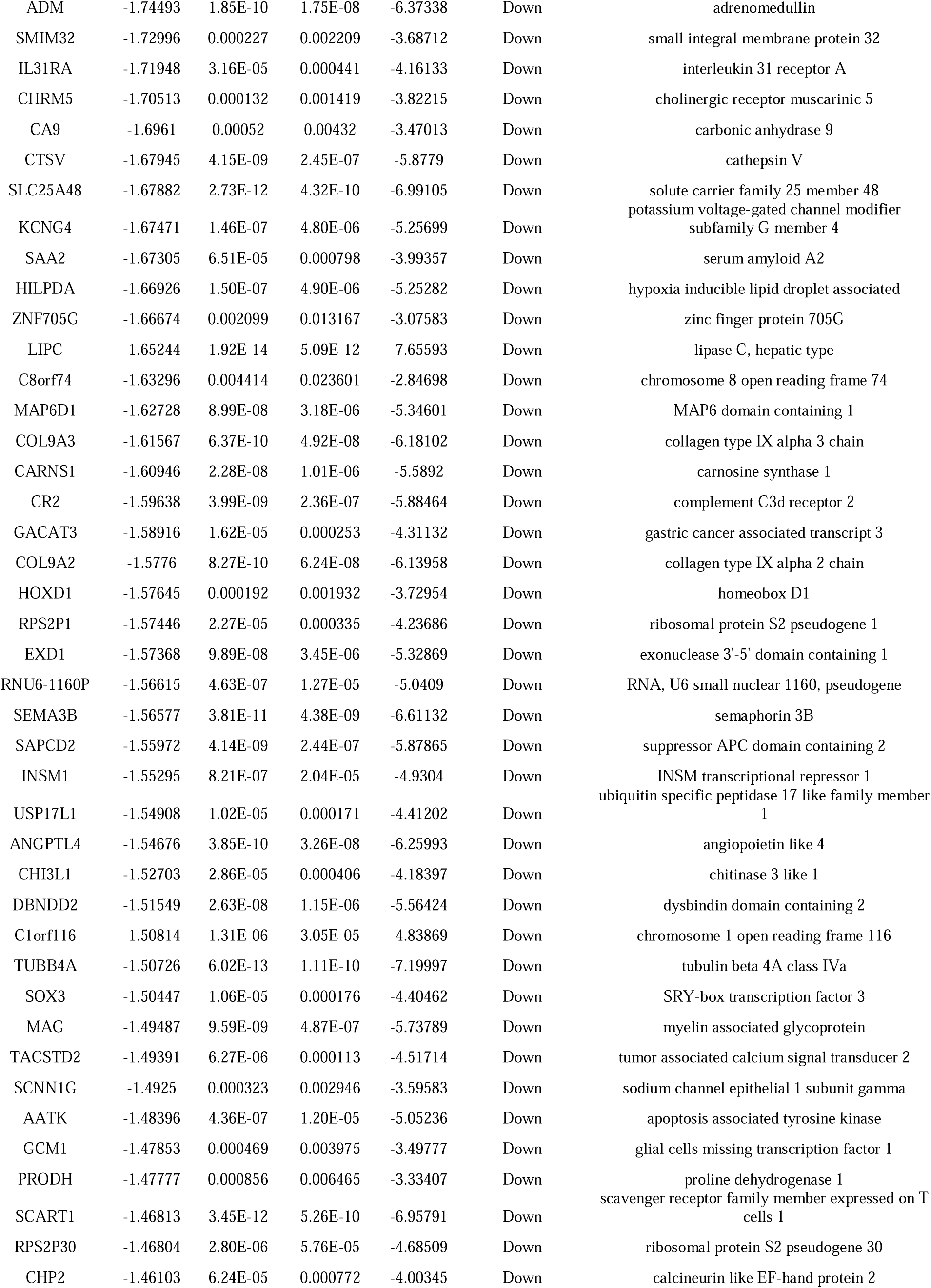
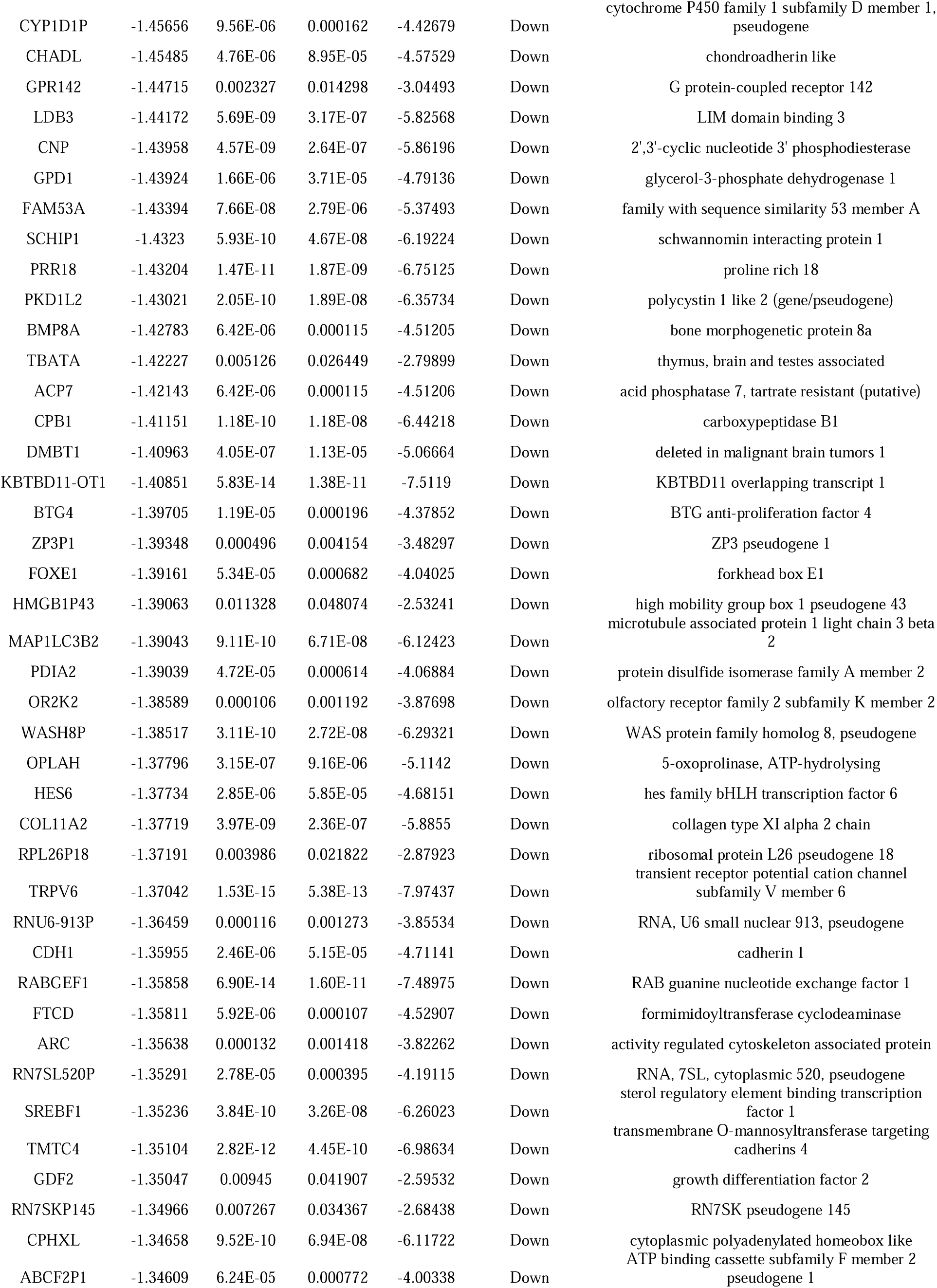
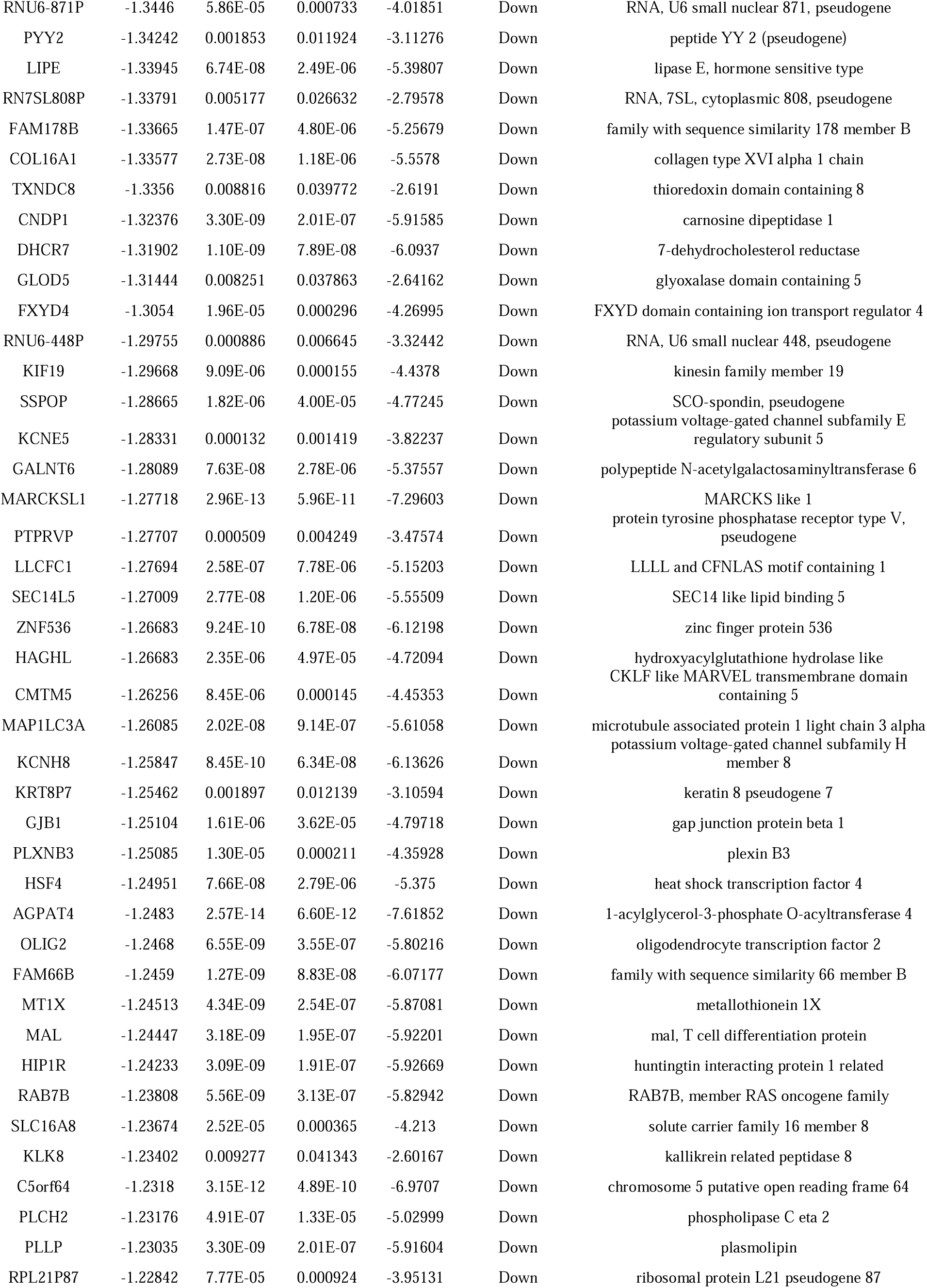
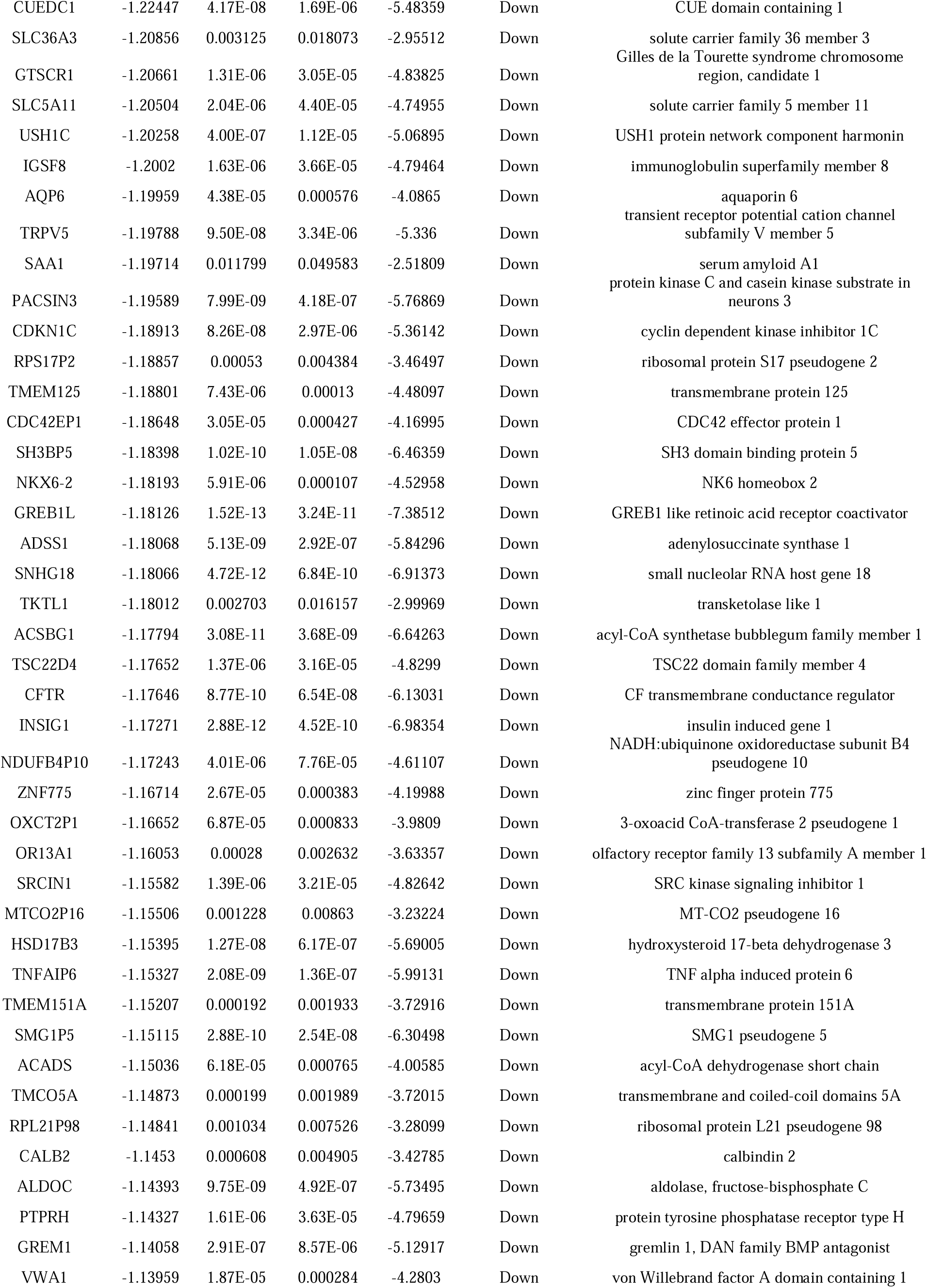
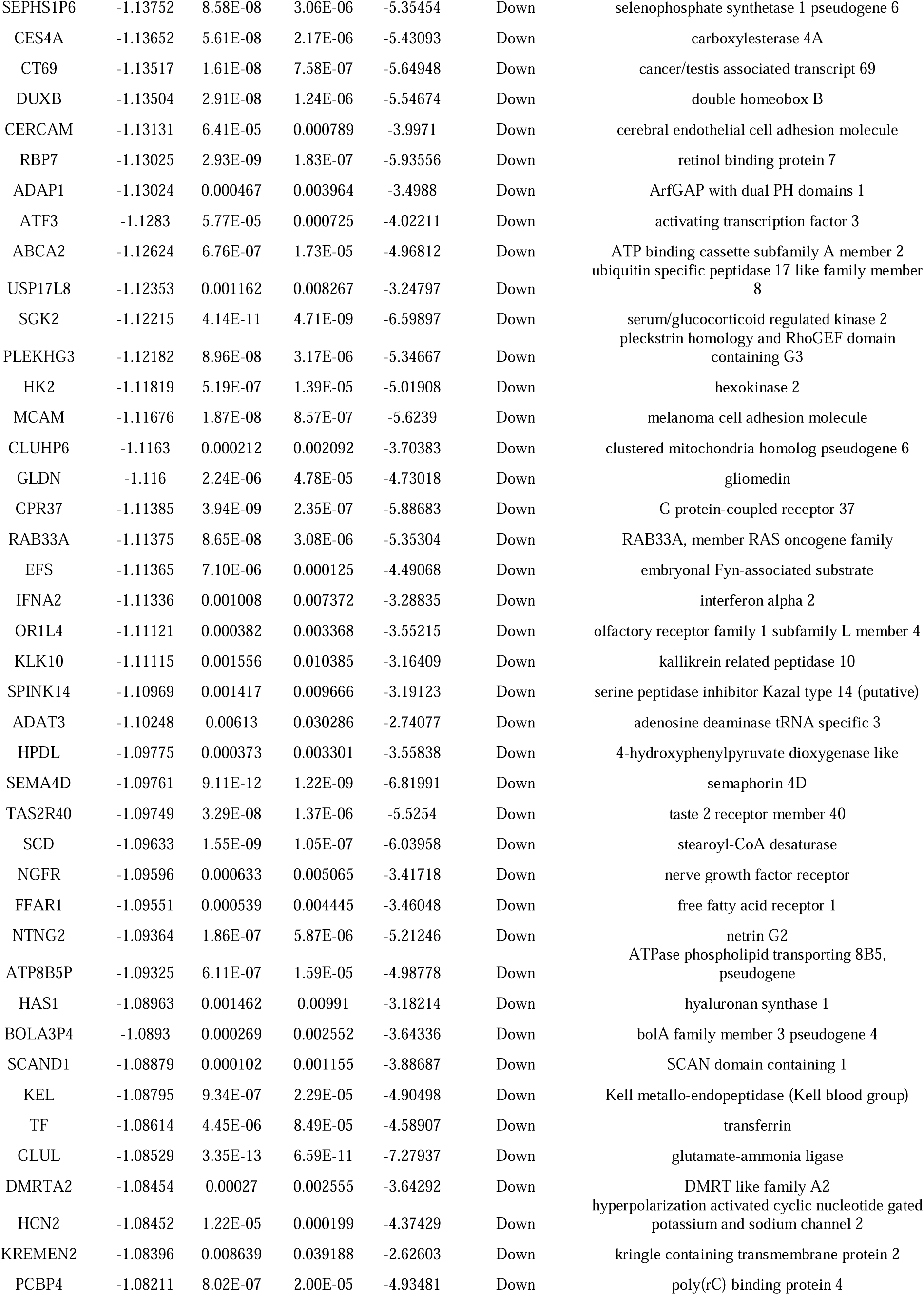
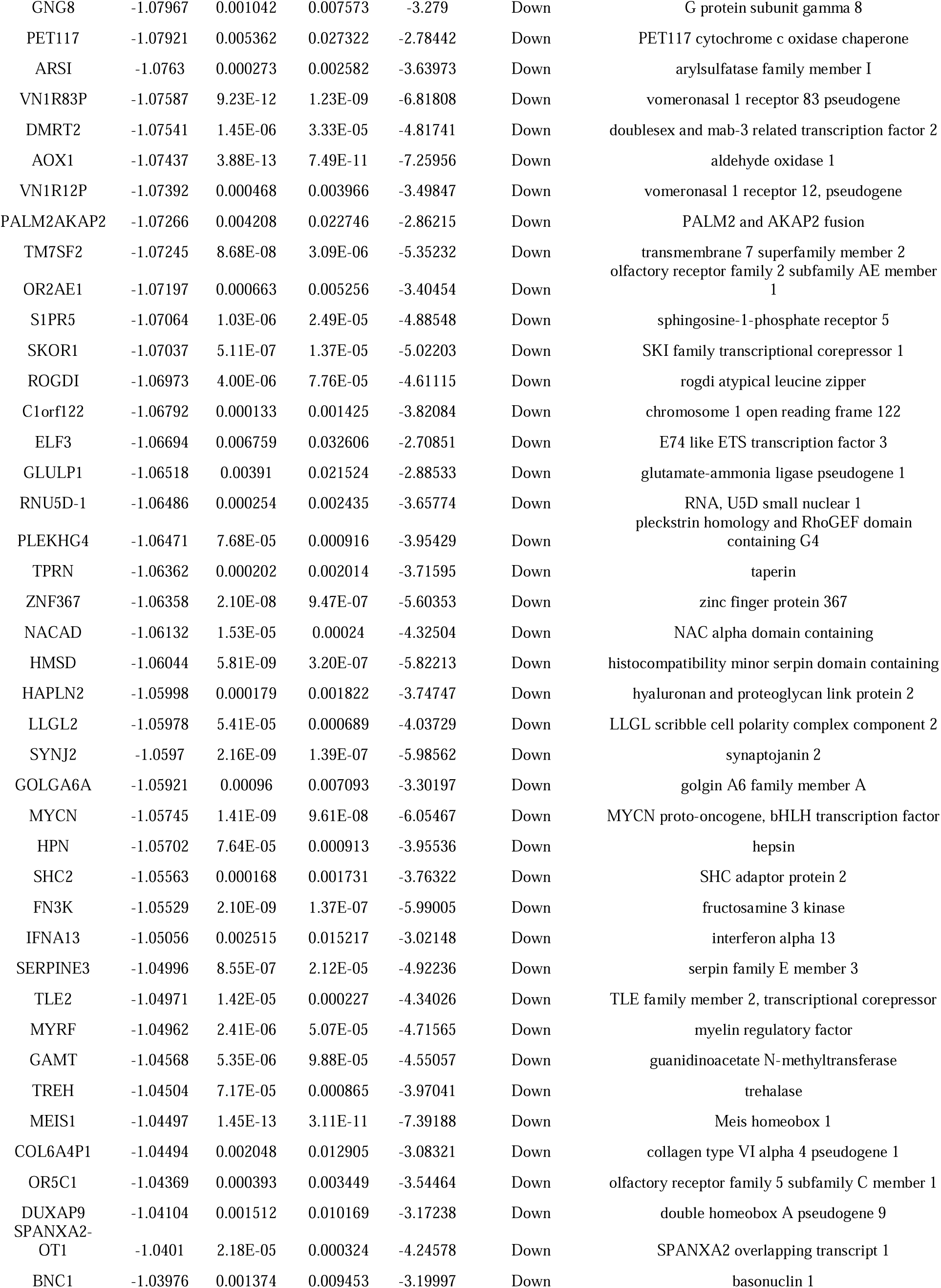
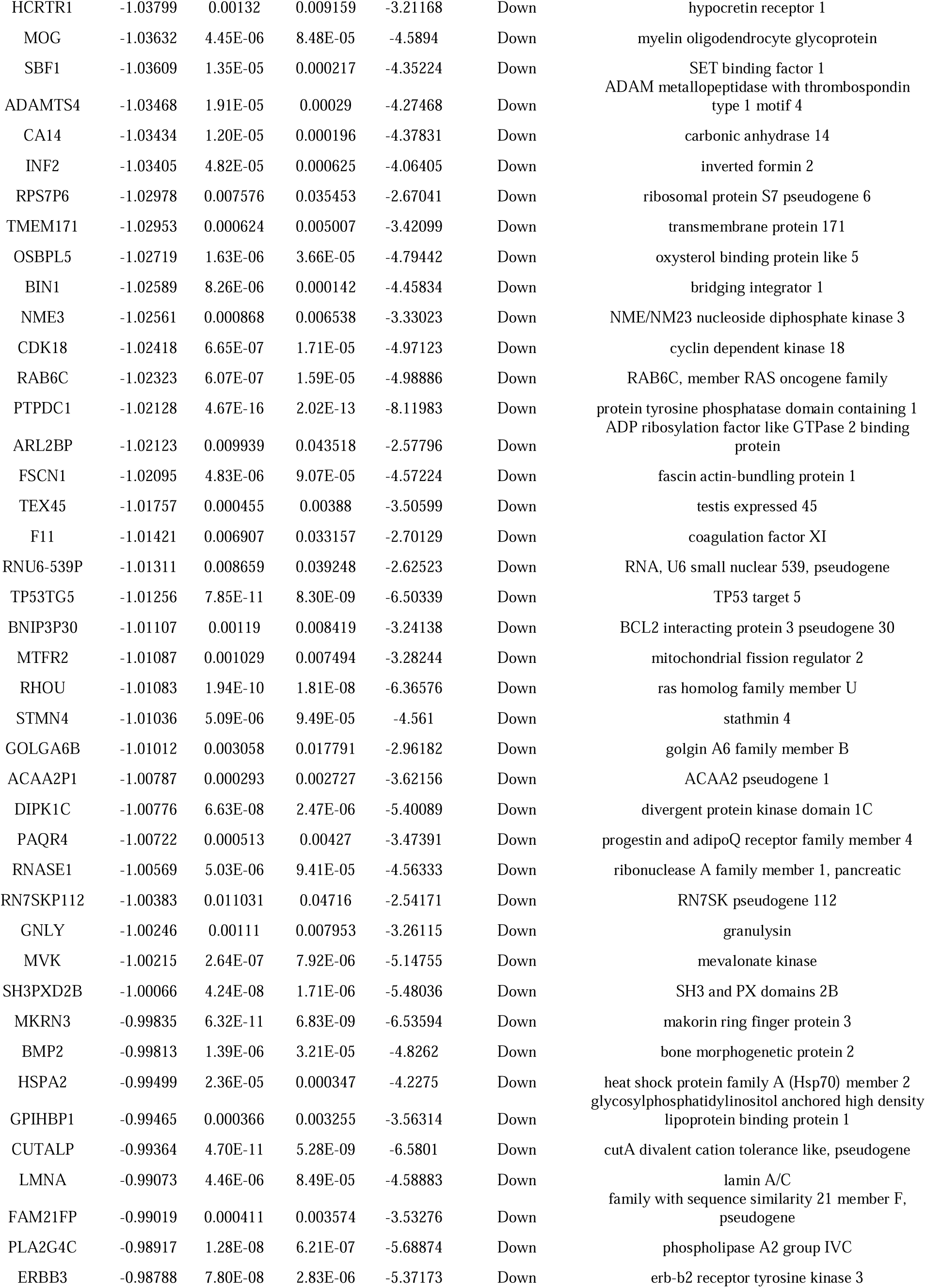
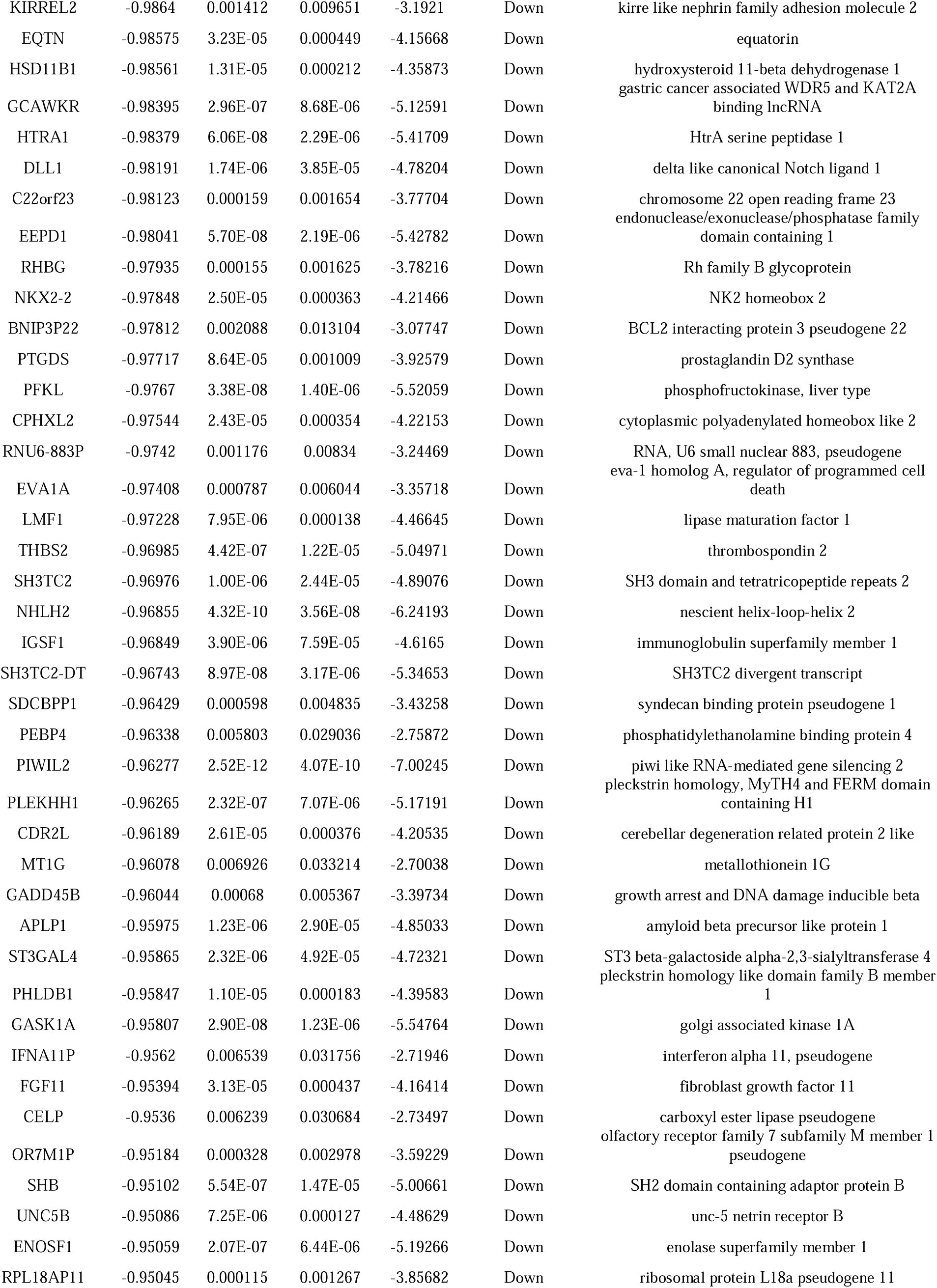
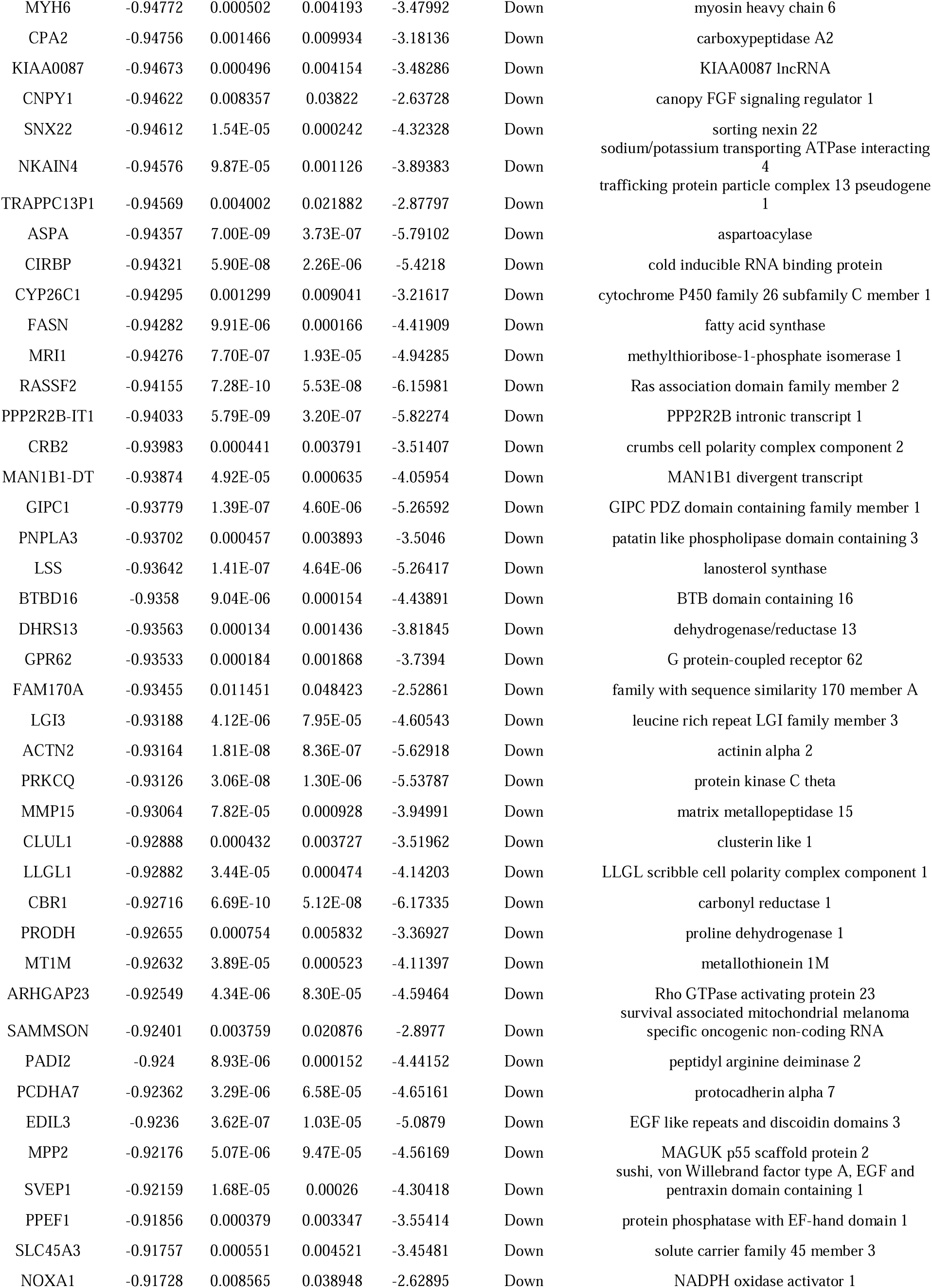
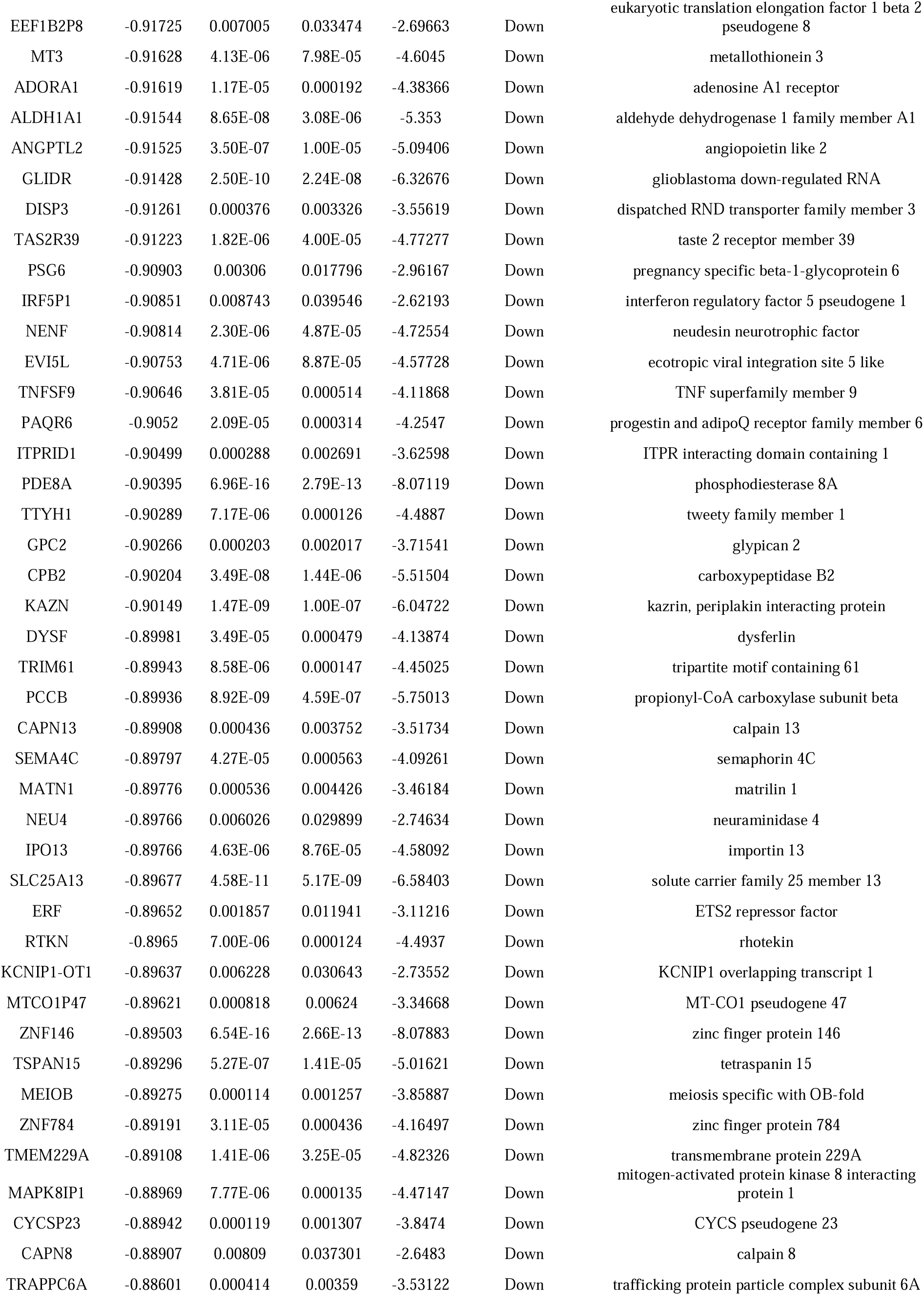
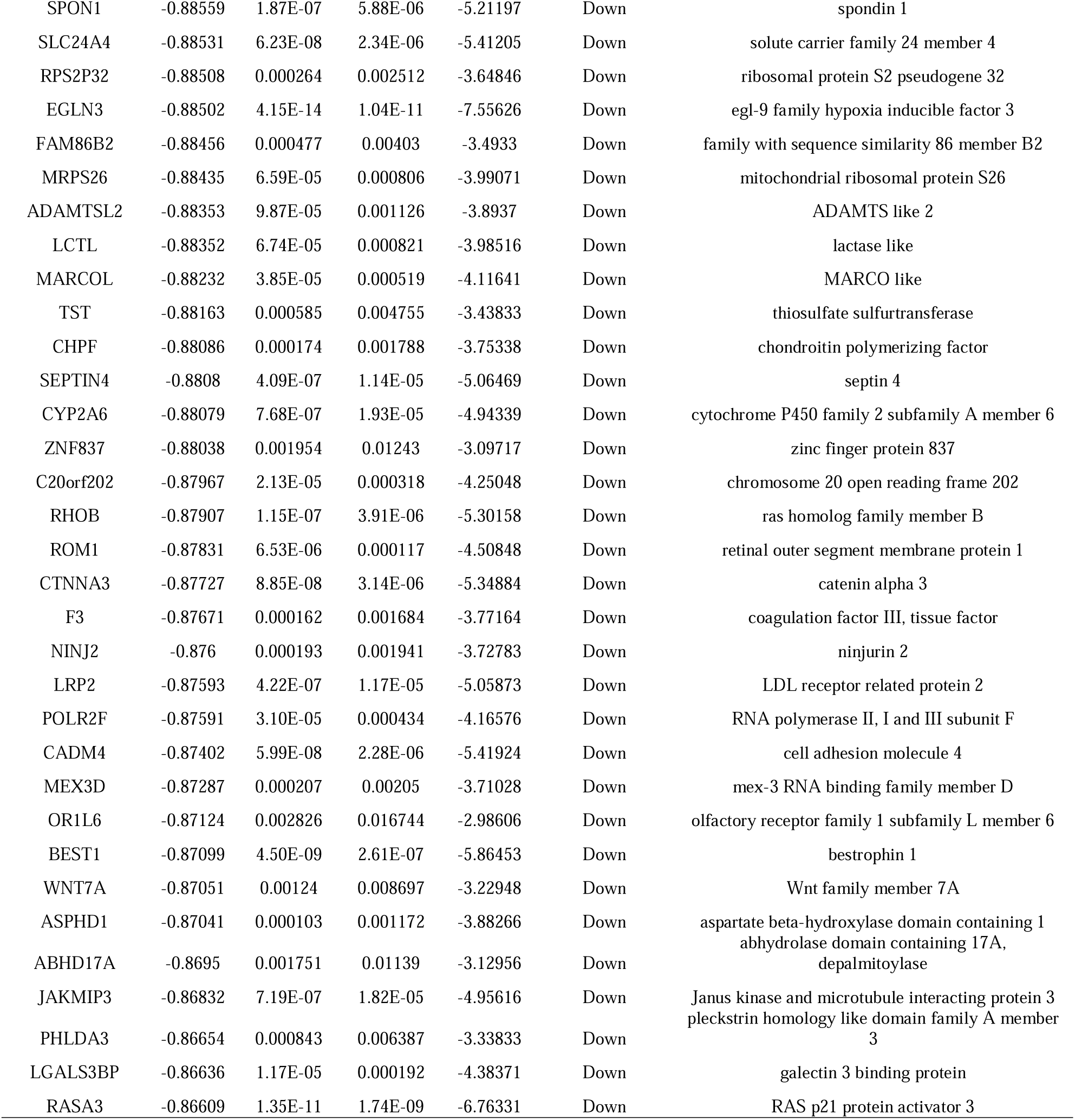
The statistical metrics for key differentially expressed genes (DEGs)

### GO and pathway enrichment analyses of DEGs

The DEGs were submitted to g:Profiler online tool for REACTOME pathway enrichment and GO enrichment analysis involving BP, CC and MF. Eligible terms for GO enrichment analysis of up and down regulated genes were shown in Fig. 4 and Fig. 5, and the number of enriched genes and adjusted P-value were shown in Table 3. From the figures and tables, it is evident that the DEGs were primarily enriched in immune system process, response to stimulus, developmental process and multicellular organismal process (BP); membrane, protein-containing complex, cellular anatomical entity and cytoplasm (CC); antigen binding, signaling receptor binding, molecular function regulator activity and molecular function activator activity (MF). Eligible terms for pathway enrichment analysis of up and down regulated genes were shown in were shown in Fig. 4 and Fig. 5, and the number of enriched genes and adjusted P-value were shown in Table 4. From the figures and tables, it is evident that the DEGs were primarily enriched in immune system, TCR signaling, regulation of cholesterol biosynthesis by SREBP (SREBF) and degradation of the extracellular matrix.

**Fig. 4.**
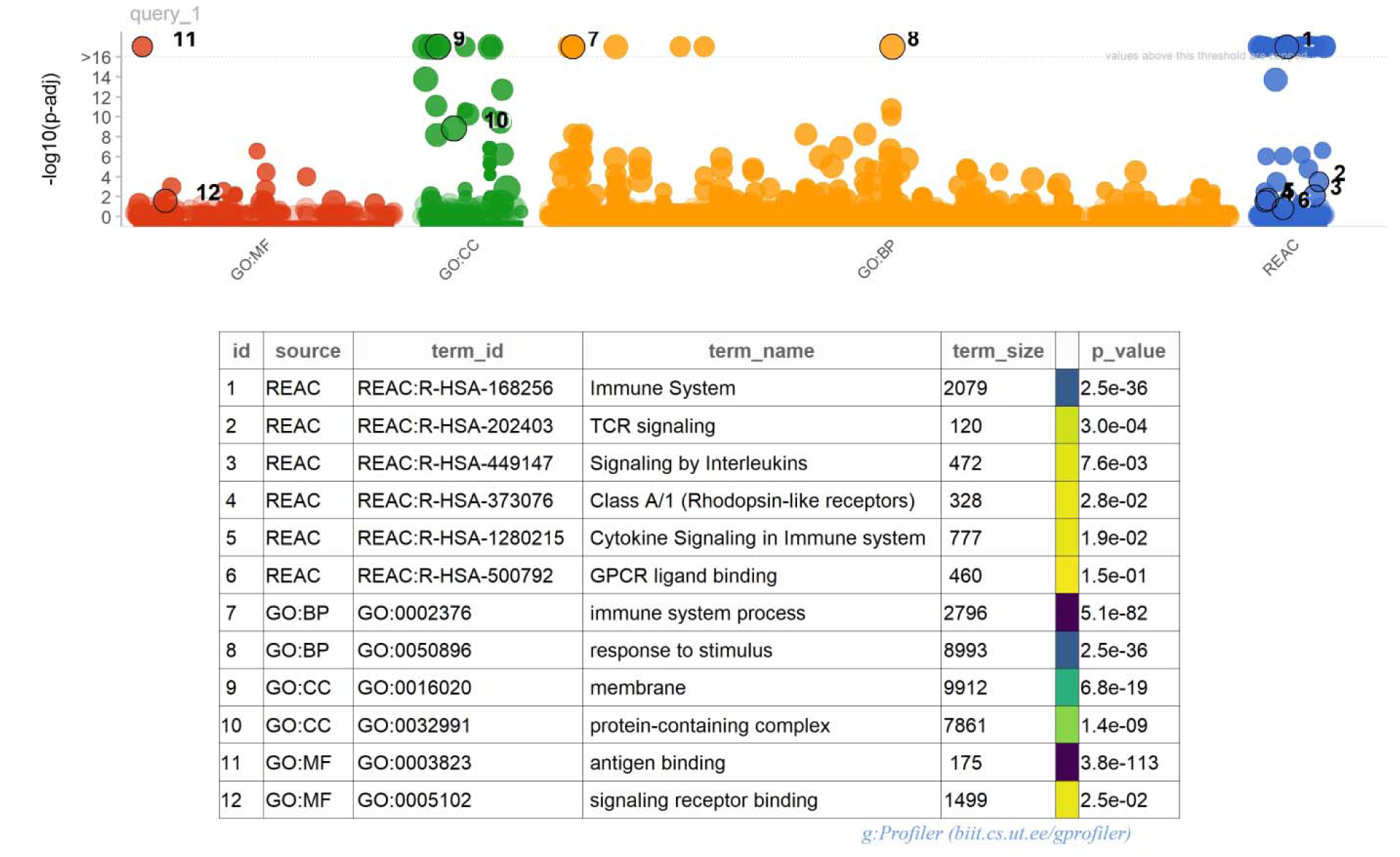
GO and REACTOME pathway enrichment analysis for up regulated genes. p < 0.05. Abbreviations: BP, biological process; CC, cell component; MF, molecular function. GO, Gene Ontology; REAC, REACTOME. The size of the circle represents the number of genes involved, and the abscissa represents the frequency of the genes involved in the term total genes.

**Fig. 5.**
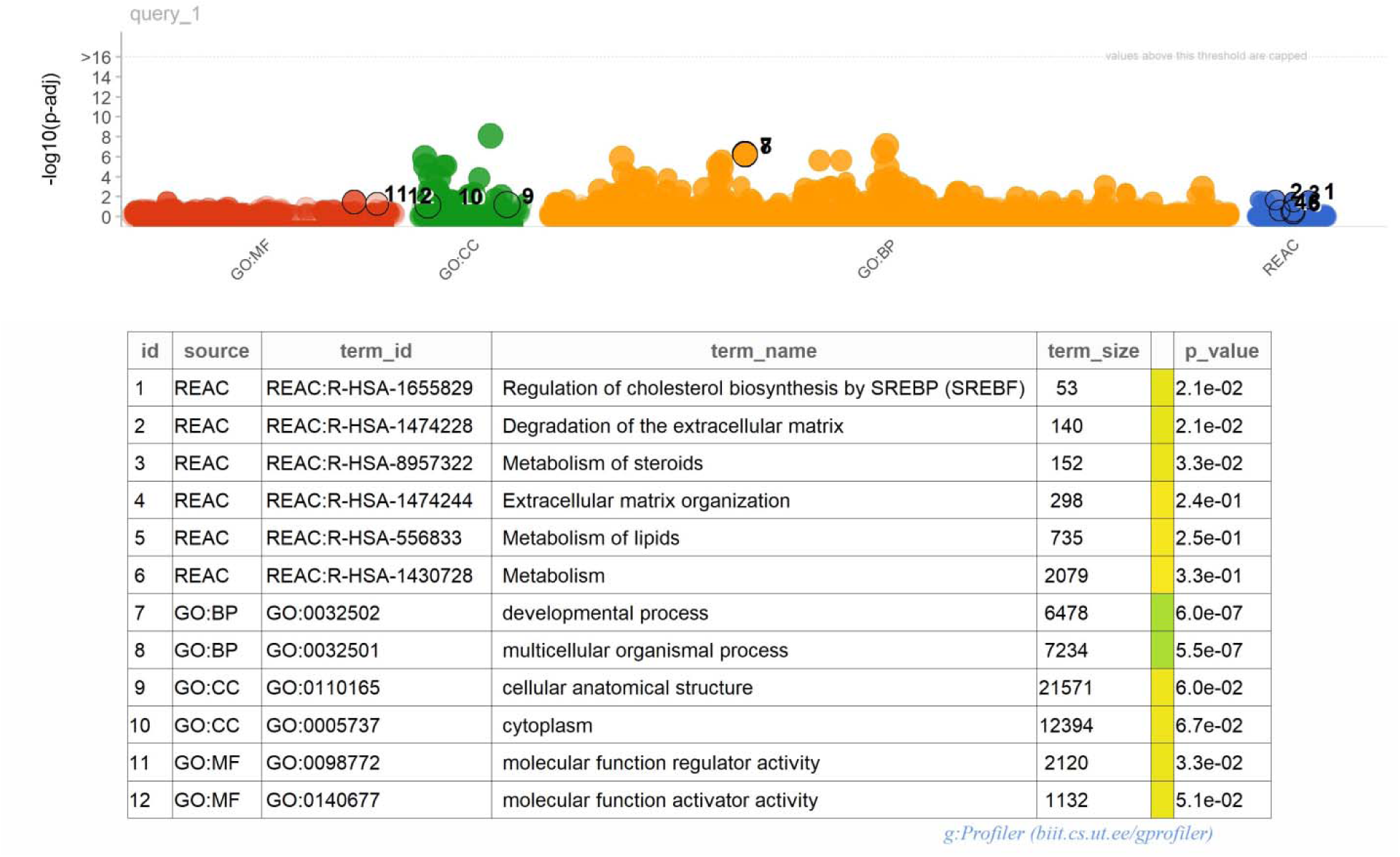
GO and REACTOME pathway enrichment analysis for down regulated genes. p < 0.05. Abbreviations: BP, biological process; CC, cell component; MF, molecular function. GO, Gene Ontology; REAC, REACTOME. The size of the circle represents the number of genes involved, and the abscissa represents the frequency of the genes involved in the term total genes.

**Table 3.**
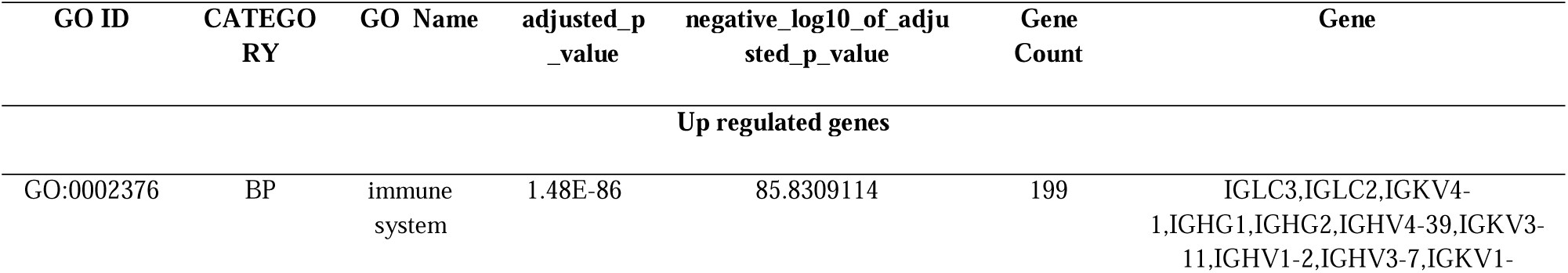
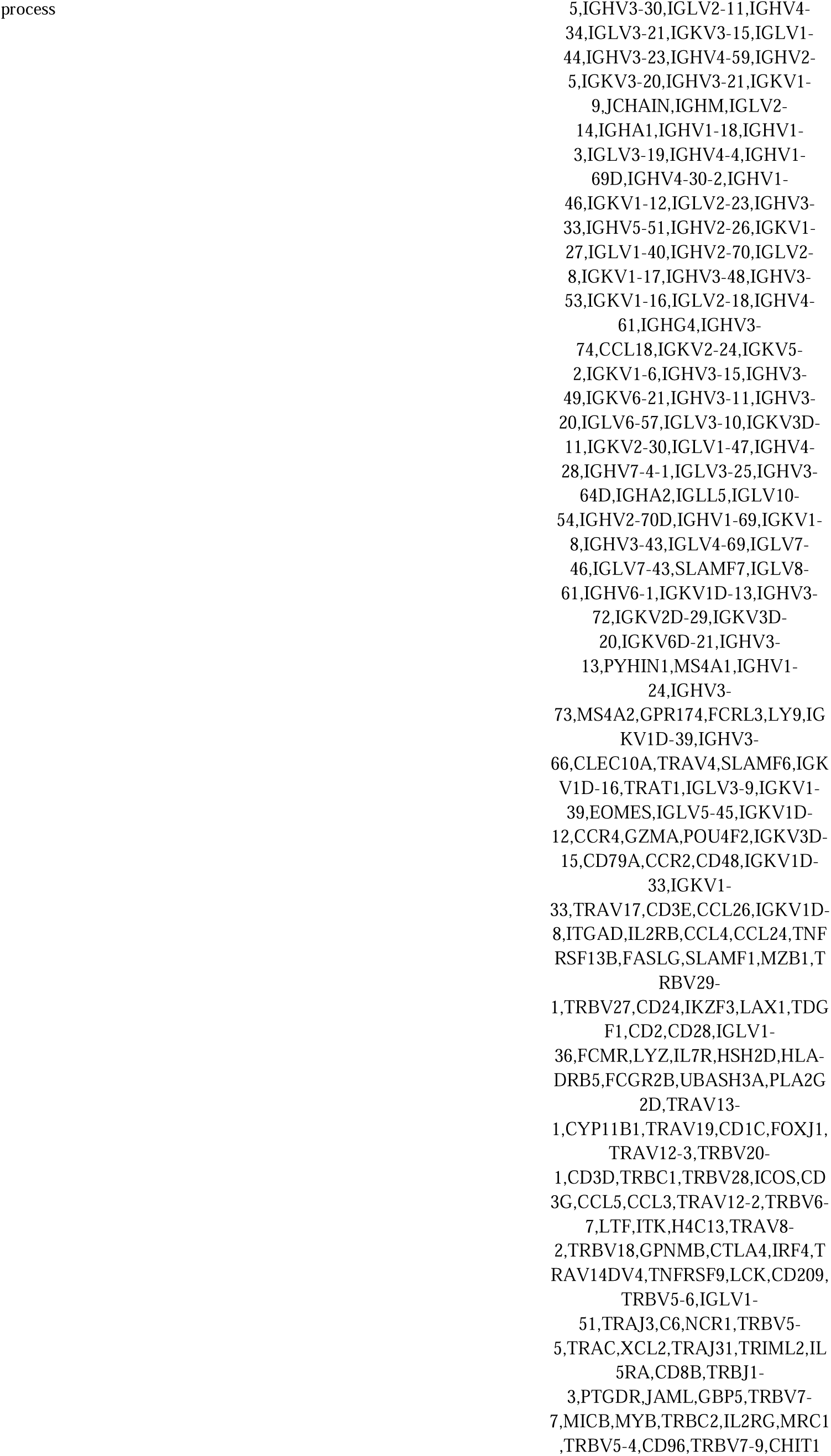
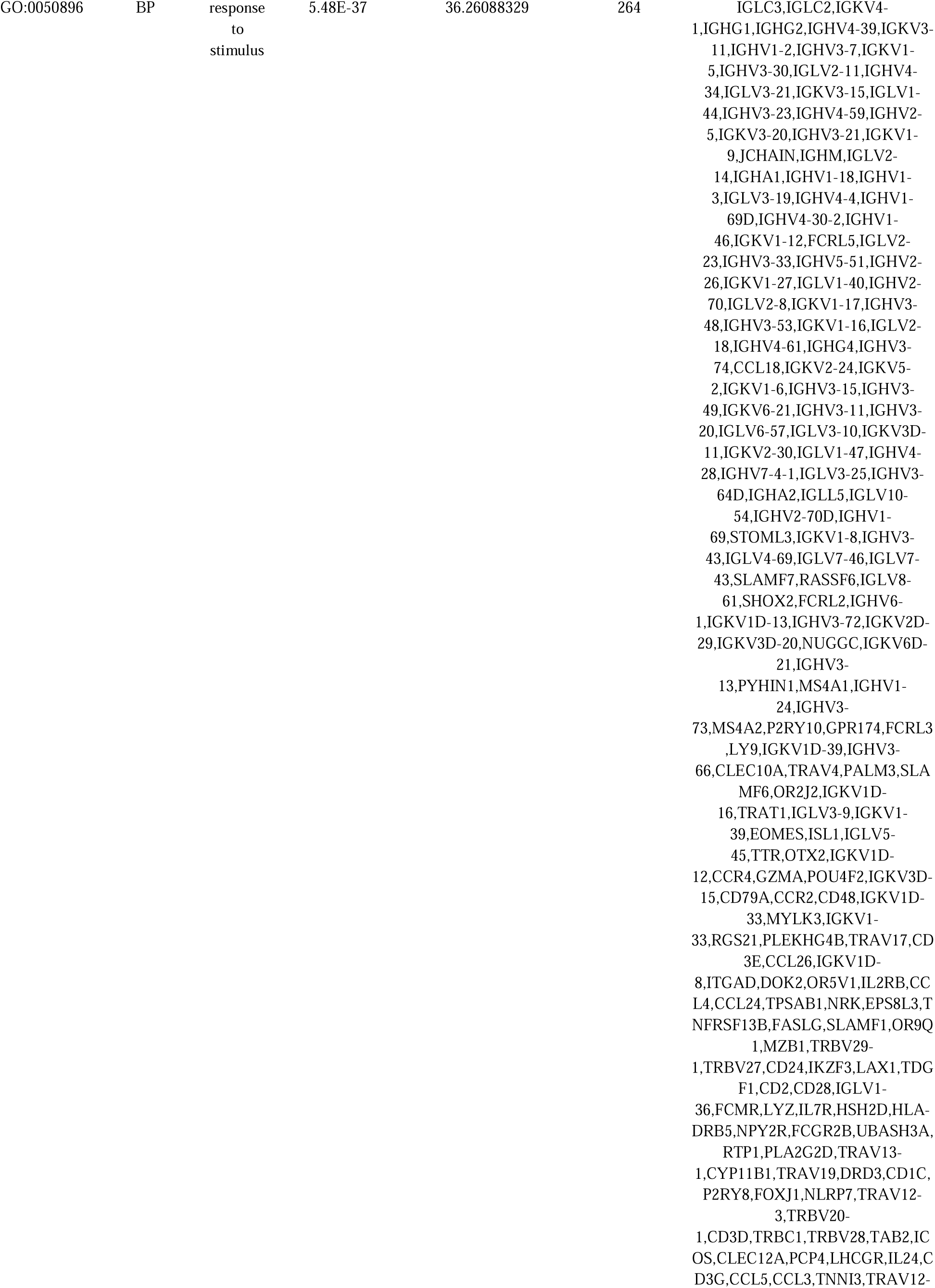
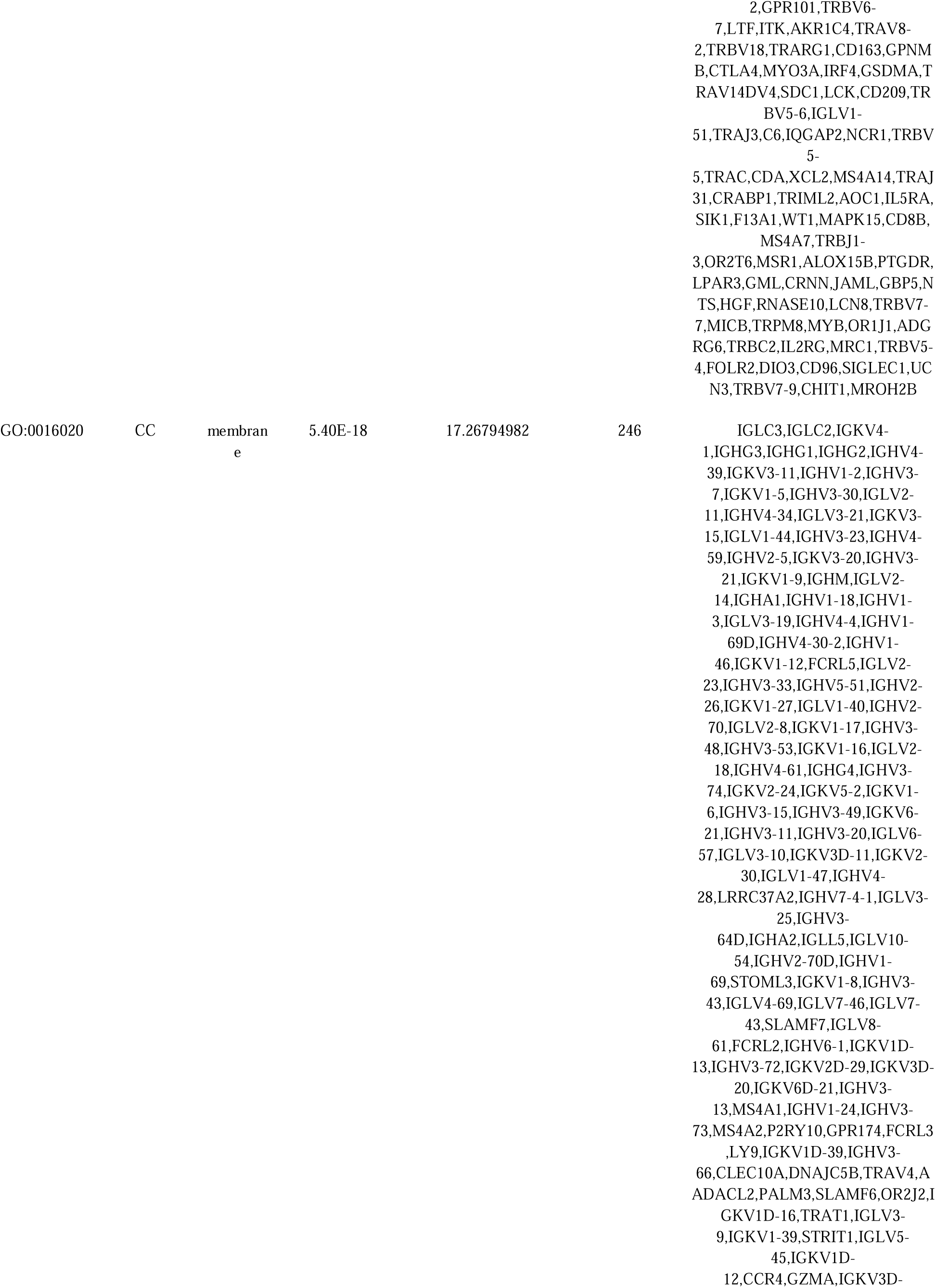
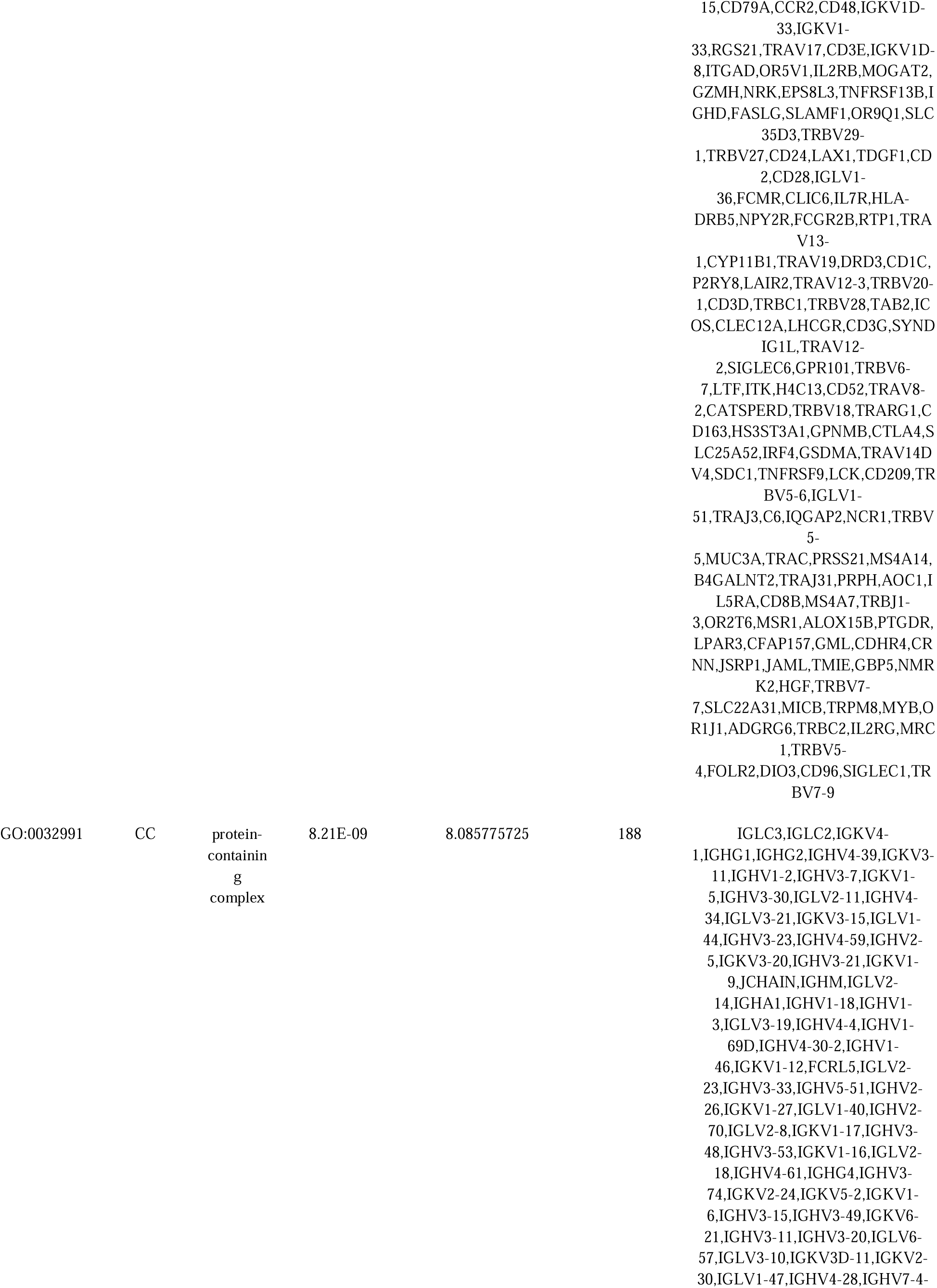
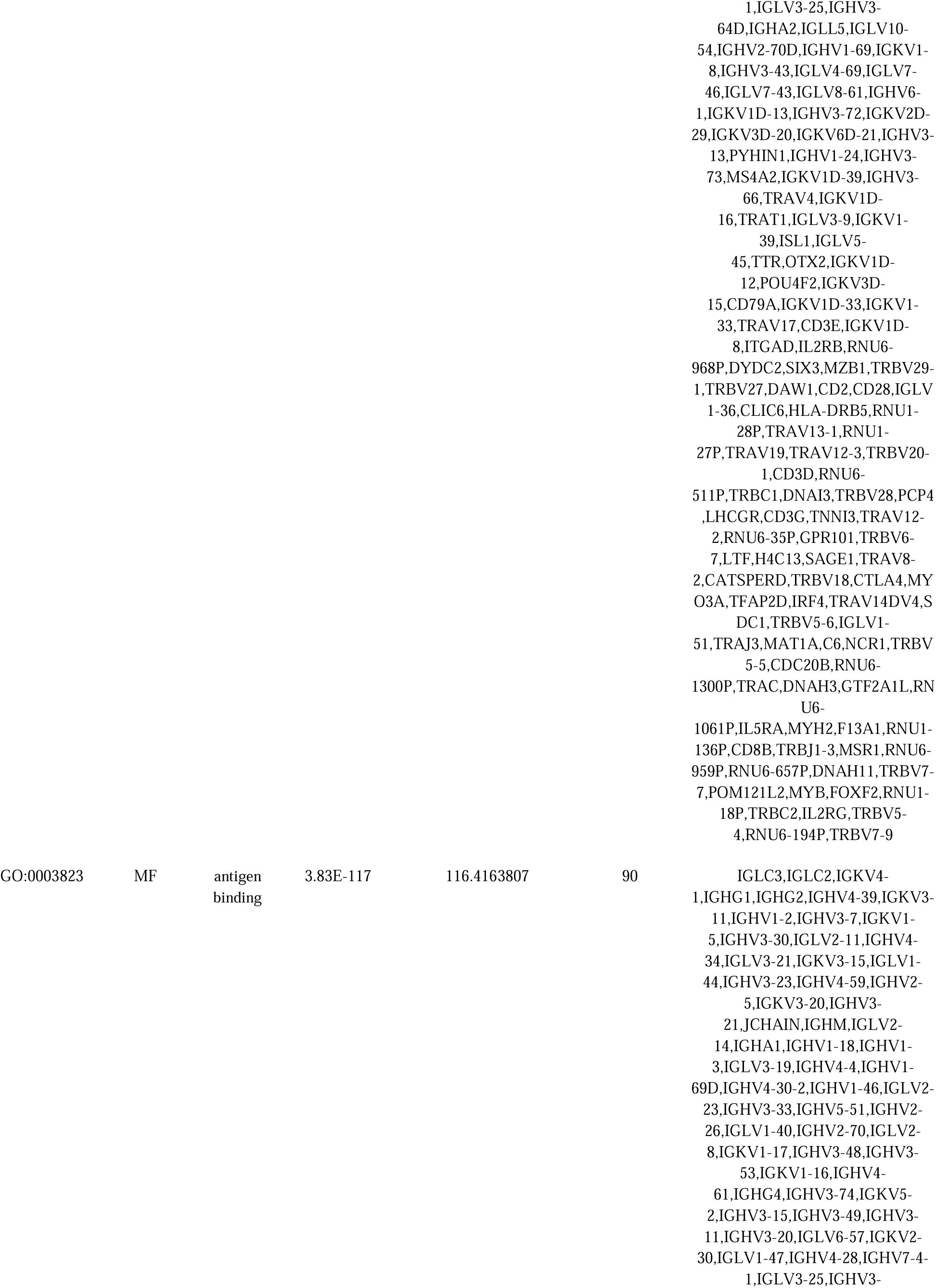
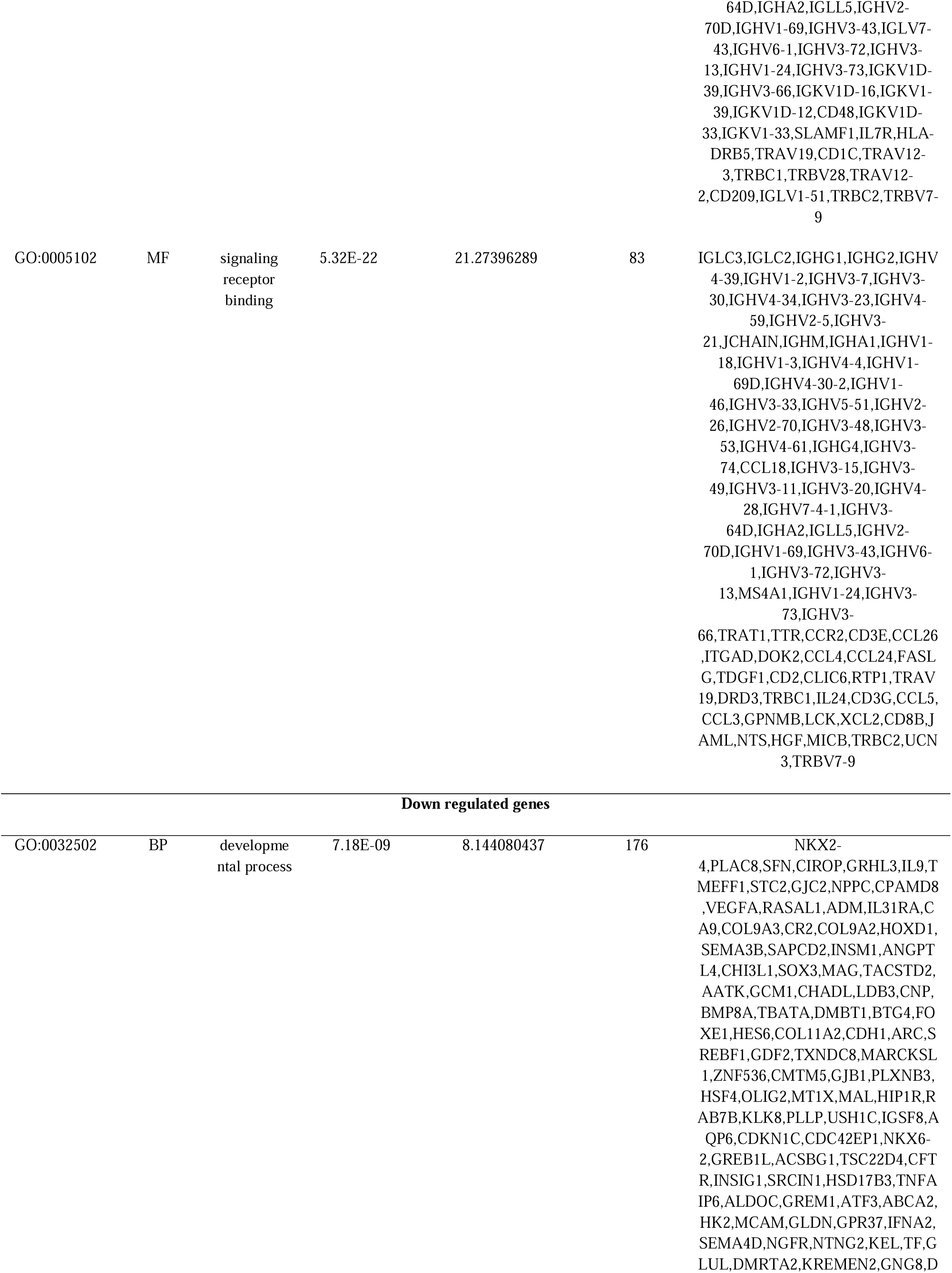
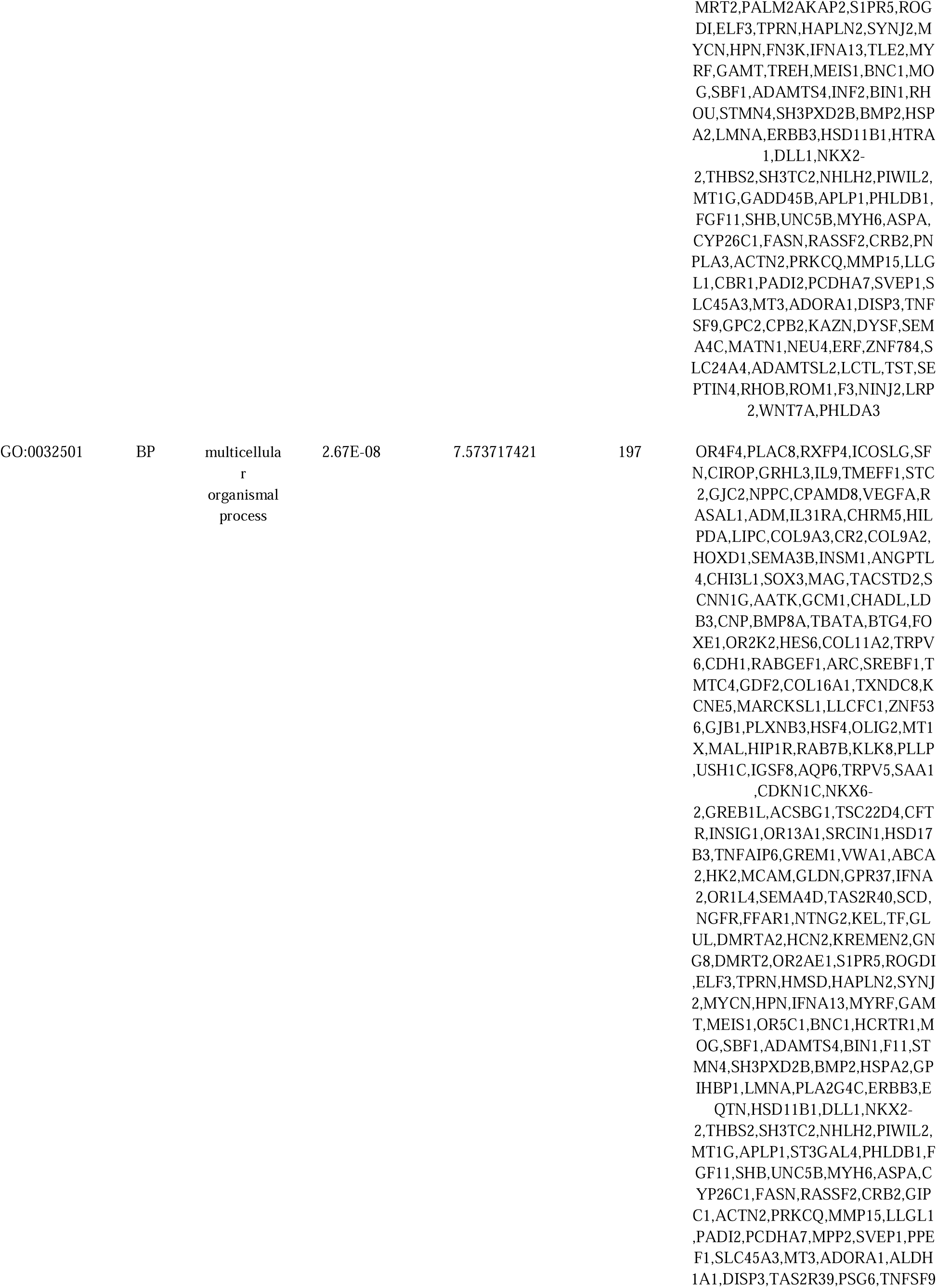
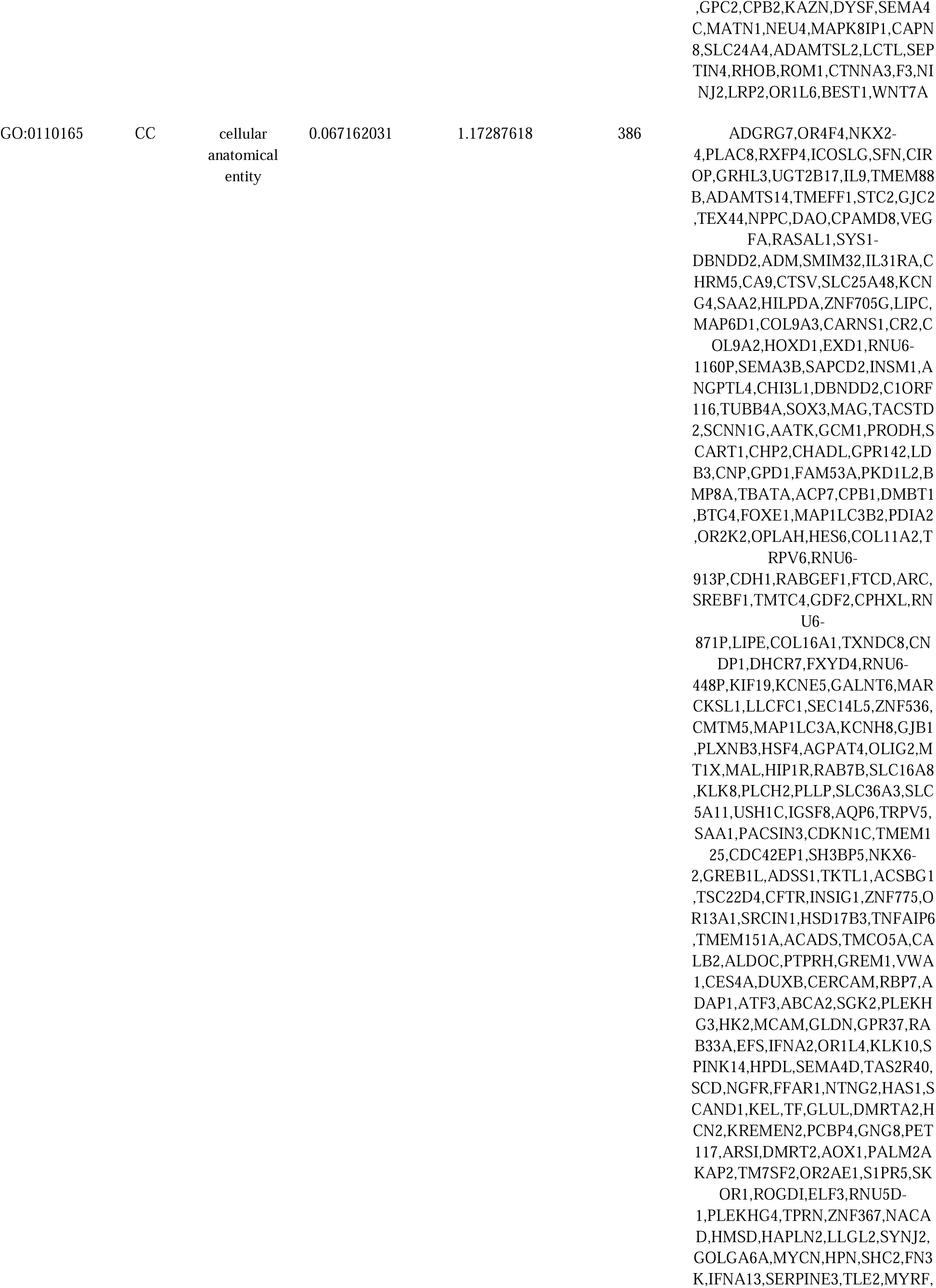
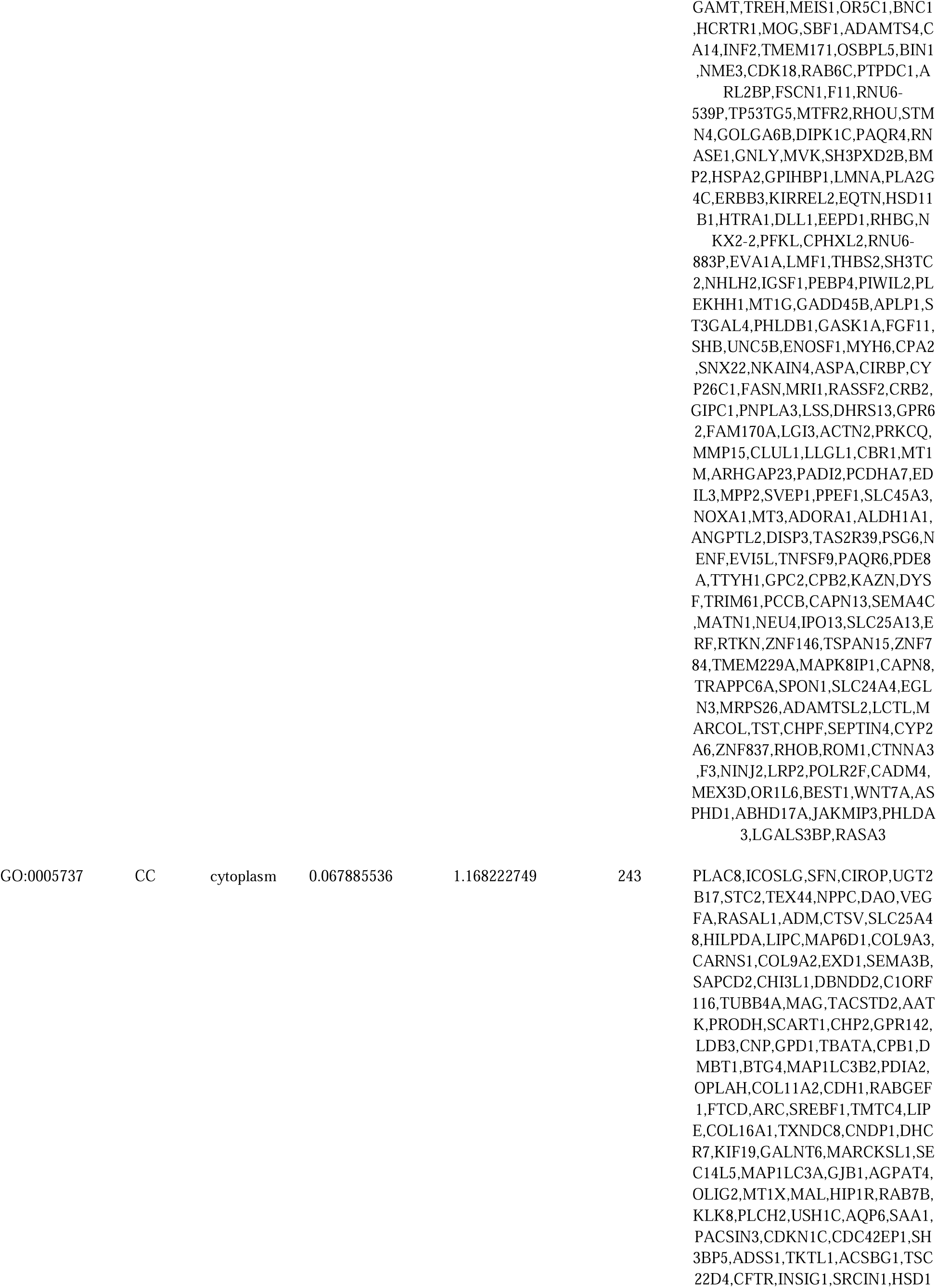
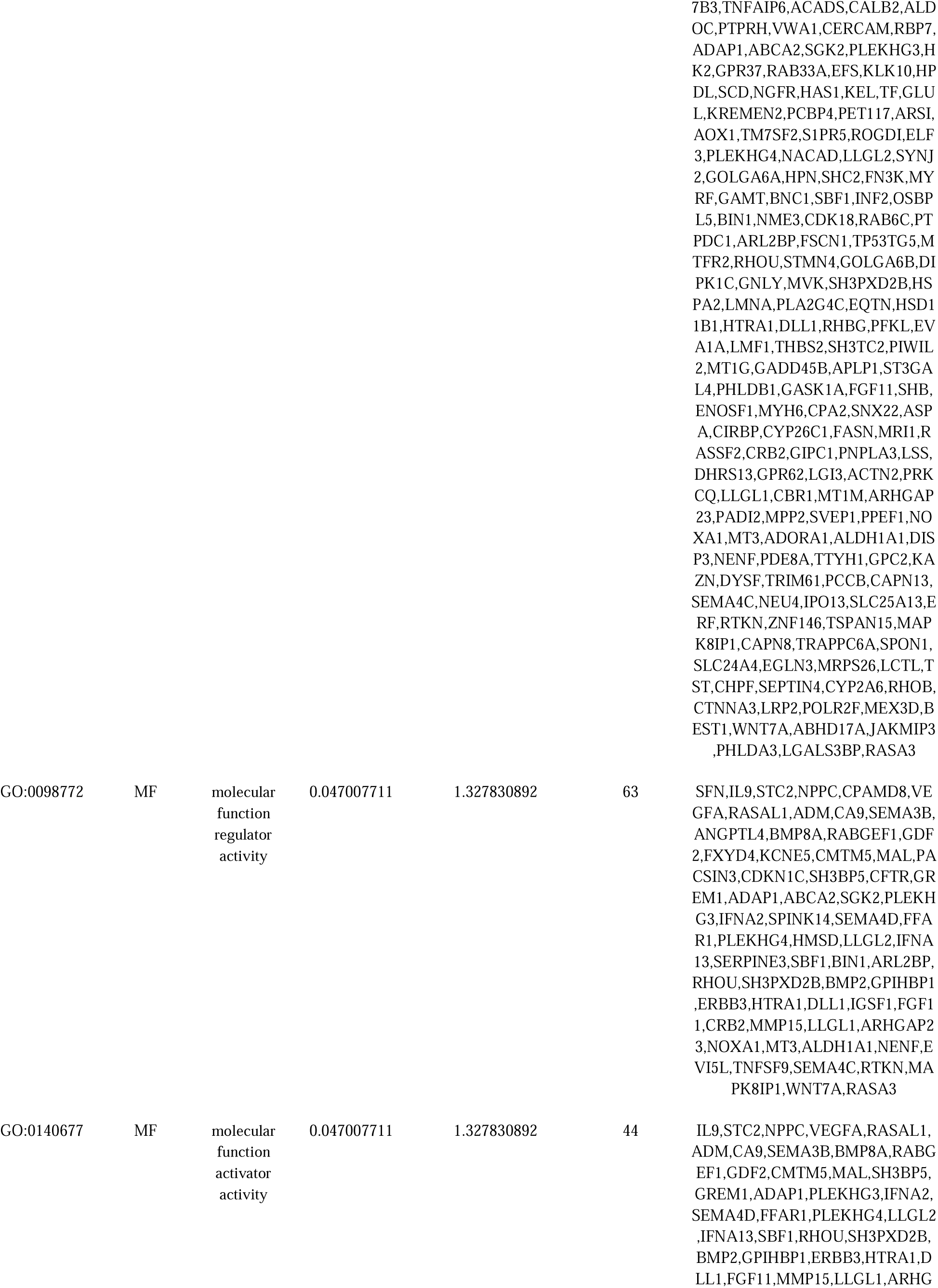
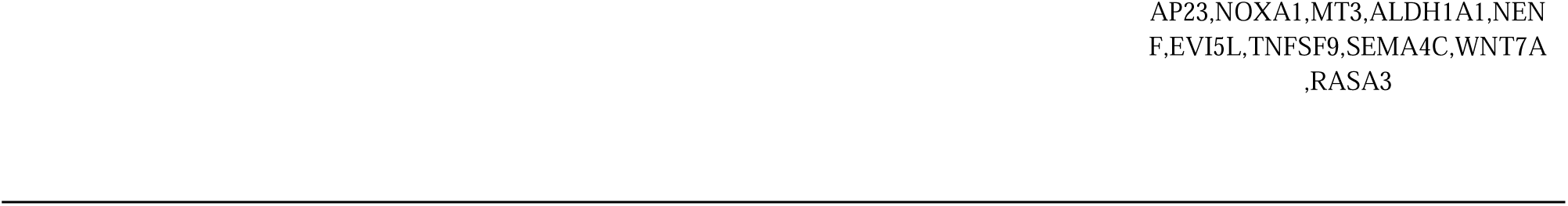
The enriched GO terms of the up and down regulated differentially expressed genes.

**Table 4.**
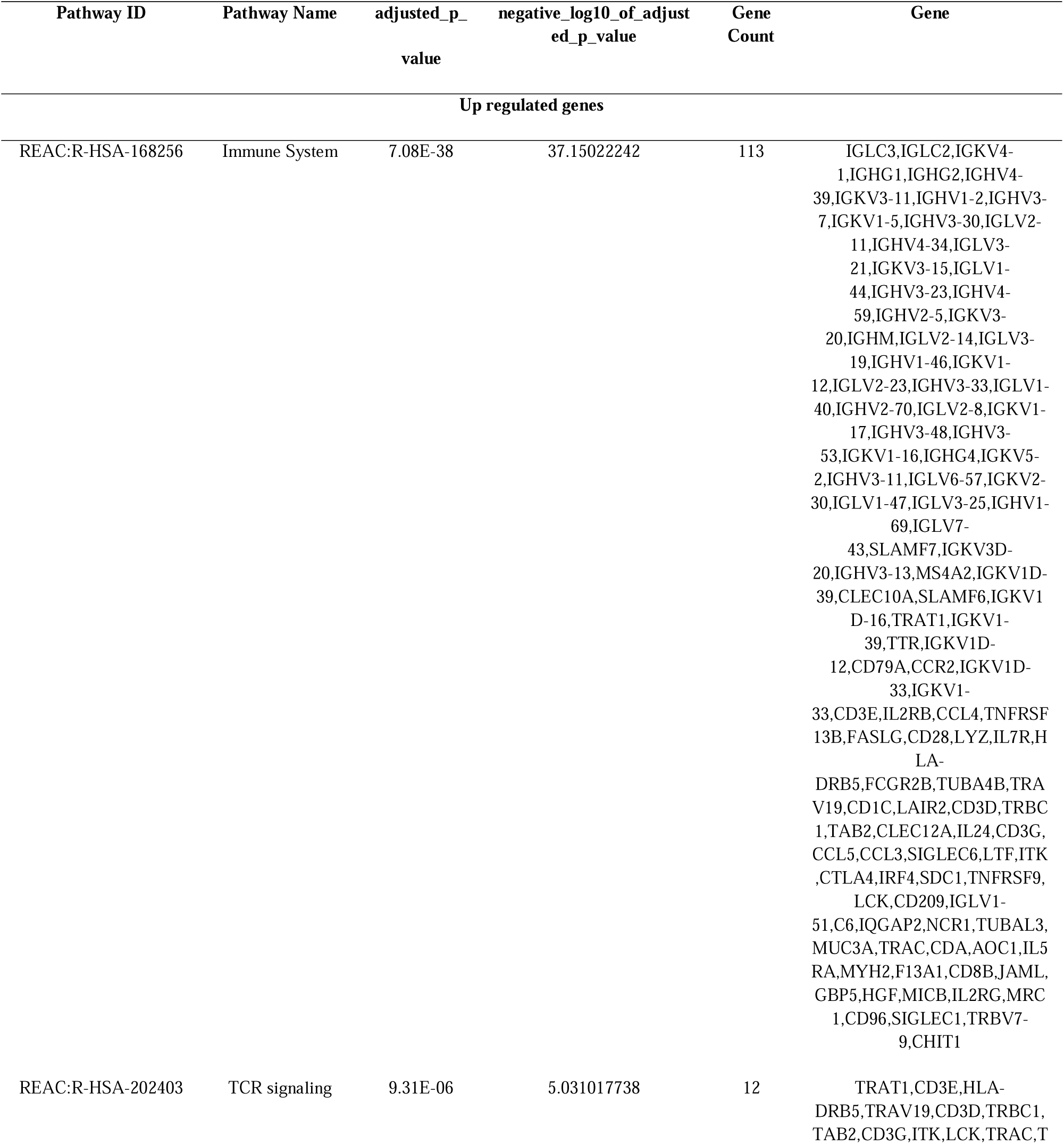
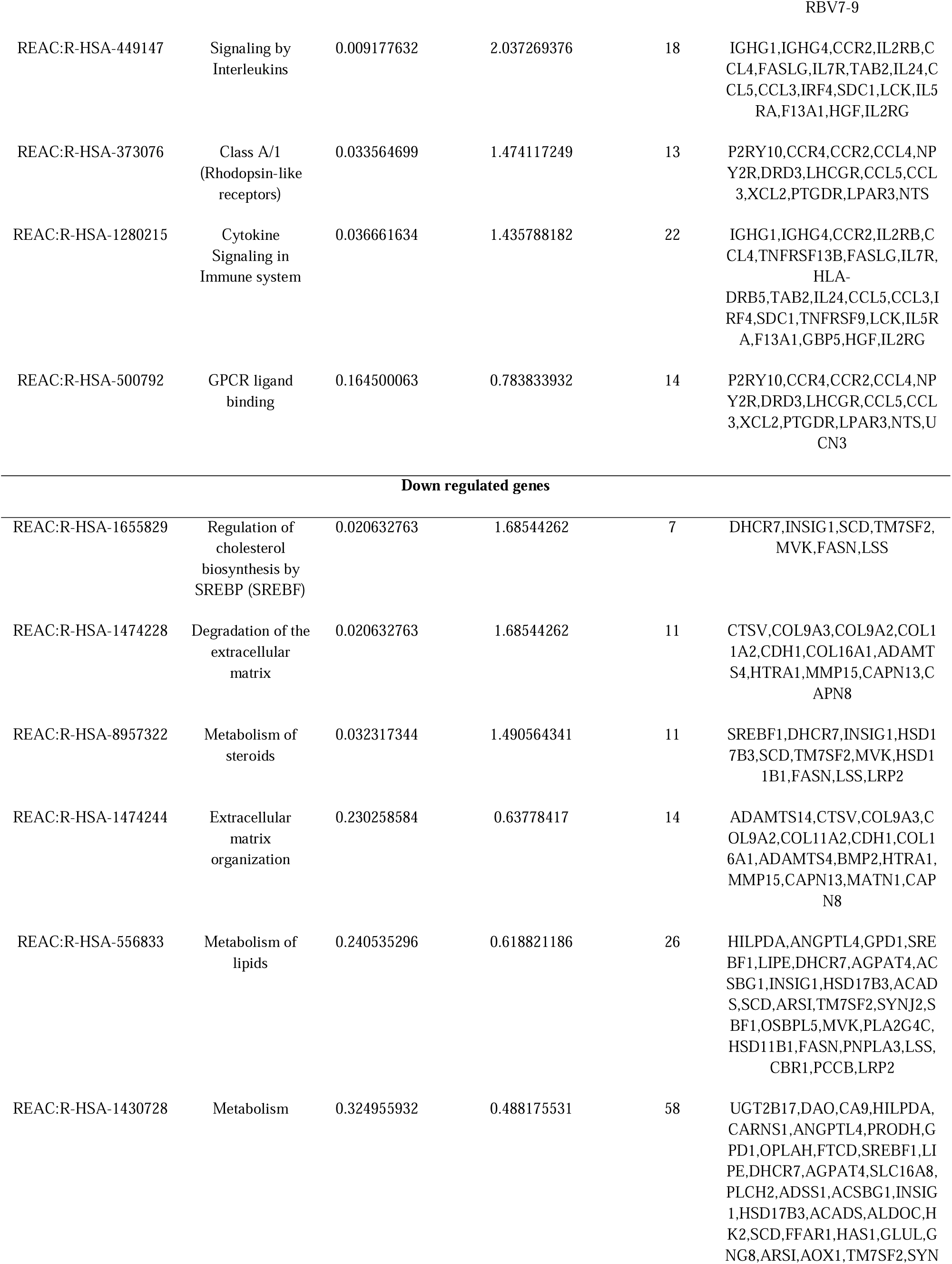
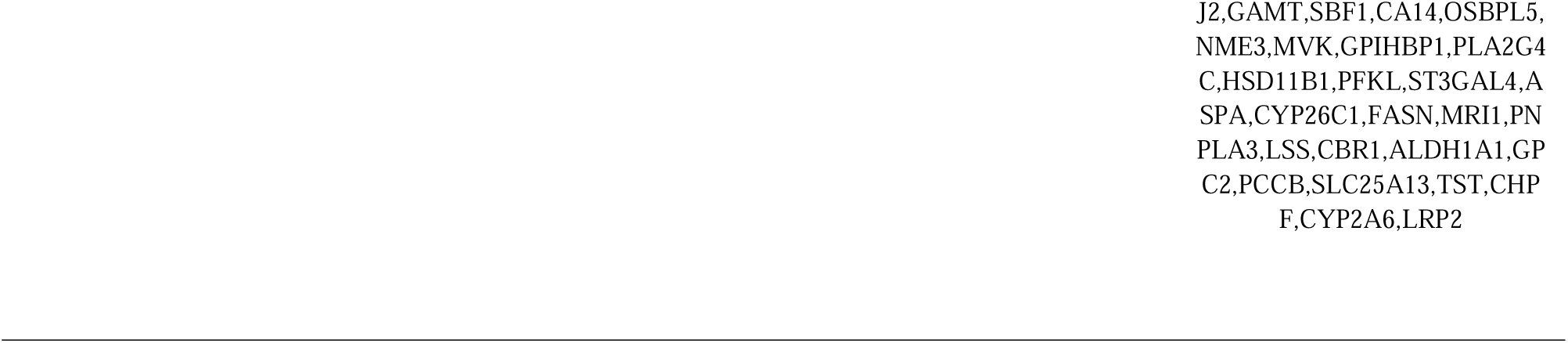
The enriched pathway terms of the up and down regulated differentially expressed genes.

### Construction of the PPI network and module analysis

To explore the interactions between the proteins expressed by the DEGs, we constructed a PPI network using the HiPPIE online PPI database. Cytoscape (version 3.10.1) was used for visualization of PPI network. The PPI network of DEGs contained 4247 nodes (genes) and 6403 edges (interactions) (Fig 6). The hub genes include LCK, PYHIN1, SLAMF1, DOK2, TAB2, CFTR, RHOB, LMNA, EGLN3 and ERBB3 had relatively highest node degree, betweenness, stress and closeness scores (Table 5). Furthermore, PEWCC application results were indicative of two significant modules. Module 1 and module 2 were significant modules in the PPI network. The GO term and REACTOME pathway enrichment analyses of genes involved in two significant modules were analyzed. A total of 47 nodes and 54 edges were included in Module 1 (Fig. 7). Results showed that hub genes in module 1 was mainly enriched in immune system process, immune system and signaling by interleukins (Fig. 8). A total of 28 nodes and 32 edges were included in module 2 (Fig. 9). Results showed that hub genes in module 2 was mainly enriched in cellular anatomical entity (Fig. 10).

**Fig. 6.**
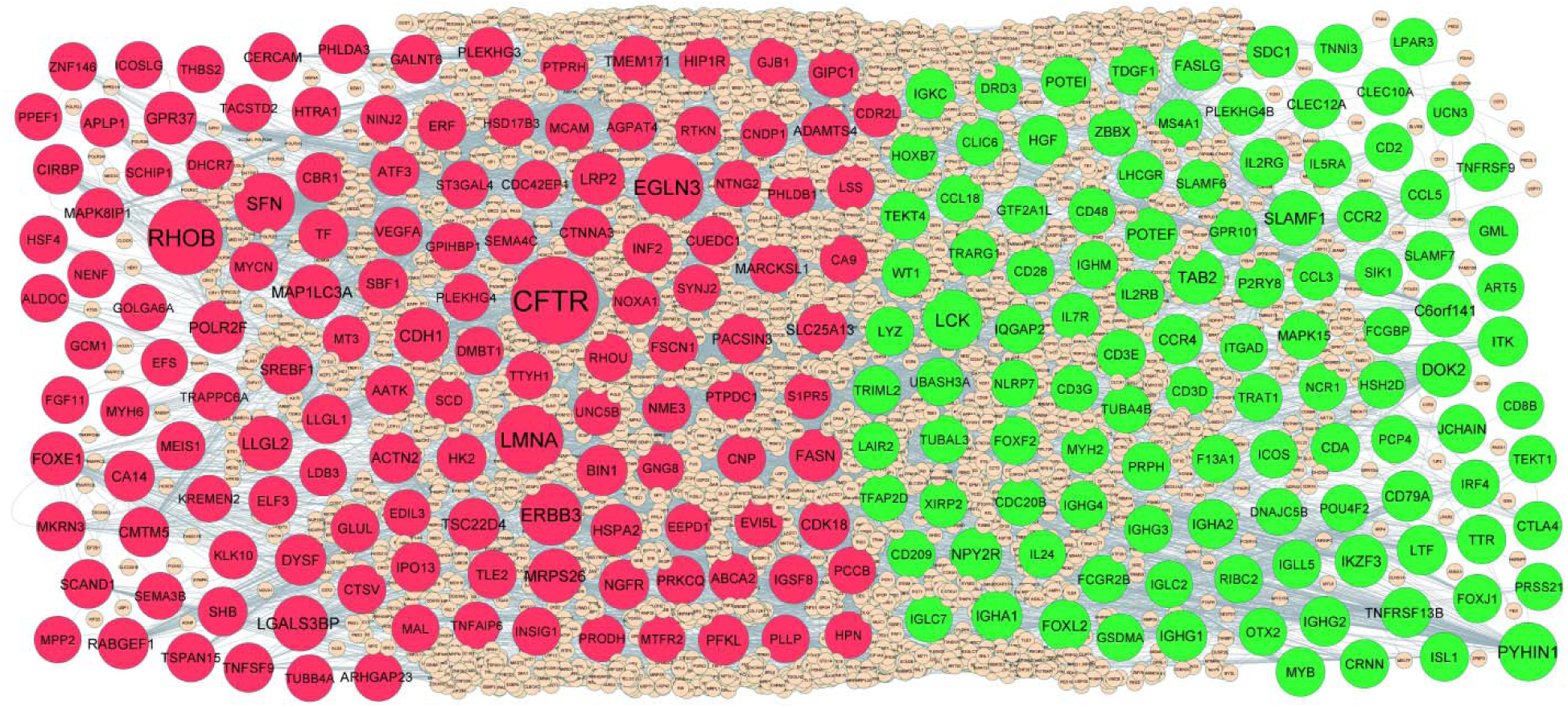
PPI network of DEGs. Up regulated genes are marked in parrot green; down regulated genes are marked in red.

**Fig. 7.**
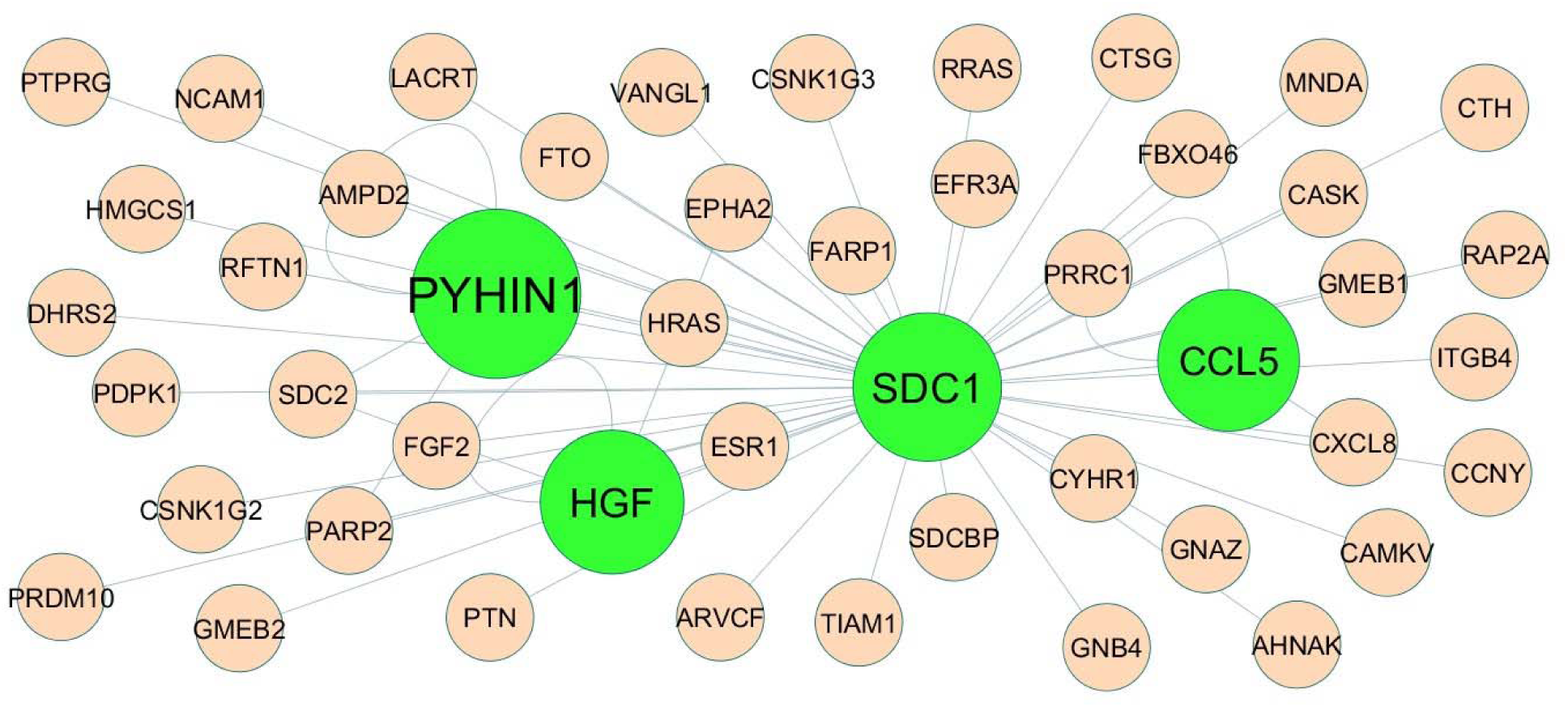
Modules 1 was isolated form PPI of up regulated genes. Module 1 has 47 nodes and 54 edges for up regulated genes. Up regulated genes are marked in green

**Fig. 8.**
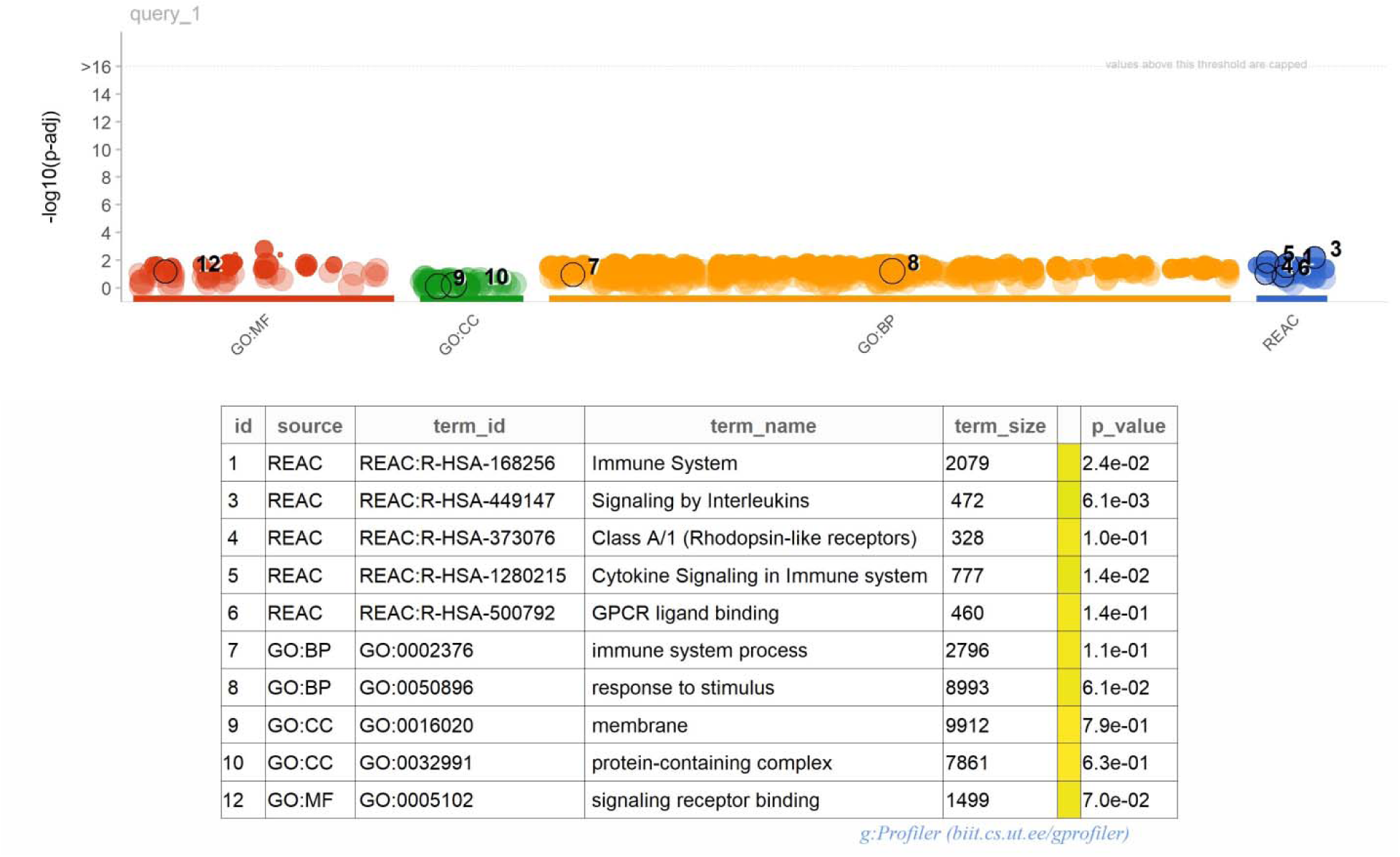
Enrichment analysis for module 1. The size of the circle represents the number of genes involved, and the abscissa represents the frequency of the genes involved in the term total genes.

**Fig. 9.**
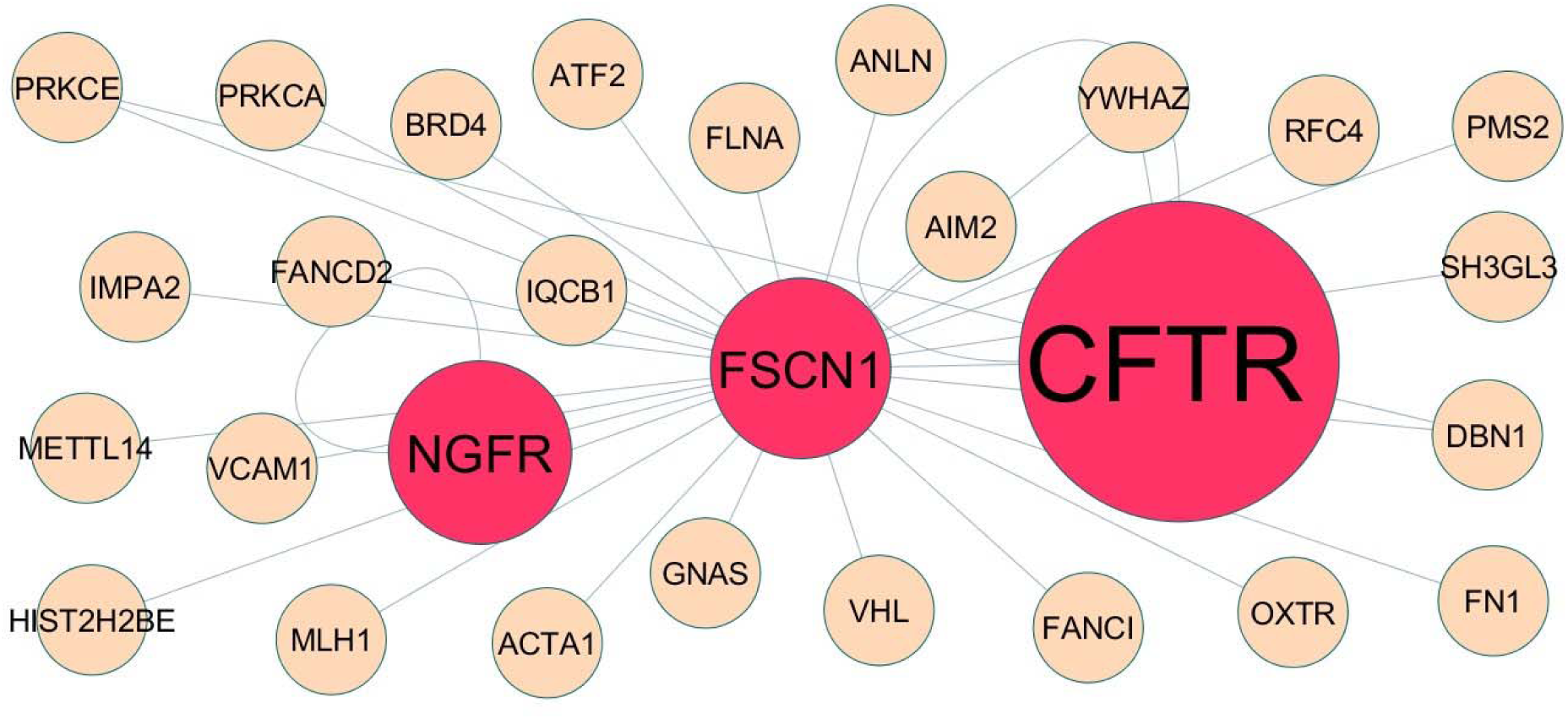
Module 2 was isolated form PPI of up regulated genes. Module 2 has 28 nodes and 32 edges for down regulated genes. Down regulated genes are marked in red

**Fig. 10.**
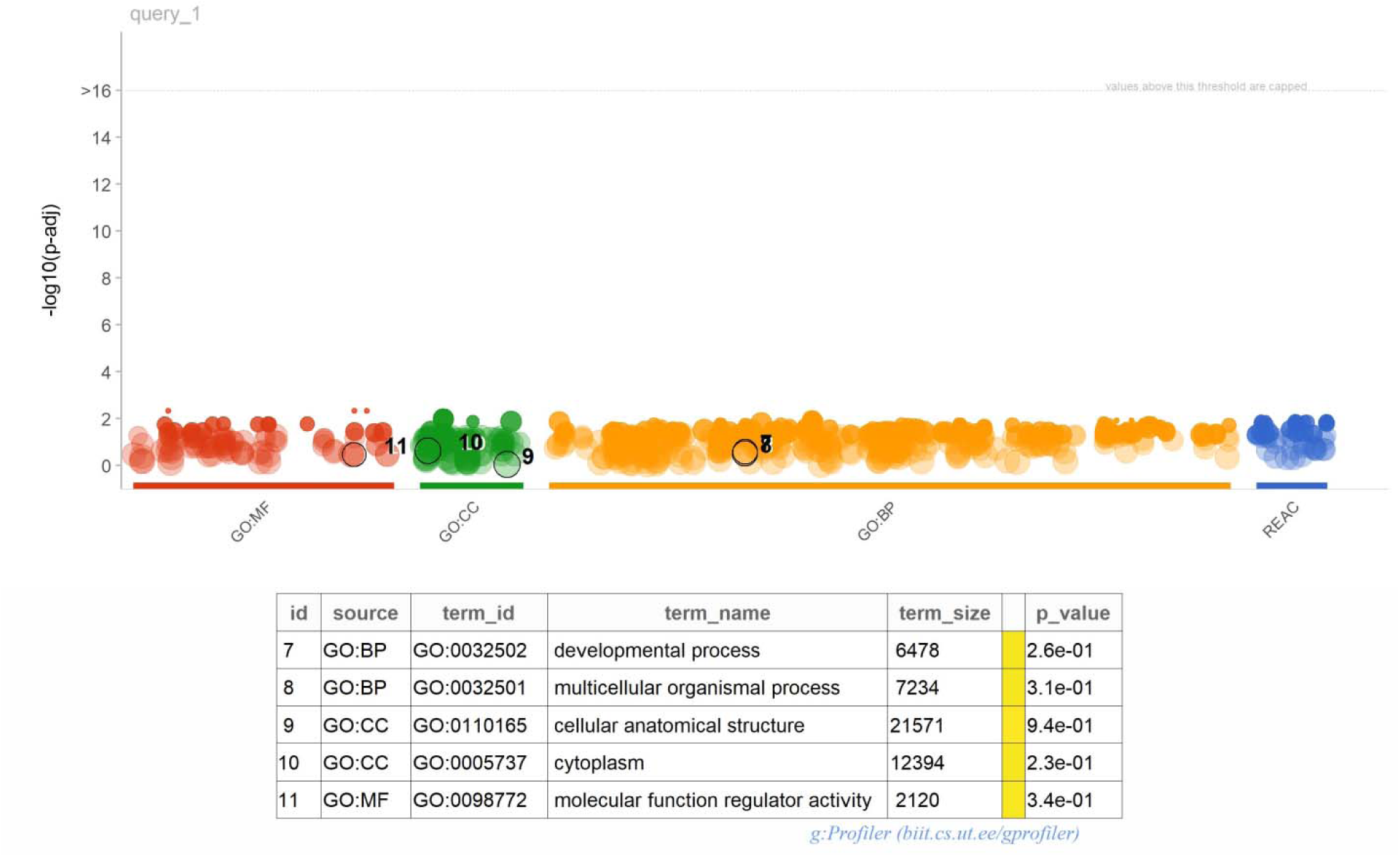
Enrichment analysis for module 2. The size of the circle represents the number of genes involved, and the abscissa represents the frequency of the genes involved in the term total genes.

**Table 5.**
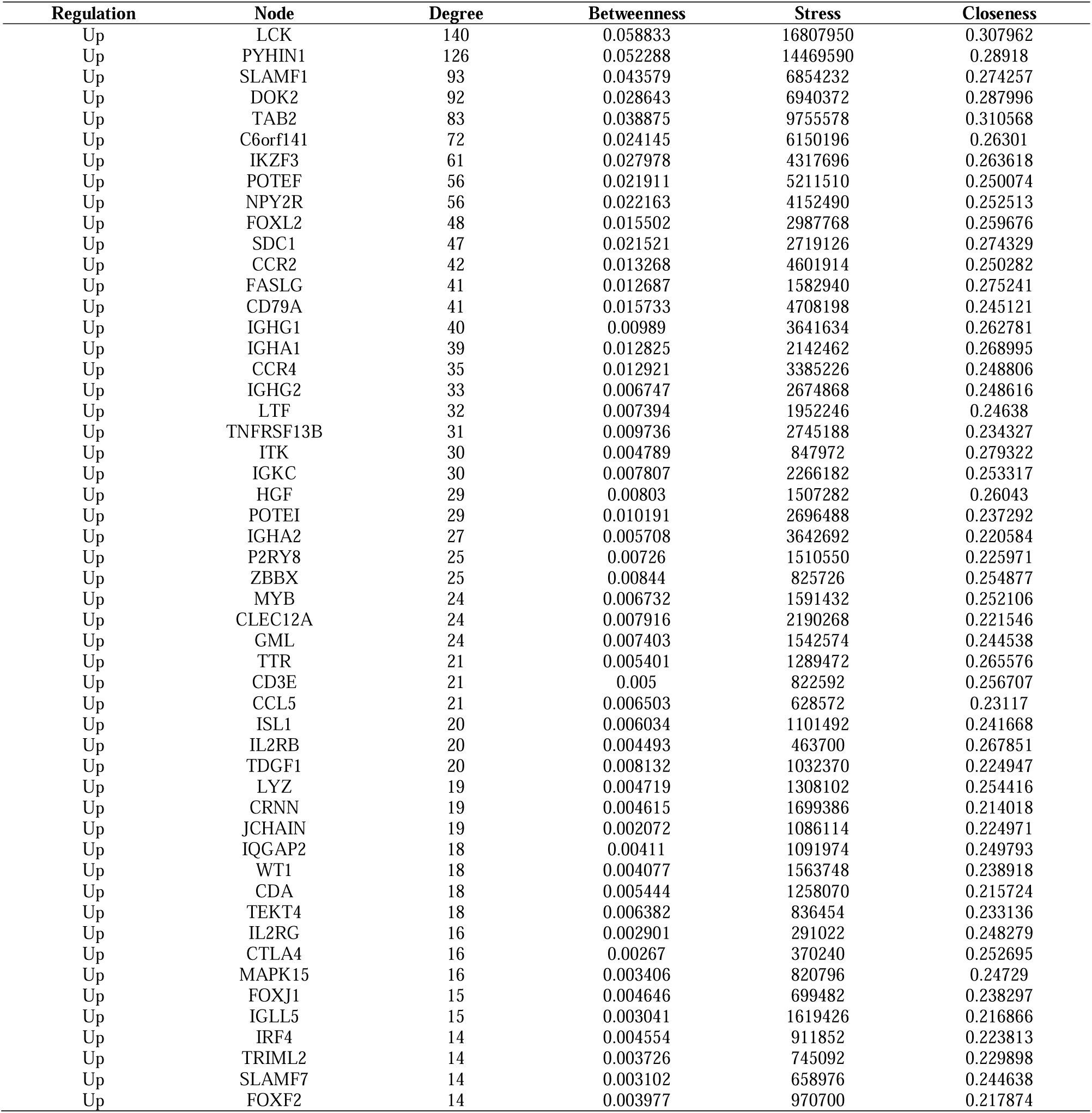
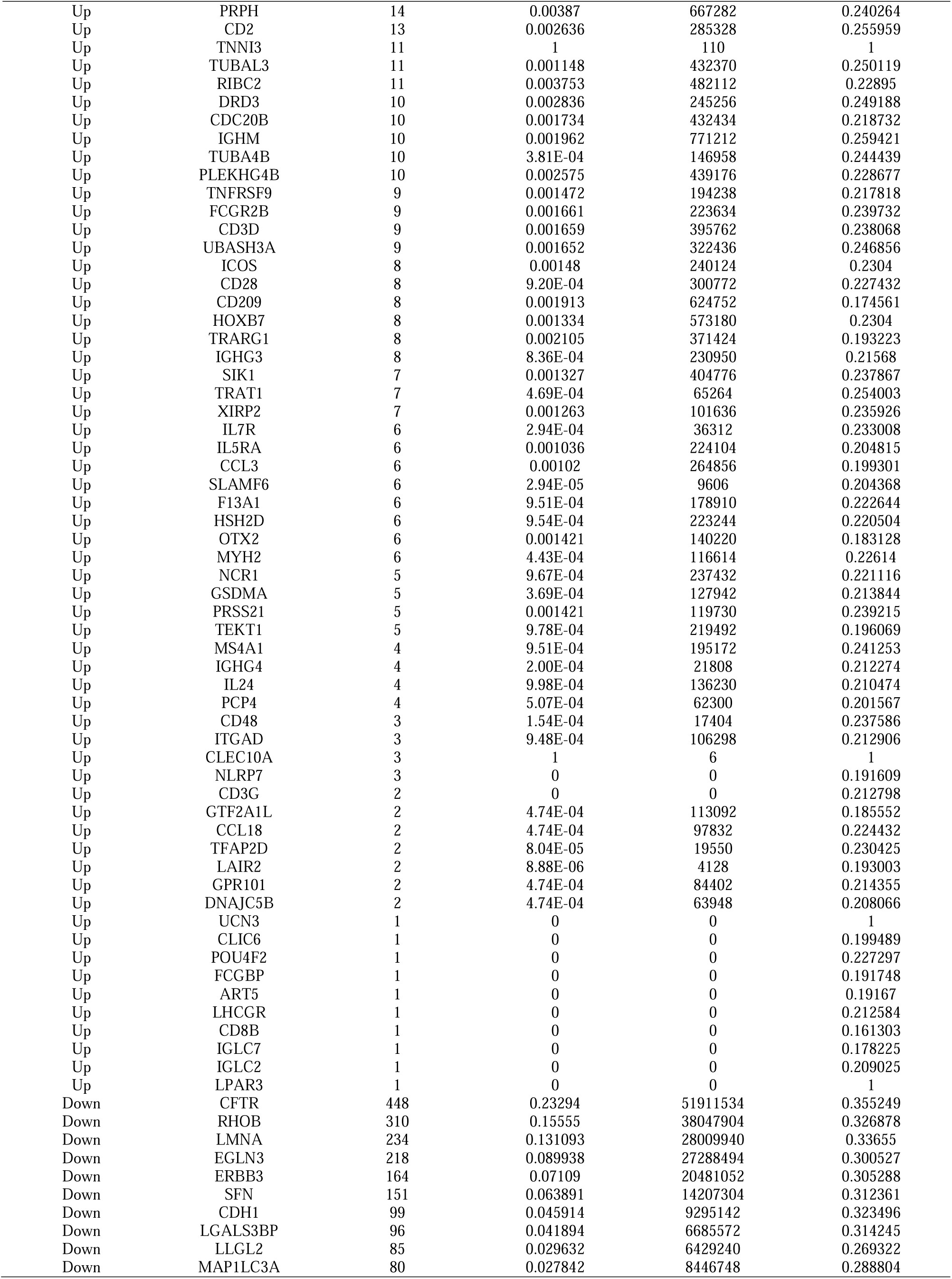
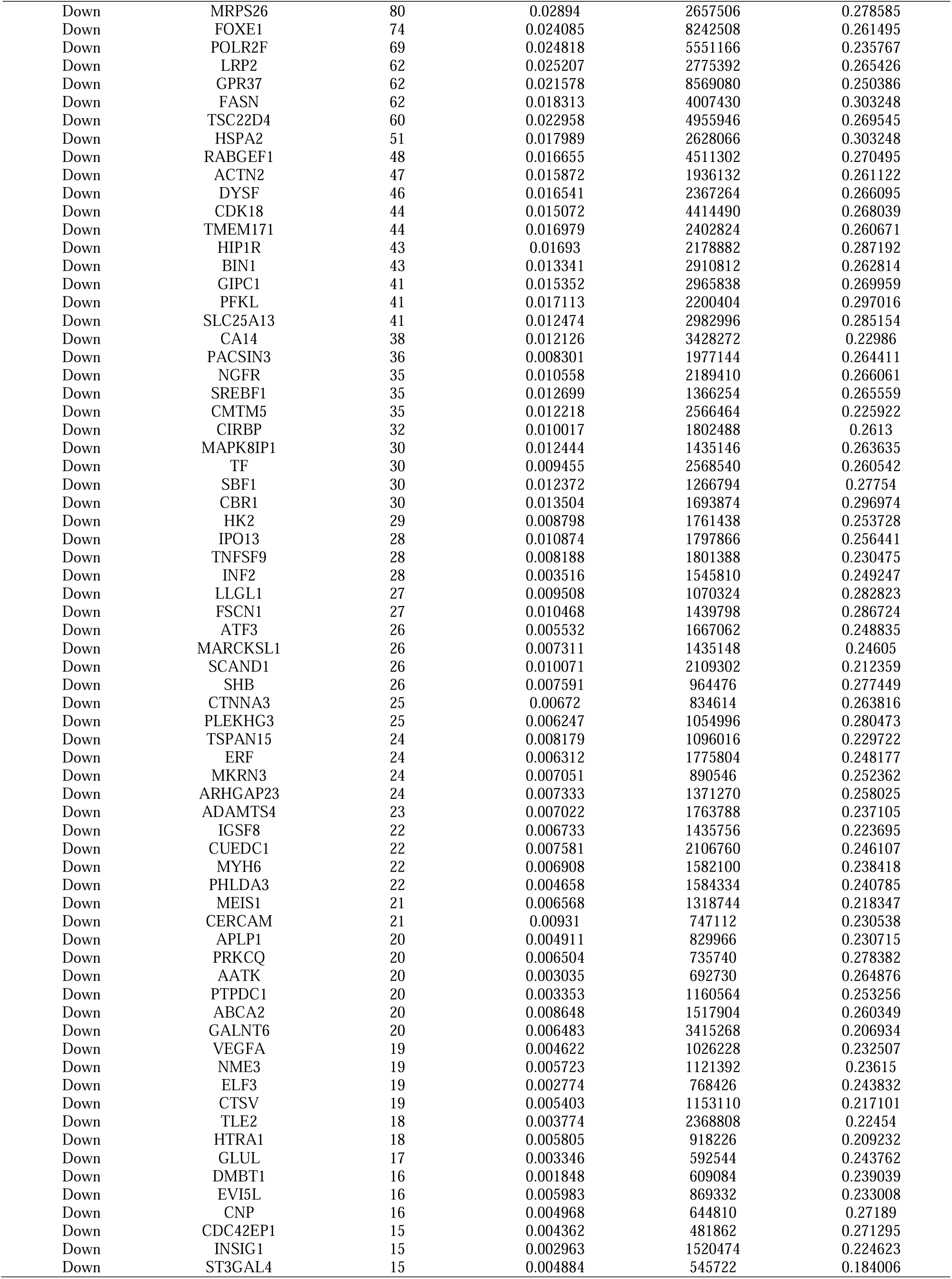
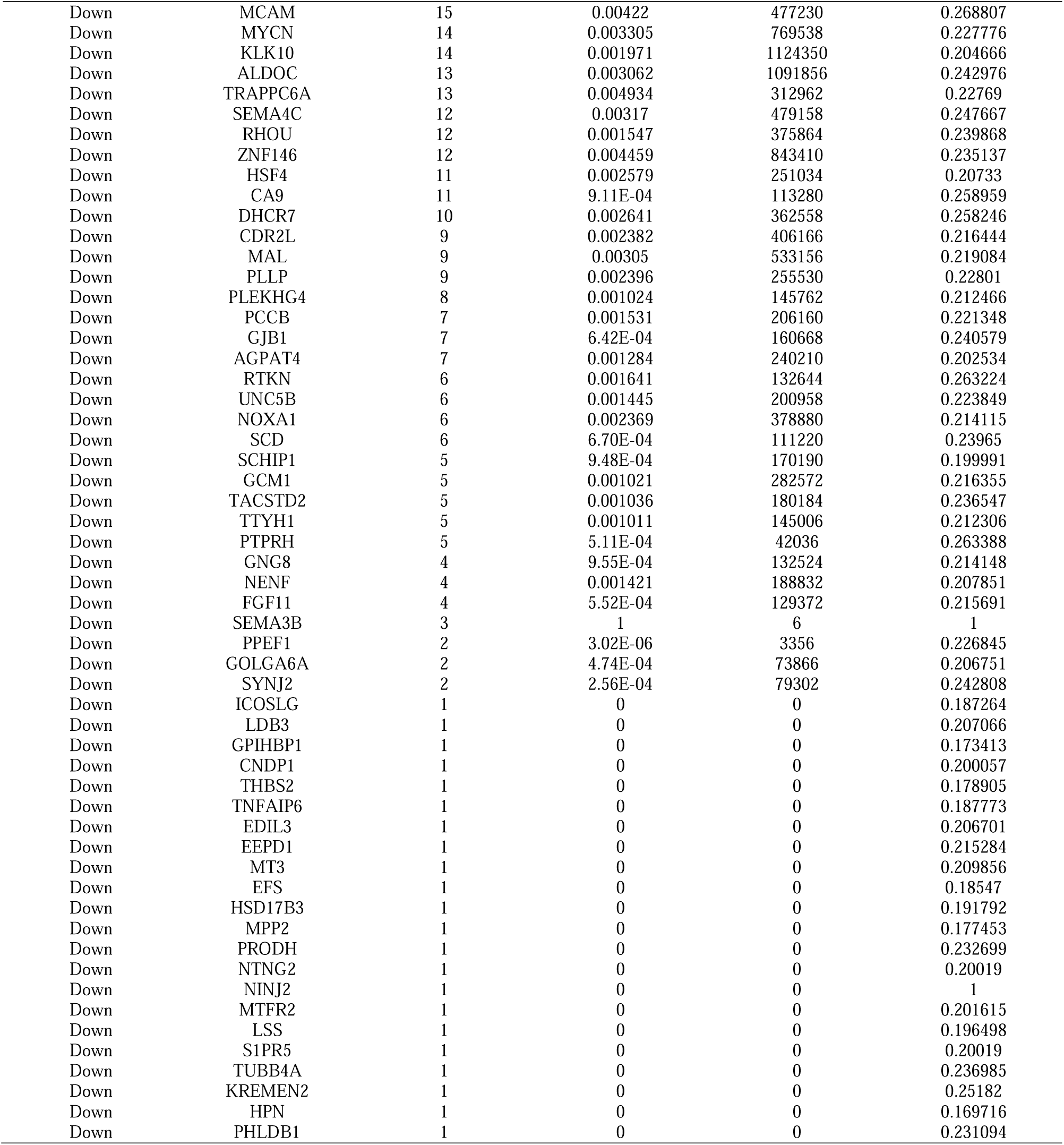
Topology table for up and down regulated genes.

### Construction of the miRNA-hub gene regulatory network

For the hub genes we identified, a miRNA-hub gene regulatory network was constructed including 8791 edges (interaction pairs among the selected hub genes) and 2203 nodes (miRNA: 1973; Hub Gene: 230) (Fig. 11). While IKZF3 was found to be regulated by 147 miRNAs (ex: hsa-mir-6794-3p); TAB2 was found to be regulated by 118 miRNAs (ex: hsa-mir-3689a-3p); C6orf141 was found to be regulated by 59 miRNAs (ex: hsa-mir-4449); SDC1 was found to be regulated by 47 miRNAs (ex: hsa-mir-146a-5p); CCR4 was found to be regulated by 45 miRNAs (ex: hsa-mir-4695-5p); RHOB was found to be regulated by 216 miRNAs (ex: hsa-mir-4651); LGALS3BP was found to be regulated by 99 miRNAs (ex: hsa-mir-548q); SFN was found to be regulated by 91 miRNAs (ex: hsa-mir-8485); CDH1 was found to be regulated by 75 miRNAs (ex: hsa-mir-195- 5p); POLR2F was found to be regulated by 68 miRNAs (ex: hsa-mir-4739) (Table 6).

**Fig. 11.**
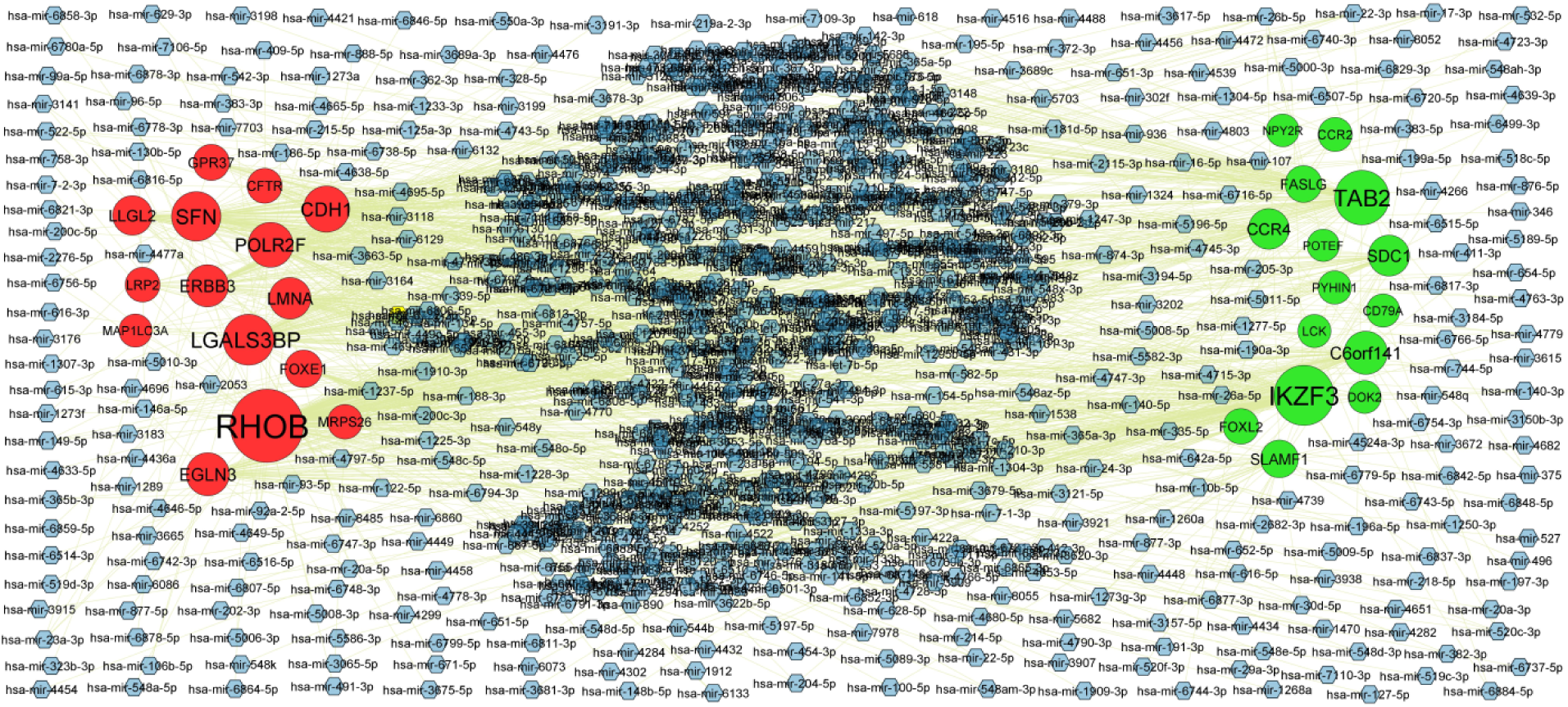
Hub gene - miRNA regulatory network. The blue color diamond nodes represent the key miRNAs; up regulated genes are marked in green; down regulated genes are marked in red.

**Table 6.** MiRNA - hub gene and TF – hub gene topology table.

| Regulation | Hub Genes | Degree | MicroRNA | Regulation | Hub Genes | Degree | TF |
| --- | --- | --- | --- | --- | --- | --- | --- |
| Up | IKZF3 | 147 | hsa-mir-6794-3p | Up | LTF | 15 | BRCA1 |
| Up | TAB2 | 118 | hsa-mir-3689a-3p | Up | CD79A | 10 | HNF4A |
| Up | C6orf141 | 59 | hsa-mir-4449 | Up | LCK | 10 | NFKB1 |
| Up | SDC1 | 47 | hsa-mir-146a-5p | Up | IKZF3 | 9 | TEAD1 |
| Up | CCR4 | 45 | hsa-mir-4695-5p | Up | TAB2 | 8 | ELK4 |
| Up | SLAMF1 | 30 | hsa-mir-4778-3p | Up | PYHIN1 | 8 | JUND |
| Up | FASLG | 25 | hsa-mir-362-3p | Up | DOK2 | 8 | USF2 |
| Up | FOX L2 | 19 | hsa-mir-3915 | Up | POTEF | 7 | STAT3 |
| Up | CCR2 | 11 | hsa-mir-6514-3p | Up | CCR4 | 6 | MEF2A |
| Up | CD79A | 7 | hsa-mir-520f-3p | Up | SLAMF1 | 6 | YY1 |
| Up | PYHIN1 | 7 | hsa-mir-1277-5p | Up | SDC1 | 6 | PPARG |
| Up | LCK | 6 | hsa-mir-520c-3p | Up | FASLG | 5 | ARID3A |
| Up | DOK2 | 5 | hsa-mir-365b-3p | Up | NPY2R | 5 | ELF5 |
| Up | NPY2R | 4 | hsa-mir-302f | Up | FOX L2 | 4 | CEBPB |
| Up | POTEF | 2 | hsa-mir-16-5p | Up | CCR2 | 4 | TP53 |
| Down | RHOB | 216 | hsa-mir-4651 | Down | MAP1LC3A | 14 | TFAP2C |
| Down | LGALS3BP | 99 | hsa-mir-548q | Down | CFTR | 10 | NR2F1 |
| Down | SFN | 91 | hsa-mir-8485 | Down | LLGL2 | 8 | FOXO3 |
| Down | CDH1 | 75 | hsa-mir-195-5p | Down | POLR2F | 8 | NRF1 |
| Down | POLR2F | 68 | hsa-mir-4739 | Down | FOX E1 | 8 | NFYA |
| Down | EGLN3 | 58 | hsa-mir-122-5p | Down | LGALS3BP | 8 | HOXA5 |
| Down | ERBB3 | 53 | hsa-mir-6842-5p | Down | MRPS26 | 8 | SOX17 |
| Down | LMNA | 50 | hsa-mir-744-5p | Down | LRP2 | 8 | E2F6 |
| Down | LLGL2 | 40 | hsa-mir-200c-3p | Down | CDH1 | 7 | ESR1 |
| Down | FOX E1 | 29 | hsa-mir-4723-3p | Down | SFN | 6 | FOS |
| Down | GPR37 | 22 | hsa-mir-186-5p | Down | RHOB | 5 | RELA |
| Down | CFTR | 16 | hsa-mir-154-5p | Down | EGLN3 | 4 | POU2F2 |
| Down | LRP2 | 15 | hsa-mir-196a-5p | Down | ERBB3 | 4 | GATA2 |
| Down | MRPS26 | 14 | hsa-mir-3065-5p | Down | LMNA | 4 | JUN |
| Down | MAP1LC3A | 6 | hsa-mir-22-3p | Down | GPR37 | 4 | FOX L1 |

### Construction of the TF-hub gene regulatory network

For the hub genes we identified, a TF-hub gene regulatory network was constructed including 1809 edges (interaction pairs among the selected hub genes) and 312 nodes (TF: 80; Hub Gene: 232) (Fig. 12). While LTF was found to be regulated by 15 TFs (ex: BRCA1); CD79A was found to be regulated by 10 TFs (ex: HNF4A); LCK was found to be regulated by 10 TFs (ex: NFKB1); IKZF3 was found to be regulated by 9 TFs (ex: TEAD1); TAB2 was found to be regulated by 8 TFs (ex: ELK4); MAP1LC3A was found to be regulated by 14 TFs (ex: TFAP2C); CFTR was found to be regulated by 10 TFs (ex: NR2F1); LLGL2 was found to be regulated by 8 TFs (ex: FOXO3); POLR2F was found to be regulated by 8 TFs (ex: NRF1); FOXE1 was found to be regulated by 8 TFs (ex: NFYA) (Table 6).

**Fig. 12.**
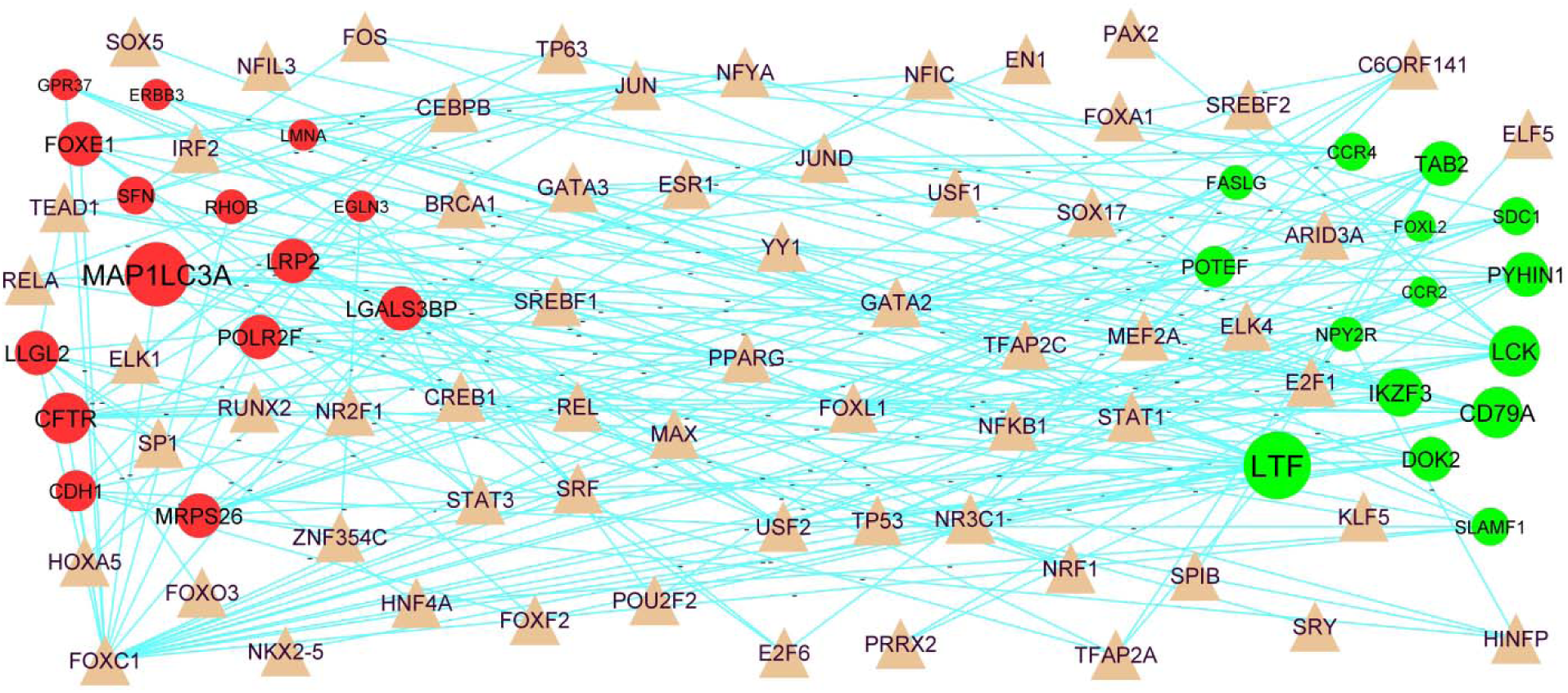
Hub gene - TF regulatory network. The brown color triangle nodes represent the key TFs; up regulated genes are marked in green; down regulated genes are marked in red.

### Construction of the drug-hub gene interaction network

Drug-hub gene interaction network was constructed. For up regulated genes or targets (Fig. 13 and Table 7), 20 drugs (ex: Palivizumab) targeted FCGR2B, 15 drugs (ex: Cu-Bicyclam) targeted LYZ, 14 drugs (ex: Nintedanib) targeted LCK, 7 drugs (ex: Goserelin) targeted LHCGR and 5 drugs (ex: Ofatumumab) targeted MS4A1, while down regulated genes or targets (Fig. 14 and Table 6), 8 drugs (ex: Lumacaftor) targeted CFTR, 6 drugs (ex: Zonisamide) targeted CA9, 5 drugs (ex: Bentiromide) targeted HPN, 4 drugs (ex: Tamoxifen) targeted PRKCQ and 4 drugs (ex: Urokinase) targeted LRP2.

**Fig. 13.**
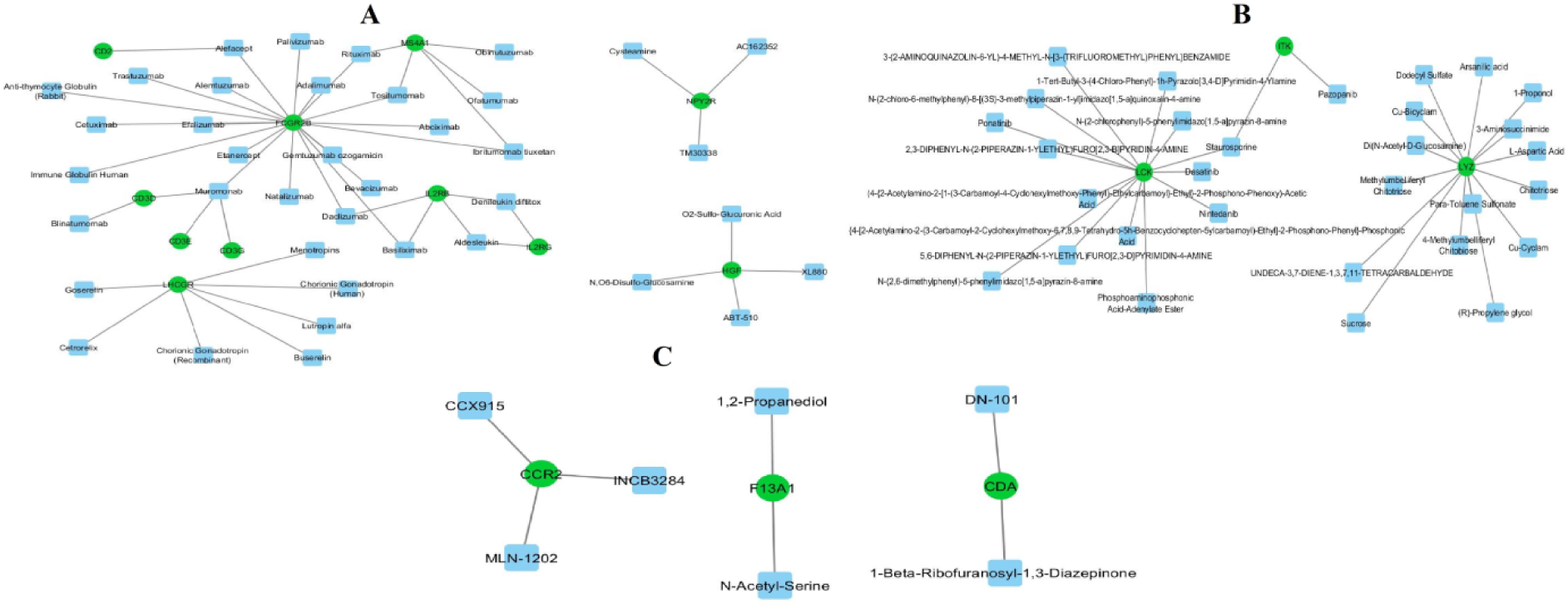
Drug-hub gene interaction network. The blue color rectangle nodes represent the drug molecule; up regulated genes are marked in green

**Fig. 14.**
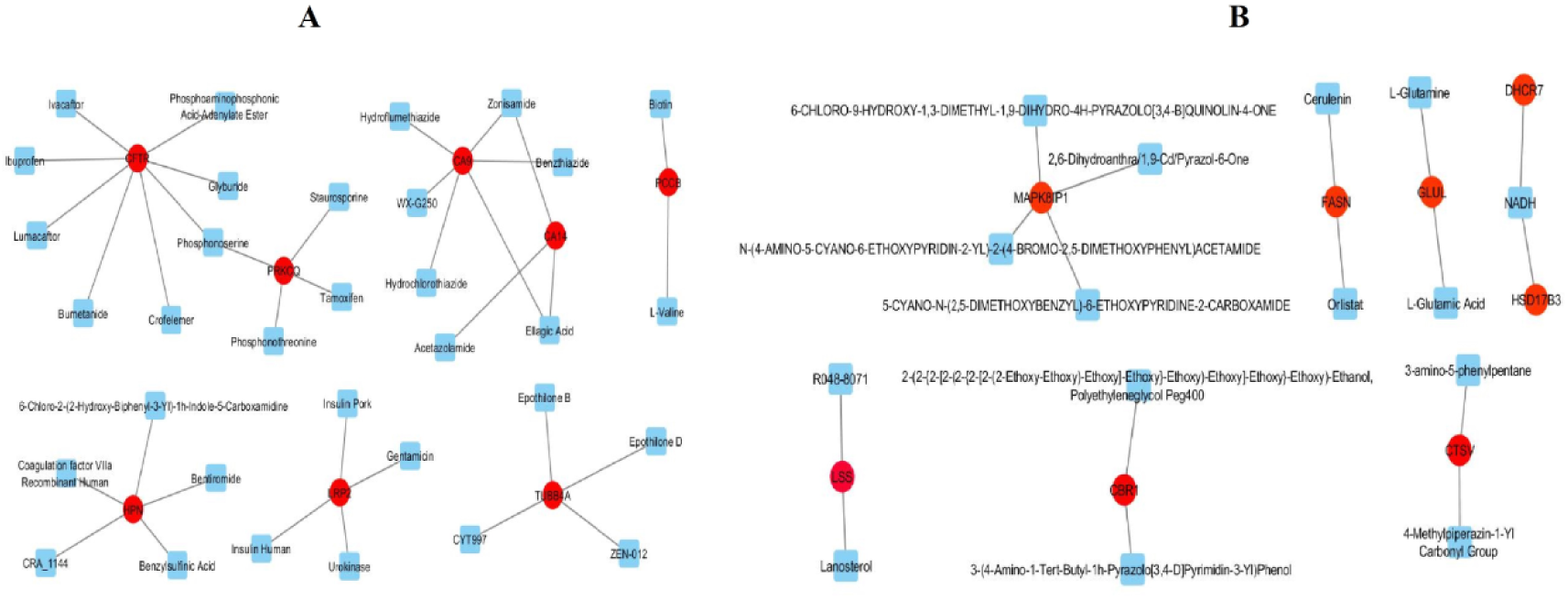
Drug-hub gene interaction network. The blue color rectangle nodes represent the key drug molecules; down regulated genes are marked in red

**Table 7.**
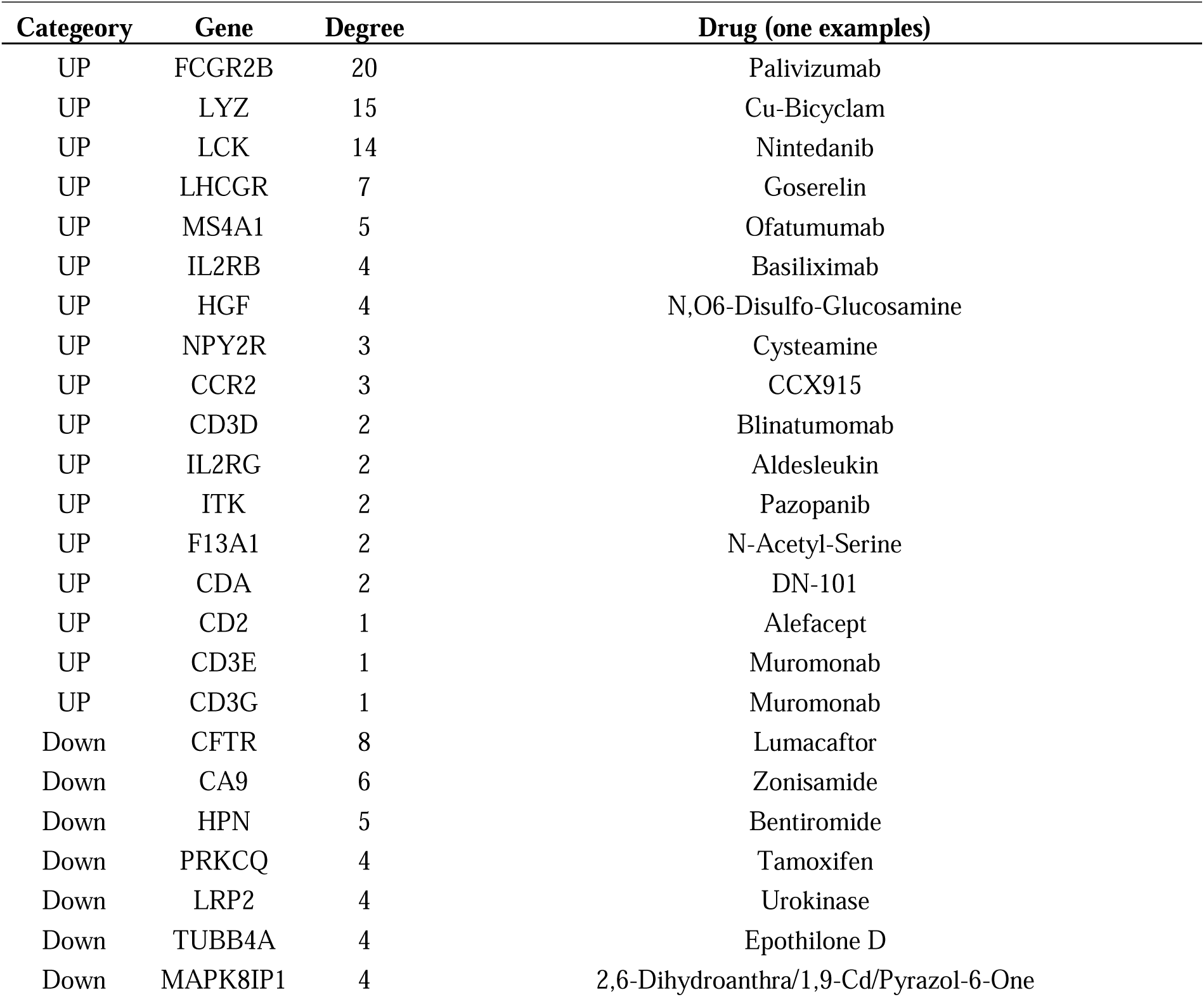
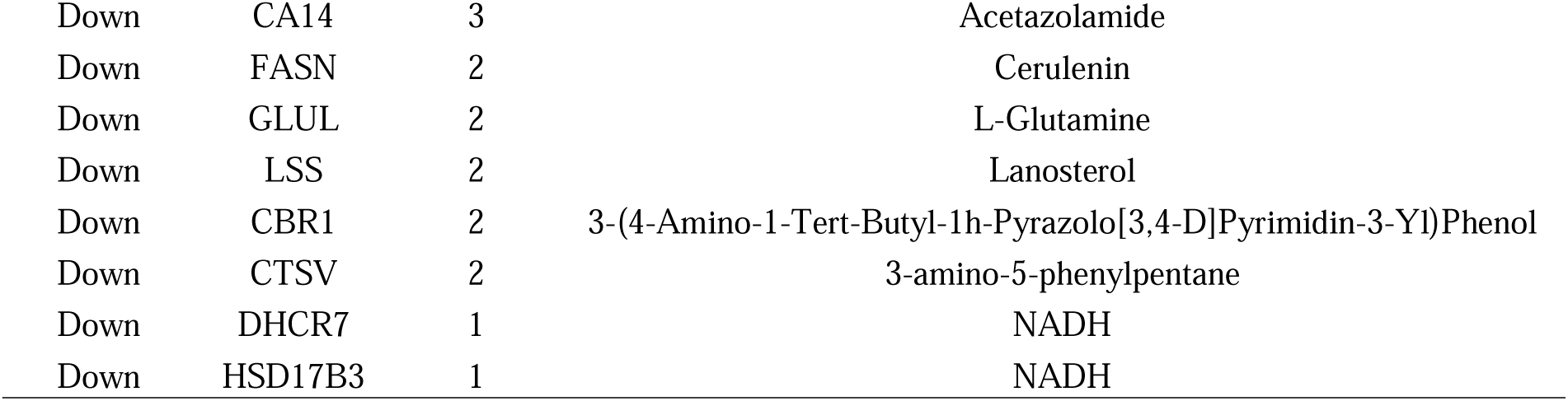
Drug- hub gene topology table.

### Receiver operating characteristic curve (ROC) analysis

To access the potential prediction function of hub genes in MS, ROC curves were established to evaluate the diagnostic specificity and sensitivity of hub genes using the “pROC” R bioconductor package. The AUC was calculated for each hub gene. The results are as follows: LCK (AUC = 0.931), PYHIN1 (AUC = 0.902), SLAMF1 (AUC = 0.912), DOK2 (AUC = 0.933), TAB2 (AUC = 0.916), CFTR (AUC = 0.920), RHOB (AUC = 0.935), LMNA (AUC = 0.921), EGLN3 (AUC = 0.927) and ERBB3 (AUC = 0.925) (Fig. 15). All hub genes exhibited high diagnostic value for MS.

**Fig. 15.**
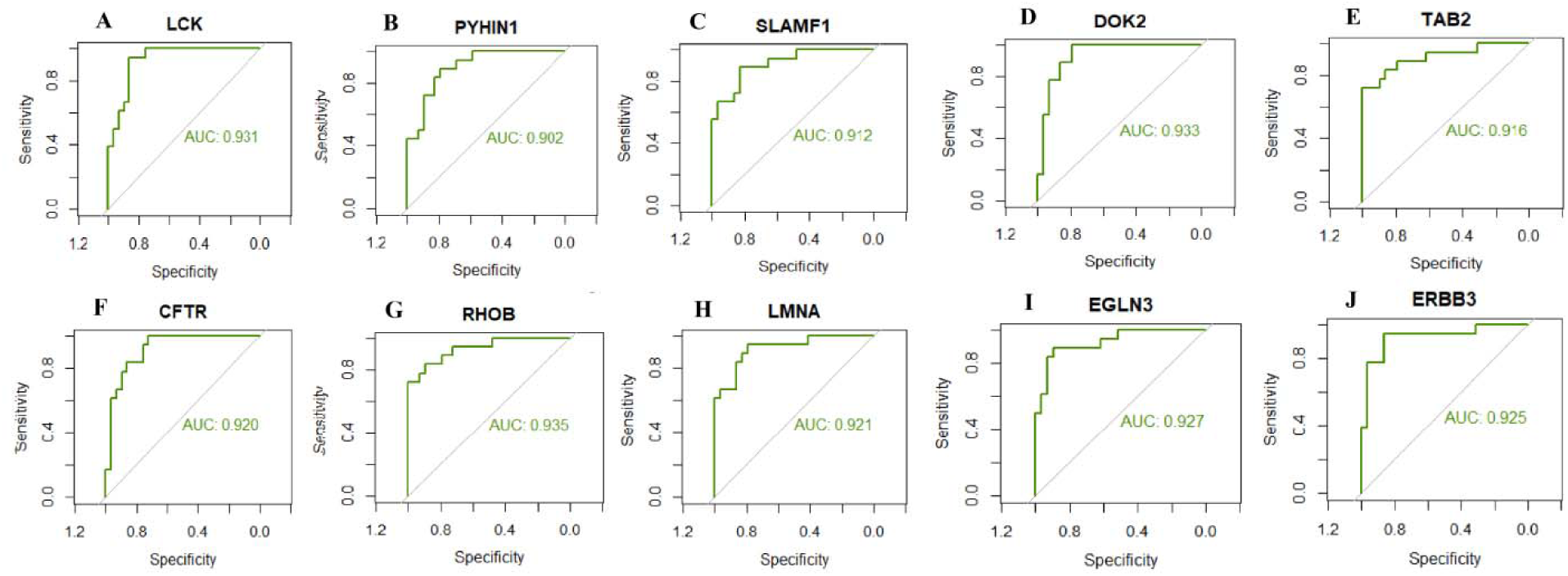
ROC curve analyses of hub genes. A) LCK B) PYHIN1 C) SLAMF1 D) DOK2 E) TAB2 F) CFTR G) RHOB H) LMNA I) EGLN3 J) ERBB3

### *Insilico* molecular docking studies

The ligands were obtained from network analysis and used to find the best suitable fit for their respective receptors and the data were described belowMolecular docking study of selected ligands were performed well, and results analysis was carried out, it was indicative of all molecule was bound well, Nintedanib with LCK (Pdb id: 2go8), Glyburide with CFTR (Pbd id: 2pzg), Glyburide with CFTR (Pdb id: 2bbs), Caffeine with CA9 (Pdb id: 5dvx), Goserelin with LHCGR (1qfw), Caffeine with CA14 (Pdb id: 4lu3), Glucosamine 6-sulfate with HGF (Pdb id: 3hms), Phosphoserine with PRKCQ (Pdb id: 1xjd), receptor with binding affinity of −9.7, −8.5, −7.5, −6.7, −6.6, −5.6, and −5.2 Kcal/mol respectively (Table 8). Among all the screened compounds, Ninatanib and Glyburide exhibited the highest negative binding energy with −9.7 and −8.5 kcal/mol, and also exhibited electrostatic, hydrogen bonding, hydrophobic, and ionic bonding interactions with the amino acids of the receptors, as depicted in Fig. 16 and Fig. 17. Other molecules exhibited moderate to good binding affinity with respective receptors, and 2D and 3D images are given below Fig. 18 to Fig. 23.

**Fig. 16.**
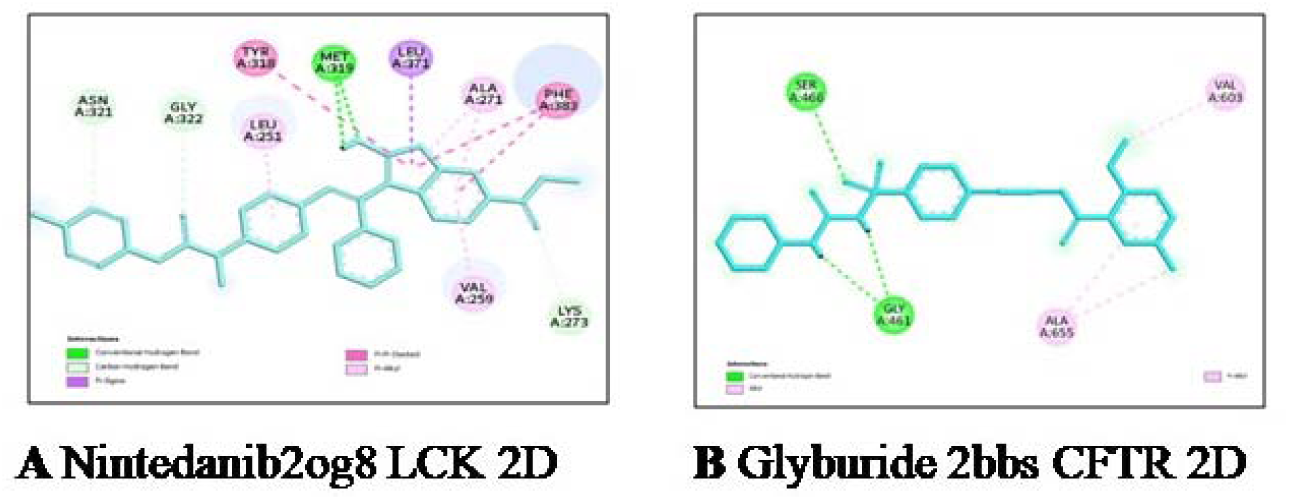
2D binding image of A) Ninatanib with LCK (pdb: 2og8) gene B) Glyburide with CFTR (pdb: 2bbs) gene.

**Fig. 17.**
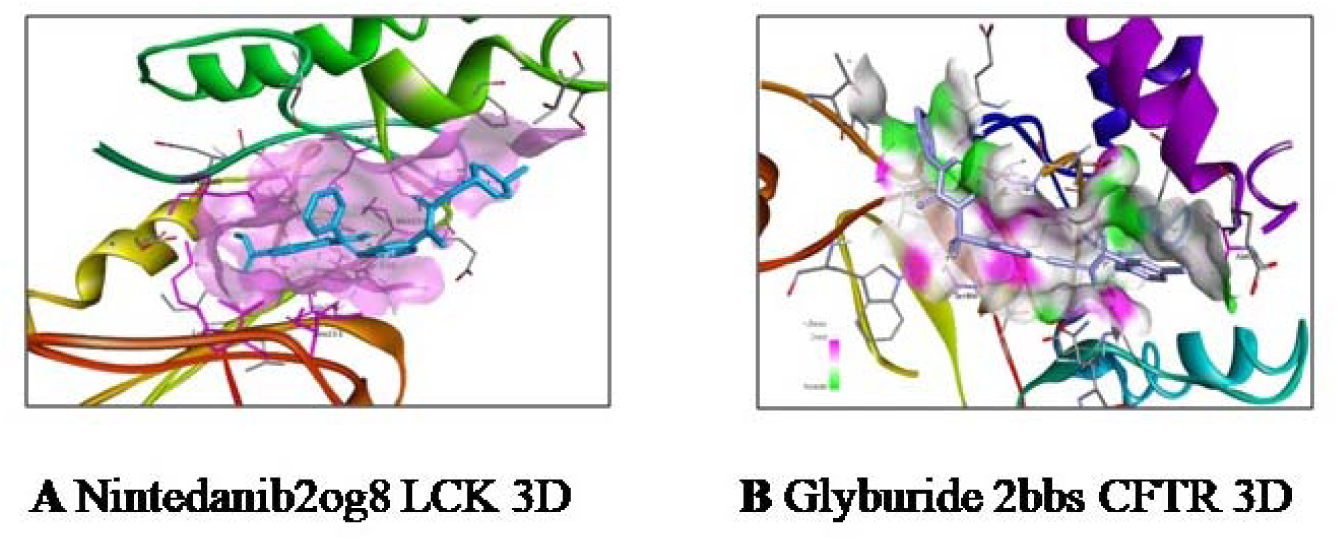
3D binding image of A) Ninatanib with LCK (pdb: 2og8) gene B) Glyburide with CFTR (pdb: 2bbs) gene.

**Fig. 18.**
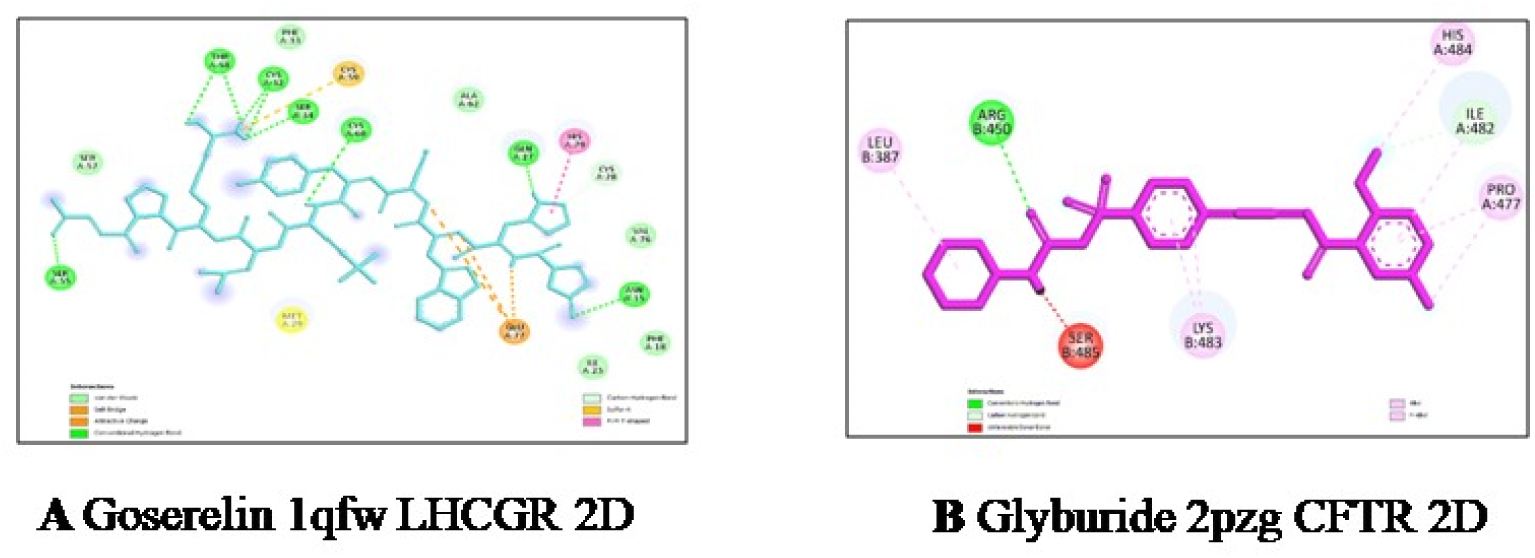
2D binding image of A) Goserelin with LHCGR (pdb: 1qfw) gene B) Glyburide with CFTR (pdb: 2pzg) gene.

**Fig. 19.**
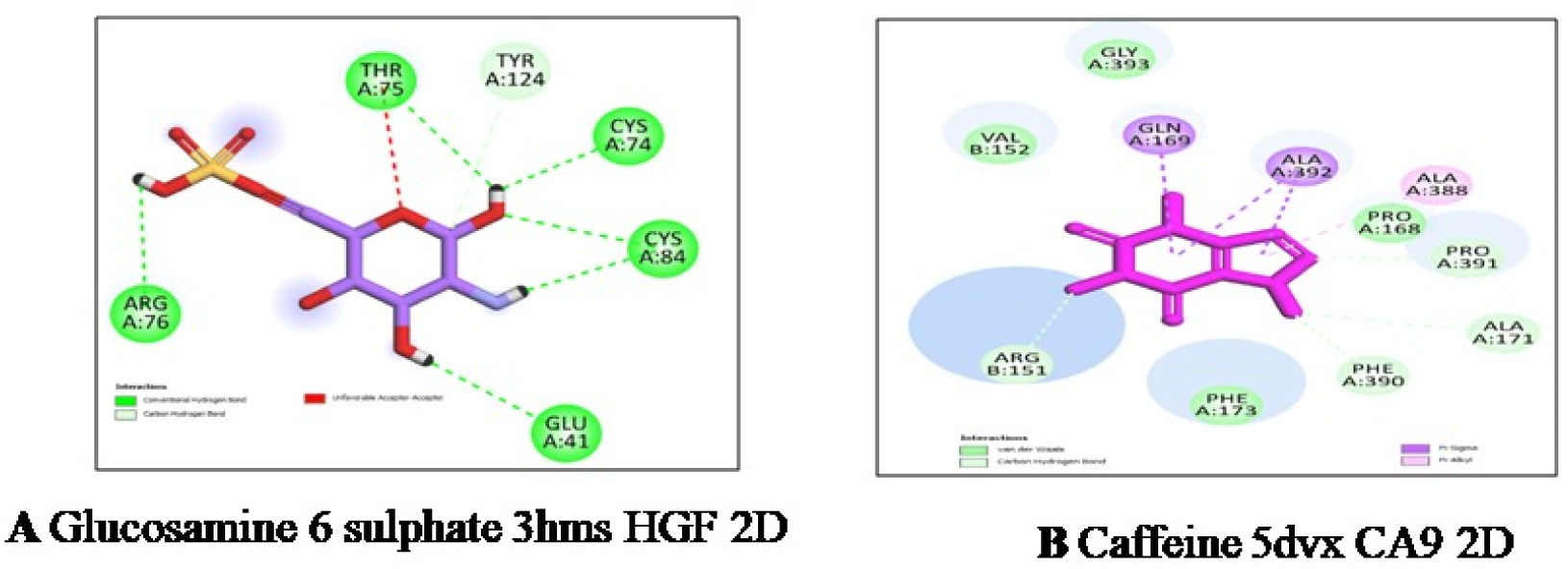
2D binding image of A) Glucosamine 6 sulphate with HGF (pdb: 3hms) gene B) Caffeine with CA9 (pdb: 5dvx) gene.

**Fig. 20.**
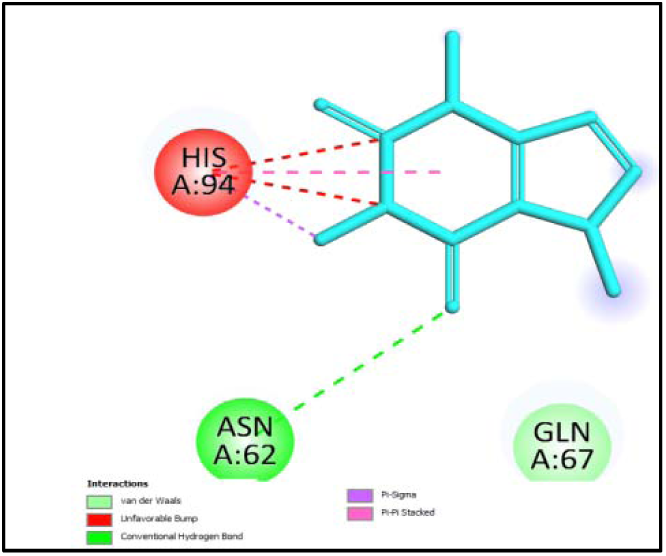
2D binding image of Caffeine with CA14 (pdb: 4lu3) gene

**Fig. 21.**
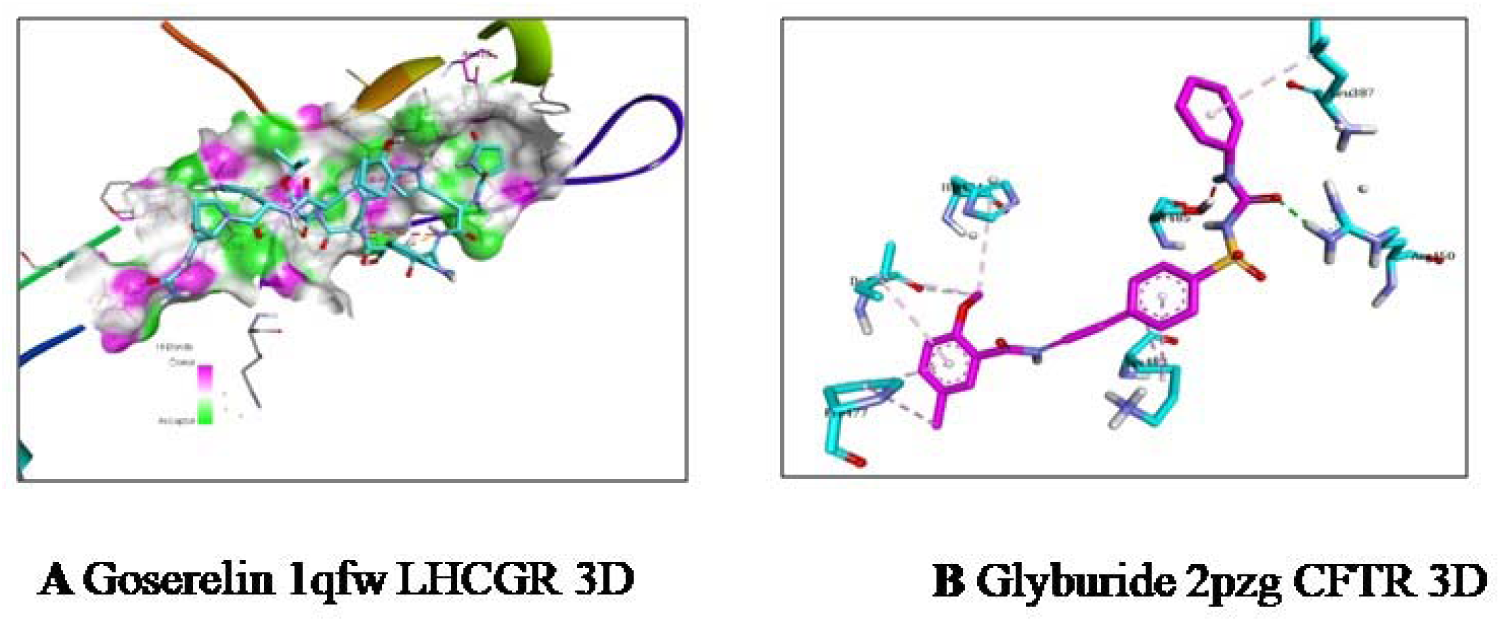
3D binding image of A) Goserelin with LHCGR (pdb: 1qfw) gene B) Glyburide with CFTR (pdb: 2pzg) gene.

**Fig. 22.**
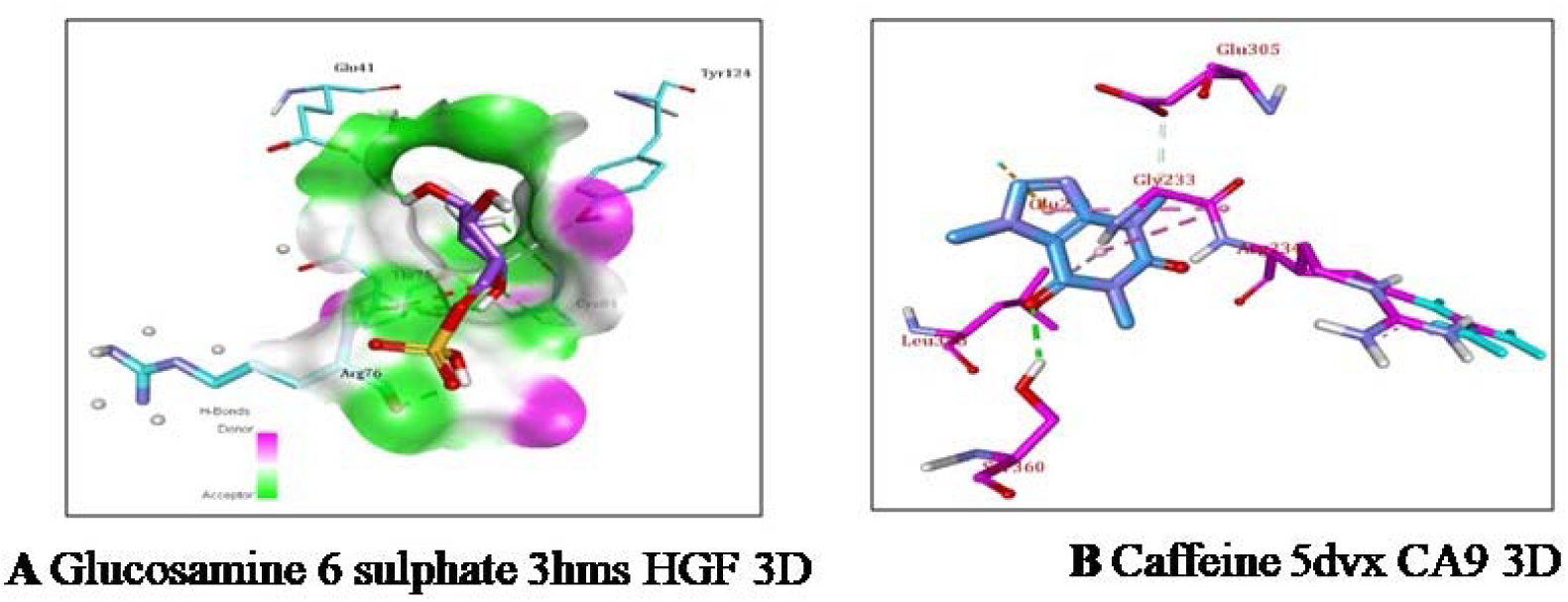
3D binding image of A) Glucosamine 6 sulphate with HGF (pdb: 3hms) gene B) Caffeine with CA9 (pdb: 5dvx) gene.

**Fig. 23.**
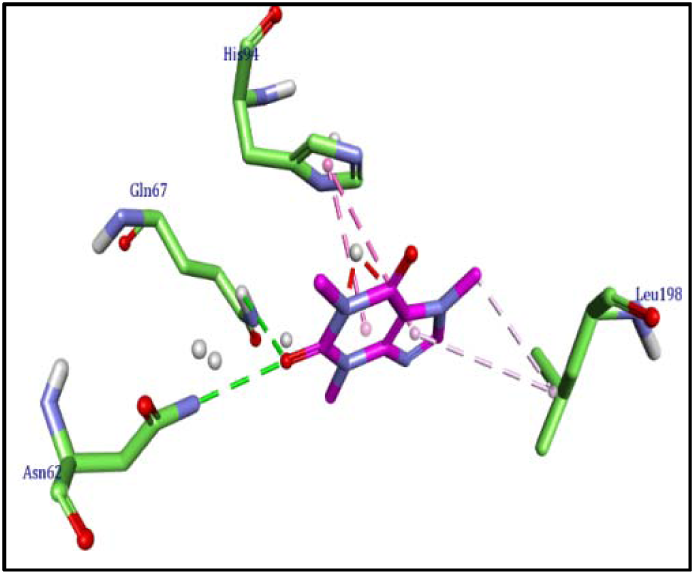
3D binding image of Caffeine with CA14 (pdb: 4lu3) gene

**Table 8.** Binding affinity and amino acid interactions of receptors and ligands.

| Compound and gene name (Receptor pdb id) | Affinity (kcal/mol) | Amino acid interactions |  |  |  |
| --- | --- | --- | --- | --- | --- |
|  |  | Hydrogen bonding interactions | Electrostatic interactions | Hydrophobic interactions | Ionic interactions |
| Nintedanib LCK (2og8) | -9.7 | MET A: 319, LYS A: 273, GLY A: 322, ASN A: 321 | -- | TYR A: 318, LEU A: 371, PHE A: 383, LEU A: 251, ALA A: 271, Val A: 259 | -- |
| Glyburide and CFTR (2pzg) | -8.5 | ARG A: 250, ILE A: 482 | SER 485 | PRO A: 477, HIS A: 484, LYS A: 483, LEU A: 387 | -- |
| Glyburide and CFTR (2bbs) | -7.5 | SER A: 466, GLY A: 461 | -- | ALA A: 551, VAL A: 603 | -- |
| Caffeine and CA9 (5dvx) | -6.7 | PHE A: 173, PHE A: 390, PRO A: 391, ALA A: 171, ARG A: 151 | -- | ALA A: 388, ALA A: 392, GLN A: 169, ARG A: 234, GLU A: 305, LEU A: 380 | -- |
| Goserelin and LHCGR (1qfw) | -6.6 | CYS A: 28, THR A: 58, CYS A: 32, SER A: 34, CYS A: 60, SER A: 55, GLN A: 27, ASN A: 15 | GLU A: 77 | CYS A: 59, HIS A: 79 | GLU A: 77 |
| Caffeine and CA14 (4lu3) | -5.6 | THR A: 200 | HIS A: 94, ASN A: 62 | LEU A: 198, VAL A: 121, ALA A: 143 | -- |
| Glucosamine 6-sulfate HGF (3hms) | -5.2 | TYR A: 124, THR A: 75, CYS A: 74, CYS A: 84, ARG A: 76, GLU A: 41 | -- | -- | -- |

## Discussion

MS is a high-incidence autoimmune inflammatory and non-traumatic disabling disease, often resulting in heterogeneous array of symptoms and signs because of differential involvement of motor, sensory, visual and autonomic systems [55]. However, in most cases, the clinical results are suboptimal. Therefore, understanding the pathological changes and molecular mechanisms of MS is essential for clinical diagnosis and treatment. Efficient bioinformatics analyses of NGS data help us understand the molecular mechanisms of the disease onset and advancement, which is invaluable for diagnostic and prognostic biomarkers, and therapeutic target identification [56–57]. In our study, 959 DEGs (479 up-regulated and 480 down-regulated genes) were identified between MS brain white matter samples and normal control brain white matter samples based on NGS dataset. Recent studies have shown that IGHV4-39 [58] play important roles in MS. Previous studies have demonstrated that PLAC8 [59] was shown to be involved in the obesity. Studies have shown that PLAC8 [59] might contribute to the pathogenesis of diabetes mellitus and could be a potential biomarkers for predicting disease progression and therapeutic effects in diabetes mellitus patients.. These findings might contribute to a better understanding of the molecular mechanisms underlying the pathogenesis of MS and its associated complications.

GO and REACTOME pathway enrichment analyses were used to explore the molecular mechanisms of the genes involved in the occurrence and development of MS and its associated complications. It is well known that the signaling pathways include immune system [60], TCR signaling [61], cytokine signaling in immune system [62], degradation of the extracellular matrix [63], extracellular matrix organization [63], metabolism of lipids [64] and metabolism [65] plays an important role only in MS. Recent studies have found that genes include CCL18 [66], FCRL3 [67], EOMES (eomesodermin) [68], CCR2 [69], IL2RB [70], CCL4 [71], FASLG (Fas ligand) [72], CD24 [73], IKZF3 [74], CD2 [75], CD28 [76], IL7R [77], HLA-DRB5 [78], ICOS (inducible T cell costimulator) [79], CCL5 [80], CTLA4 [81], IRF4 [82], C6 [83], NCR1 [84], CHIT1 [85], CD52 [86], CD163 [87], HGF (hepatocyte growth factor) [88], DIO3 [89], SIGLEC1 [90], TTR (transthyretin) [91], IL9 [92], VEGFA (vascular endothelial growth factor A) [93], CR2 [94], ANGPTL4 [95], CHI3L1 [96], MAG (myelin associated glycoprotein) [97], CNP (2’,3’-cyclic nucleotide 3’ phosphodiesterase) [98], CMTM5 [99], SEMA4D [100], NGFR (nerve growth factor receptor) [101], TF (transferrin) [102], MYRF (myelin regulatory factor) [103], MOG (myelin oligodendrocyte glycoprotein) [104], ADAMTS4 [105], BMP2 [106], HTRA1 [107], PNPLA3 [108], DYSF (dysferlin) [109], NINJ2 [110], LRP2 [111], ADAMTS14 [112] and DHCR7 [113] participates in the pathophysiological progression of MS. Recent evidence indicates that genes include CCL18 [114], SLAMF7 [115], GPR174 [116], CCR4 [117], POU4F2 [118]. CCR2 [119], IL2RB [120], CCL4 [121], CCL24 [122], FASLG (Fas ligand) [123], CD24 [124], TDGF1 [125], CD28 [126], IL7R [127], CYP11B1 [128], CCL5 [129], CCL3 [130], LTF (lactotransferrin) [131], GPNMB (glycoprotein nmb) [132], CD209 [133], IL2RG [134], CHIT1 [135], TAB2 [136], CD163 [137], ALOX15B [138], NMRK2 [139], HGF (hepatocyte growth factor) [140], TRPM8 [141], DIO3 [142], SIGLEC1 [143], TTR (transthyretin) [144], IL24 [145], F13A1 [146], IL9 [147], VEGFA (vascular endothelial growth factor A) [148], RASAL1 [149], ADM (adrenomedullin) [150], ANGPTL4 [151], CHI3L1 [152], LDB3 [153], CNP (2’,3’-cyclic nucleotide 3’ phosphodiesterase) [154], HES6 [155], CMTM5 [156], PLXNB3 [157], KLK8 [158], CDKN1C [159], INSIG1 [160], GREM1 [161], ATF3 [162], HK2 [163], MCAM (melanoma cell adhesion molecule) [164], SEMA4D [165], GLUL (glutamate-ammonia ligase) [166], S1PR5 [167], FN3K [168], MEIS1 [169], ADAMTS4 [170], BIN1 [171], BMP2 [172], LMNA (lamin A/C) [173], ERBB3 [174], DLL1 [175], THBS2 [176], GADD45B [177], MYH6 [178]. PNPLA3 [179], ACTN2 [180], MMP15 [181], SVEP1 [182], CPB2 [183], DYSF (dysferlin) [184], ADAMTSL2 [185], NINJ2 [186], LRP2 [111], PHLDA3 [187], LIPC (lipase C, hepatic type) [188], CARNS1 [189], PRODH (proline dehydrogenase 1) [190], TRPV6 [191], CNDP1 [192], DHCR7 [193], KCNE5 [194] SAA1 [195], ADSS1 [196], ACADS (acyl-CoA dehydrogenase short chain) [197], ADAP1 [198], KLK10 [199], HCN2 [200], F11 [201], RNASE1 [202], GNLY (granulysin) [203], EVA1A [204], CIRBP (cold inducible RNA binding protein) [205], EDIL3 [206], NOXA1 [207], ANGPTL2 [208], CYP2A6 [209] and CTNNA3 [210] might be implicated in the cardiovascular diseases progression. Gene include CCL18 [211], EOMES (eomesodermin) [212], GZMA (granzyme A) [213], HLA-DRB5 [214], GPNMB (glycoprotein nmb) [215], PRPH (peripherin) [216], HGF (hepatocyte growth factor) [217], TTR (transthyretin) [218], VEGFA (vascular endothelial growth factor A) [219], CHI3L1 [220], AATK (apoptosis associated tyrosine kinase) [221], SREBF1 [222], OLIG2 [223], ATF3 [224], NGFR (nerve growth factor receptor) [225], ADAMTS4 [226], LMNA (lamin A/C) [227], MT3 [228], DAO (D-amino acid oxidase) [229] and LGALS3BP [230] might be related to the progression of amyotrophic lateral sclerosis. SLAMF7 [231], GPR174 [232], FCRL3 [233], SLAMF6 [234], EOMES (eomesodermin) [235], CCR4 [236], CCR2 [237], CD48 [238], CD3E [239], CCL26 [240], CCL4 [241], SLAMF1 [242], CD24 [243], IKZF3 [244], CD2 [245], CD28 [246], IL7R [247], FCGR2B [248], UBASH3A [249], CD1C [250], ICOS (inducible T cell costimulator) [251], CD3G [252], CCL3 [253], LTF (lactotransferrin) [254], GPNMB (glycoprotein nmb) [255], CTLA4 [256], IRF4 [257], TNFRSF9 [258], FCRL5 [259], LAIR2 [260], TAB2 [261], CD52 [232], CD163 [263], MSR1 [264], HGF (hepatocyte growth factor) [265], SIGLEC1 [266], TTR (transthyretin) [267], IL24 [268], IL9 [269], CHI3L1 [270], CDH1 [271], OLIG2 [272], NKX6-2 [273]. HK2 [274], PNPLA3 [275], CPB2 [276], SEMA4C [277], LRP2 [278], SLC5A11 [279], F11 [280], ANGPTL2 [281] and CYP2A6 [282] might serve as molecular markers for autoimmune disease. Recent studies have found that genes include SLAMF7 [283], FCRL3 [284], CCR4 [285], GZMA (granzyme A) [286], CCR2 [287], CD48 [288], CD3E [289], CCL4 [290], FASLG (Fas ligand) [291], SLAMF1 [292], CD28 [293], IL7R [294], UBASH3A [295], CD3D [296], ICOS (inducible T cell costimulator) [297], CCL5 [298], CCL3 [299], LTF (lactotransferrin) [300], CTLA4 [301], TNFRSF9 [258], C6 [302], IL5RA [303], IL2RG [304], NPY2R [305], DRD3 [306], CD52 [307], CD163 [308], IQGAP2 [309], PRPH (peripherin) [310], MSR1 [311], HGF (hepatocyte growth factor) [312], TTR (transthyretin) [313], IL9 [314], ADM (adrenomedullin) [315], ANGPTL4 [316], CHI3L1 [317], CDH1 [318], SREBF1 [319], CDKN1C [320], TSC22D4 [321], CFTR (CF transmembrane conductance regulator) [322], INSIG1 [323], HSD17B3 [324], GREM1 [325], ATF3 [326], HK2 [327], TF (transferrin) [328], GLUL (glutamate-ammonia ligase) [329], DMRT2 [330], FN3K [331], TREH (trehalase) [332], ADAMTS4 [333], BMP2 [334], LMNA (lamin A/C) [335], ERBB3 [336], HSD11B1 [337], DLL1 [338], NKX2-2 [339], FGF11 [340], ASPA (aspartoacylase) [341], FASN (fatty acid synthase) [342], PNPLA3 [343], ADORA1 [344], WNT7A [345], LIPC (lipase C, hepatic type) [346], GPR142 [347], GPD1 [348], LIPE (lipase E, hormone sensitive type) [349], CNDP1 [192], DHCR7 [350], SAA1 [351], TKTL1 [352], ACADS (acyl-CoA dehydrogenase short chain) [353], SCD (stearoyl-CoA desaturase) [354], FFAR1 [355], HCN2 [200], F11 [356], GPIHBP1 [357], ANGPTL2 [208], MAPK8IP1 [358], CYP2A6 [359] and CTNNA3 [360] have been reported its expression in the diabetes mellitus. Studies have revealed that genes include MS4A2 [361], CD48 [362], HLA-DRB5 [363], FCGR2B [364], CCL5 [365], CCL3 [366], LTF (lactotransferrin) [367], GPNMB (glycoprotein nmb) [368], CTLA4 [369], TRIML2 [370], PTGDR (prostaglandin D2 receptor) [371], DRD3 [372], LHCGR (luteinizing hormone/choriogonadotropin receptor) [373], CD163 [374], PRPH (peripherin) [216], MSR1 [375], HGF (hepatocyte growth factor) [376], TTR (transthyretin) [377], IL9 [378], ADM (adrenomedullin) [379], CHI3L1 [380], CNP (2’,3’-cyclic nucleotide 3’ phosphodiesterase) [381], CDH1 [382], OLIG2 [383], KLK8 [384]. CFTR (CF transmembrane conductance regulator) [385], ABCA2 [386], HK2 [387], NGFR (nerve growth factor receptor) [388], TF (transferrin) [389], GLUL (glutamate-ammonia ligase) [329], ADAMTS4 [390], BIN1 [391], HSPA2 [392], LMNA (lamin A/C) [393], HSD11B1 [394], HTRA1 [395], APLP1 [396], MT3 [397], ADORA1 [398], DYSF (dysferlin) [399], SLC24A4 [400], RHOB (ras homolog family member B) [401], NINJ2 [402], LRP2 [403], LIPC (lipase C, hepatic type) [404], DHCR7 [405], SCD (stearoyl-CoA desaturase) [406], F11 [407], PFKL (phosphofructokinase, liver type) [408], ALDH1A1 [409], SPON1 [410] and CTNNA3 [411] plays a role in diagnosis of Alzheimer’s Disease. The abnormal activation of genes include CCR4 [412], CCR2 [413], CD48 [414], CD28 [415], CCL3 [416], CTLA4 [417], C6 [418], MICB (MHC class I polypeptide-related sequence B) [419], DRD3 [420], HGF (hepatocyte growth factor) [421], DIO3 [422], TTR (transthyretin) [423], GRHL3 [434], VEGFA (vascular endothelial growth factor A) [425], ADM (adrenomedullin) [426], CHI3L1 [427], SOX3 [428], MAG (myelin associated glycoprotein) [429], CNP (2’,3’-cyclic nucleotide 3’ phosphodiesterase) [430], SREBF1 [431], OLIG2 [432], ATF3 [433], NGFR (nerve growth factor receptor) [434], TF (transferrin) [435], GLUL (glutamate-ammonia ligase) [329]. MOG (myelin oligodendrocyte glycoprotein) [436], ERBB3 [437], NHLH2 [438], GADD45B [439], PADI2 [440], ADORA1 [441], NINJ2 [442], WNT7A [443], DAO (D-amino acid oxidase) [444], PRODH (proline dehydrogenase 1) [445] and HCRTR1 [446] in psychiatric disorders and are a significant cause of disease progression. Some researchers have reported that altered CCR2 [447], FASLG (Fas ligand) [448], GPNMB (glycoprotein nmb) [449], NPY2R [450], TRPM8 [451], TTR (transthyretin) [452], ADM (adrenomedullin) [453], CHI3L1 [380], CDH1 [382], HIP1R [454], SEMA4D [455], HAPLN2 [456], MYRF (myelin regulatory factor) [457], BIN1 [458], BMP2 [459], APLP1 [396], SEPTIN4 [460], DAO (D-amino acid oxidase) [461], TKTL1 [352], ADAP1 [462], HPDL (4-hydroxyphenylpyruvate dioxygenase like) [463], NME3 [464], SPON1 [465] and LGALS3BP [230] expression in the neurodegenerative diseases. Some studies have shown that genes include CCR2 [466], FASLG (Fas ligand) [467], CD28 [468], LYZ (lysozyme) [469], CCL5 [470], CCL3 [471], C6 [472], MSR1 [473], HGF (hepatocyte growth factor) [474], VEGFA (vascular endothelial growth factor A) [475], ADM (adrenomedullin) [476], CR2 [477], MAG (myelin associated glycoprotein) [478], CDH1 [479], OLIG2 [480], SEMA4D [481], ADAMTS4 [482], BMP2 [483], ERBB3 [484] and NKX2-2 [485] plays a certain role in spinal cord injury. The altered expression of genes include CCR2 [486], FASLG (Fas ligand) [291], CD24 [487], CD28 [488], LYZ (lysozyme) [489], IL7R [490], CCL5 [491], LTF (lactotransferrin) [492], GPNMB (glycoprotein nmb) [493], CTLA4 [494], IRF4 [495], CHIT1 [496], NPY2R [497], TAB2 [136], CD52 [307], CD163 [498], MSR1 [311], HGF (hepatocyte growth factor) [499], TRPM8 [500], TTR (transthyretin) [501], F13A1 [502], ADM (adrenomedullin) [503], COL9A3 [504], ANGPTL4 [505], CHI3L1 [506], BMP8A [507], SREBF1 [508], CDKN1C [509], INSIG1 [510], ATF3 [511], HK2 [512], TF (transferrin) [513], MOG (myelin oligodendrocyte glycoprotein) [514], ADAMTS4 [515], BMP2 [516], HSPA2 [517], LMNA (lamin A/C) [335], HSD11B1 [518], NKX2-2 [519], NHLH2 [438], FGF11 [520], ASPA (aspartoacylase) [341], FASN (fatty acid synthase) [521], PNPLA3 [522], MMP15 [523], CBR1 [524], LRP2 [525], WNT7A [345], GPR142 [526], GPD1 [348], LIPE (lipase E, hormone sensitive type) [527], CNDP1 [528], SAA1 [529], ACADS (acyl-CoA dehydrogenase short chain) [353], SCD (stearoyl-CoA desaturase) [530], GPIHBP [531], LMF1 [532], IGSF1 [533], LSS (lanosterol synthase) [534], ALDH1A1 [535], ANGPTL2 [536], NENF (neudesin neurotrophic factor) [537] and CYP2A6 [538] promotes occurrence and progression of obesity. A previous study reported that the genes include CCR2 [539], CCL4 [540], TNFRSF13B [541], FASLG (Fas ligand) [542], CD28 [543], CYP11B1 [544], CCL5 [545], LTF (lactotransferrin) [546], CTLA4 [547], DRD3 [548], HGF (hepatocyte growth factor) [499], TRPM8 [549], IL9 [550], NPPC (natriuretic peptide C) [551], ANGPTL4 [552], CHI3L1 [553], SREBF1 [554], GDF2 [555], CFTR (CF transmembrane conductance regulator) [556], ATF3 [557], HK2 [558], NGFR (nerve growth factor receptor) [559], MEIS1 [560], MOG (myelin oligodendrocyte glycoprotein) [561], BMP2 [562], LMNA (lamin A/C) [563], ERBB3 [336], HSD11B1 [564], THBS2 [565], FASN (fatty acid synthase) [566], PNPLA3 [567], SLC45A3 [568], WNT7A [569], SCNN1G [570], TRPV5 [571], ACADS (acyl-CoA dehydrogenase short chain) [572], GNLY (granulysin) [573], NOXA1 [574], ANGPTL2 [575], SPON1 [576], CYP2A6 [577] and RASA3 [578] were associated with hypertension. Altered expression of genes include FASLG (Fas ligand) [579], GJC2 [580] and GJB1 [581] have been demonstrated to be regulated in spasticity. Genes include CD24 [582], CD28 [583], HLA-DRB5 [584], LTF (lactotransferrin) [585], GPNMB (glycoprotein nmb) [586], TNFRSF9 [587], LRRC37A2 [588], DRD3 [589], CD163 [374], HGF (hepatocyte growth factor) [590], TRPM8 [451], TTR (transthyretin) [591], VEGFA (vascular endothelial growth factor A) [592], CHI3L1 [593], MAG (myelin associated glycoprotein) [594], SREBF1 [222], HIP1R [595], HK2 [596], GPR37 [597], NGFR (nerve growth factor receptor) [598], TF (transferrin) [599], HAPLN2 [456], MOG (myelin oligodendrocyte glycoprotein) [600], BIN1 [601], BMP2 [602], GADD45B [603], UNC5B [604], ADORA1 [605], SEPTIN4 [606], DHCR7 [607], SCD (stearoyl-CoA desaturase) [608], GIPC1 [609], ALDH1A1 [610] and CTNNA3 [611] have been shown to influence the genetic risk of Parkinson’s disease. However, further investigations are needed to explore and confirm the potentially significant GO terms and signaling pathways for MS and to achieve a comprehensive understanding of this process.

Based on the PPI network constructed and modules analyzed by the online PPI database HiPPIE, we identified hub genes. Previous studies have proven that the hub genes include SLAMF1 [242], TAB2 [261] and HGF (hepatocyte growth factor) [265] as a genetic locus associated with autoimmune disease susceptibility. Recent studies have shown that hub genes include SLAMF1 [292], CFTR (CF transmembrane conductance regulator) [322], LMNA (lamin A/C) [335], ERBB3 [336], HGF (hepatocyte growth factor) [312] and CCL5 [298] are genetic risk factors have been shown to increase susceptibility to diabetes mellitus. Hub genes include TAB2 [136], LMNA (lamin A/C) [173], ERBB3 [174], HGF (hepatocyte growth factor) [140] and CCL5 [129] have been reported its expression in the cardiovascular diseases. Altered levels of hub genes include TAB2 [136], LMNA (lamin A/C) [335], HGF (hepatocyte growth factor) [499] and CCL5 [491] have been detected in the obesity. Hub genes include CFTR (CF transmembrane conductance regulator) [385], RHOB (ras homolog family member B) [401], LMNA (lamin A/C) [393], HGF (hepatocyte growth factor) [376], CCL5 [365] and NGFR (nerve growth factor receptor) [388] expression was shown to be regulated in a Alzheimer’s Disease. Altered level of hub genes include CFTR (CF transmembrane conductance regulator) [556], LMNA (lamin A/C) [563], ERBB3 [336], HGF (hepatocyte growth factor) [499], CCL5 [545] and NGFR (nerve growth factor receptor) [559] can facilitate hypertension progression. Hub Genes include LMNA (lamin A/C) [227], HGF (hepatocyte growth factor) [217] and NGFR (nerve growth factor receptor) [225] have been identified the involvement in amyotrophic lateral sclerosis. Hub genes include ERBB3 [437], HGF (hepatocyte growth factor) [421] and NGFR (nerve growth factor receptor) [434] could potentially play a role in the onset of psychiatric disorders. Hub genes include ERBB3 [484], HGF (hepatocyte growth factor) [474] and CCL5 [470] were found to be linked with spinal cord injury. Hub genes include HGF (hepatocyte growth factor) [88], CCL5 [80] and NGFR (nerve growth factor receptor) [101] were recently found to be a promoter of MS. Hub genes include HGF (hepatocyte growth factor) [590] and NGFR (nerve growth factor receptor) [598] plays a crucial role in establishing Parkinson’s disease. Our findings suggest that hub genes include LCK (LCK proto-oncogene, Src family tyrosine kinase), PYHIN1, DOK2, EGLN3, SDC1 and FSCN1 served as a novel biomarkers for MS. However, the exact molecular mechanism of MS progression remains unknown. This investigation might provide reference for research the connection between MS and its associated complication.

MiRNA-hub gene regulatory network analyses and TF-hub gene regulatory network are formed from the interactions of hub genes, miRNAs and TFs, and participate in all steps of the MS advancement. Recent research indicated that biomarkers include IKZF3 [74], hsa-mir-146a-5p [612], BRCA1 [613] and NFKB1 [614] might be considered as a pathogenic genetic factors for MS. Previous studies have shown that biomarkers include IKZF3 [244], TAB2 [261], CCR4 [236], LTF (lactotransferrin) [254], CDH1 [271], BRCA1 [615], NFKB1 [616] and FOXO3 [617] might have a significant role in the development of autoimmune disease. Several studies suggested that biomarkers include TAB2 [136], CCR4 [117], LTF (lactotransferrin) [131], hsa-mir-146a-5p [618], hsa-mir-8485 [619], BRCA1 [620], NFKB1 [621], TEAD1 [622], TFAP2C [623], FOXO3 [624] and NRF1 [625] might serve as therapeutic targets for cardiovascular diseases. A recent study showed that biomarkers include TAB2 [136], LTF (lactotransferrin) [492], BRCA1 [626], NFKB1 [627], TEAD1 [628], FOXO3 [629] and NRF1 [630] improves progression of obesity. The dysregulation of cellular processes that involve biomarkers include CCR4 [285], LTF (lactotransferrin) [300], CDH1 [318], CFTR (CF transmembrane conductance regulator) [322], hsa-mir-146a-5p [631], hsa-mir-195-5p [632], hsa-mir-4739 [633], HNF4A [634], NFKB1 [635], NR2F1 [636], FOXO3 [637] and NRF1 [638] have been linked to the development of diabetes mellitus. A previous study reported that the biomarkers include CCR4 [412], hsa-mir-146a-5p [639] and NFKB1 [640] were associated with psychiatric disorders. LTF (lactotransferrin) [367], RHOB (ras homolog family member B) [401], CDH1 [382], CFTR (CF transmembrane conductance regulator) [385], hsa-mir-146a-5p [641], hsa-mir-195-5p [642], BRCA1 [643], ELK4 [644] and FOXO3 [645] were associated with Alzheimer’s disease. Biomarkers include LTF (lactotransferrin) [546], CFTR (CF transmembrane conductance regulator) [556], hsa-mir-195-5p [632], BRCA1 [646], NFKB1 [647], TFAP2C [648], FOXO3 [649] and NRF1 [650] might be a potential therapeutic targets for hypertension. In many clinical studies, biomarkers include LTF (lactotransferrin) [585], hsa-mir-146a-5p [651], hsa-mir-195-5p [642], HNF4A [652], NFKB1 [643], NR2F1 [654], FOXO3 [655 and NRF1 [656] were alerted expression in Parkinson’s disease. Findings have shown that biomarkers include LGALS3BP [230] and BRCA1 [657] expression is regulated in amyotrophic lateral sclerosis. Studies have shown that biomarkers include LGALS3BP [230], CDH1 [382], hsa-mir-146a-5p [658], BRCA1 [659], HNF4A [660], FOXO3 [661] and NRF1 [662] plays an important role in the development of neurodegenerative diseases. A previous study found that biomarkers include CDH1 [479] and FOXO3 [663] have been linked to the development of spinal cord injury. Hence, the results of these regulatory networks analysis were in accordance with experimental results on MS. The roles of novel biomarkers in the these networks were determined and still not reported in MS and its associated complications, such as C6orf141, SDC1, CD79A, LCK (LCK proto-oncogene, Src family tyrosine kinase), SFN (stratifin), POLR2F, MAP1LC3A, LLGL2, FOXE1, hsa-mir-6794-3p, hsa-mir-3689a-3p, hsa-mir-4449, hsa-mir- 4695-5p, hsa-mir-4651, hsa-mir-548q and NFYA (nuclear transcription factor Y subunit alpha). The intricate interaction between MSs, TFs and hub genes made great improvement to the advancement of MS.

All the screened molecules were very well bound with respective protein molecules, and some of them showed potential binding affinity Nintedanib to LCK and Glyburide to CFTR genes, Nintedanib and Glyburide might be the potential lead molecule to bring breakthrough in the field of medicine in future. All the data needs further research on these two molecules in the future to combat the burden of MS on humankind.

Nevertheless, our study has some limitations. Firstly, due to the fact that the data we used were obtained from brain white matter, derived from of MS patients are desired for clinical application. Secondly, although we have validated the hub gene in ROC analysis, the validity of the hub genes, miRNAs and TFs for MS diagnosis must be confirmed in in-vitro cell culture experiments and in-vivo animal or human experiments as possible in the future. In addition, due to the lack of comprehensive clinical data, we were inadequate to check the relationship between patients with distinct clinical symptoms and complications, and hub genes, miRNAs and TFs. Future investigation could expand the sample size and execute comprehensive phenotyping of patients, to superior understand the potential molecular mechanisms underlying MS and its associated complications. We hope that future investigation can overcome these limitations and progresses deeper into the molecular pathogenesis of MS and its associated complications, providing more scientific confirmation for the growth of better treatment options.

## Conclusions

To summarize, a total of 959 DEGs, hub genes (LCK, PYHIN1, SLAMF1, DOK2, TAB2, CFTR, RHOB, LMNA, EGLN3 and ERBB3), miRNAs (hsa-mir-6794-3p, hsa-mir-3689a-3p, hsa-mir-4651and hsa-mir-548q) and TFs (BRCA1, HNF4A, TFAP2C and NR2F1) were identified, which could be considered as MS biomarkers. However, further investigations are needed to clarify the biological roles of these genes in MS.

## Abbreviations

MS: Multiple sclerosis
DEGs: Differentially expressed genes
NGS: Next generation sequencing
GEO: Gene expression omnibus
GO: Gene 0ntology
PPI: Protein-protein interaction
miRNA: Micro ribonuclic acid
TF: Transcription factor
ROC: Receiver operating characteristic curve
LCK: LCK proto-oncogene, Src family tyrosine kinase
PYHIN1: Pyrin and HIN domain family member 1
SLAMF1: Signaling lymphocytic activation molecule family member 1
DOK2: Docking protein 2
TAB2: TGF-beta activated kinase 1 (MAP3K7) binding protein 2
CFTR: CF transmembrane conductance regulator
RHOB: Ras homolog family member B
LMNA: Lamin A/C
EGLN3: Egl-9 family hypoxia inducible factor 3
ERBB3: Erb-b2 receptor tyrosine kinase 3

## Acknowledgement

I Tobias Frisch, University of Southern Denmark, Campusvej 55, Odense M, Denmark, very much, the author who deposited their NGS dataset GSE138614, into the public GEO database.

## Ethical approval

This article does not contain any studies with human participants or animals performed by any of the authors.

## Informed consent

No informed consent because this study does not contain human or animals participants.

## Availability of data and materials

The datasets supporting the conclusions of this article are available in the GEO (Gene Expression Omnibus) (https://www.ncbi.nlm.nih.gov/geo/) repository. [(GSE138614) https://www.ncbi.nlm.nih.gov/geo/query/acc.cgi?acc=GSE138614]

## Consent for publication

Not applicable.

## Author Contributions

B. V. - Writing original draft, and review and editing

S.P. - Formal analysis and validation

C. V. - Software and investigation

